# CGAS (Chloroplast Genome Analysis Suite): An Automated Python Pipeline for Comprehensive Comparative Chloroplast Genomics

**DOI:** 10.64898/2025.12.21.695765

**Authors:** Abdullah, Rushan Yan, Xiaoxuan Tian

## Abstract

**Background:** Chloroplast genome analysis underpins plant phylogenetics, comparative genomics, and molecular marker development. Although high-quality chloroplast genomes are now routinely generated, downstream comparative analyses still rely on fragmented toolchains requiring separate tools for annotation, codon analysis, and SNP detection, extensive manual curation, and ad hoc scripting, which limit reproducibility and scalability.

**Results:** We present the Chloroplast Genome Analysis Suite (CGAS), an automated Python-based pipeline designed as a comprehensive solution for comparative chloroplast genomics. CGAS integrates nine core analytical modules into a single, reproducible workflow operating directly on annotated GenBank files and pre-aligned FASTA datasets. The pipeline performs automated gene content analysis, publication-ready gene table generation, chloroplast structural characterization (LSC/SSC/IR boundaries and GC content), codon usage and amino acid composition analyses enabling batch processing of 10-50+ genomes, SNP detection with transition/transversion statistics, intron feature characterization, simple sequence repeat identification, and nucleotide diversity profiling. Unlike existing tools, CGAS emphasizes biologically informed handling of edge cases, including trans-spliced genes (e.g., *rps12*), IR-mediated gene duplication, and annotation artifacts. All outputs are generated in standardized, publication-ready Excel and Word formats with timestamped provenance.

**Conclusions:** CGAS provides a unified, automated, and biologically robust framework for comparative chloroplast genome analysis. By minimizing manual intervention and emphasizing batch processing, the suite substantially accelerates methodologically consistent analyses. The pipeline is openly available at https://github.com/abdullah30/Chloroplast-Genome-Analysis-Suite-CGAS and readily extensible, supporting its adoption as a standardized analytical backend for chloroplast comparative genomics.

## Introduction

Chloroplast genomes (cp genomes) are central to studies of plant evolution, phylogeny, and molecular identification due to their conserved structure, uniparental inheritance, and moderate evolutionary rate (Abdullah et al., 2025; Daniell et al., 2016). Typical angiosperm chloroplast genomes exhibit a quadripartite organization consisting of a large single-copy (LSC) region, a small single-copy (SSC) region, and two inverted repeats (IRa and IRb), typically spanning 120-160 kb with 110-130 genes (Daniell et al., 2016; Palmer, 1985).

Advances in next-generation sequencing have dramatically increased the availability of complete cp genomes, shifting the analytical bottleneck from genome generation to downstream comparative analysis.

Despite this data abundance, comprehensive chloroplast genome analyses remain methodologically fragmented. Researchers commonly rely on multiple independent tools to extract gene inventories, assess IR boundaries and junction genes, calculate codon usage, simple sequence repeats, long repeats, identify SNPs, and characterize intron structures (Amiryousefi et al., 2018; Beier et al., 2017; Kearse et al., 2012; Kumar et al., 2018; Kurtz et al., 2001; Li et al., 2023; Tillich et al., 2017). These steps often require manual file conversion, spreadsheet compilation, and repeated parameter tuning across different software packages, introducing inconsistency and limiting reproducibility—particularly in large comparative datasets.

Recent efforts have been made to develop integrated workflows. CPStools (Huang et al., 2024) provides valuable resources for relative synonymous codon usage calculations, nucleotide diversity, and simple sequence repeats analysis. CPGview (Liu et al., 2023) effectively visualizes gene distributions, variable sites, repetitive sequences, and trans-spliced gene structures such as *rps12*. However, several analytical gaps remain: CPStools lacks amino acid frequency analysis, substitution analysis, publication-ready gene content tables, intron characterization, and formatted outputs for genomic region lengths and GC content profiles. CPGview requires individual GenBank file uploads with manual consolidation of results, limiting its utility for large-scale batch processing. Moreover, to the best of our knowledge, no publicly available tools systematically account for biological complexities such as trans-spliced genes or annotation-derived pseudogene inflation while simultaneously performing comprehensive comparative analyses across multiple genomes and generating publication-ready outputs. To address these limitations, we developed the Chloroplast Genome Analysis Suite (CGAS) as a methods-focused, automation-first pipeline that emphasizes reproducibility, directory-based batch processing of multiple genomes, and automated generation of publication-ready outputs suitable for direct inclusion in manuscripts or supplementary materials.

### Overview of the Chloroplast Genome Analysis Suite

CGAS is implemented in Python (≥3.8) and organized as nine interoperable analytical modules. The suite can be executed as a unified pipeline or as individual modules, depending on analytical needs. Design priorities included: (1) Directory-based batch processing for comparative studies, (2) Biologically informed logic, particularly for trans-spliced and IR-duplicated genes, (3) Publication-ready outputs requiring no post-processing, and (4) Robust error handling to ensure continued execution in heterogeneous datasets.

The complete source code and documentation are hosted at: https://github.com/abdullah30/Chloroplast-Genome-Analysis-Suite-CGAS

## Methods

### Software Architecture and Dependencies

CGAS is written in Python and relies on widely adopted, actively maintained libraries. Core dependencies include Biopython (≥1.79) for sequence parsing, Pandas (≥1.3) for tabular data handling, OpenPyXL (≥3.0) for Excel output formatting, and python-docx (≥0.8.11) for automated Word document generation. The modular architecture enables flexible execution modes, including full-pipeline runs, selective module execution, and function-level integration into custom analytical workflows or Jupyter notebooks. For Jupyter Notebook, the individual scripts along with the unified analyzer script are provided as **Supplementary Data File 1**.

### Execution Modes and Cross-Platform Compatibility

CGAS supports multiple execution modes to accommodate diverse computational environments and user expertise levels (**Figure 1**):

**Figure 1.**
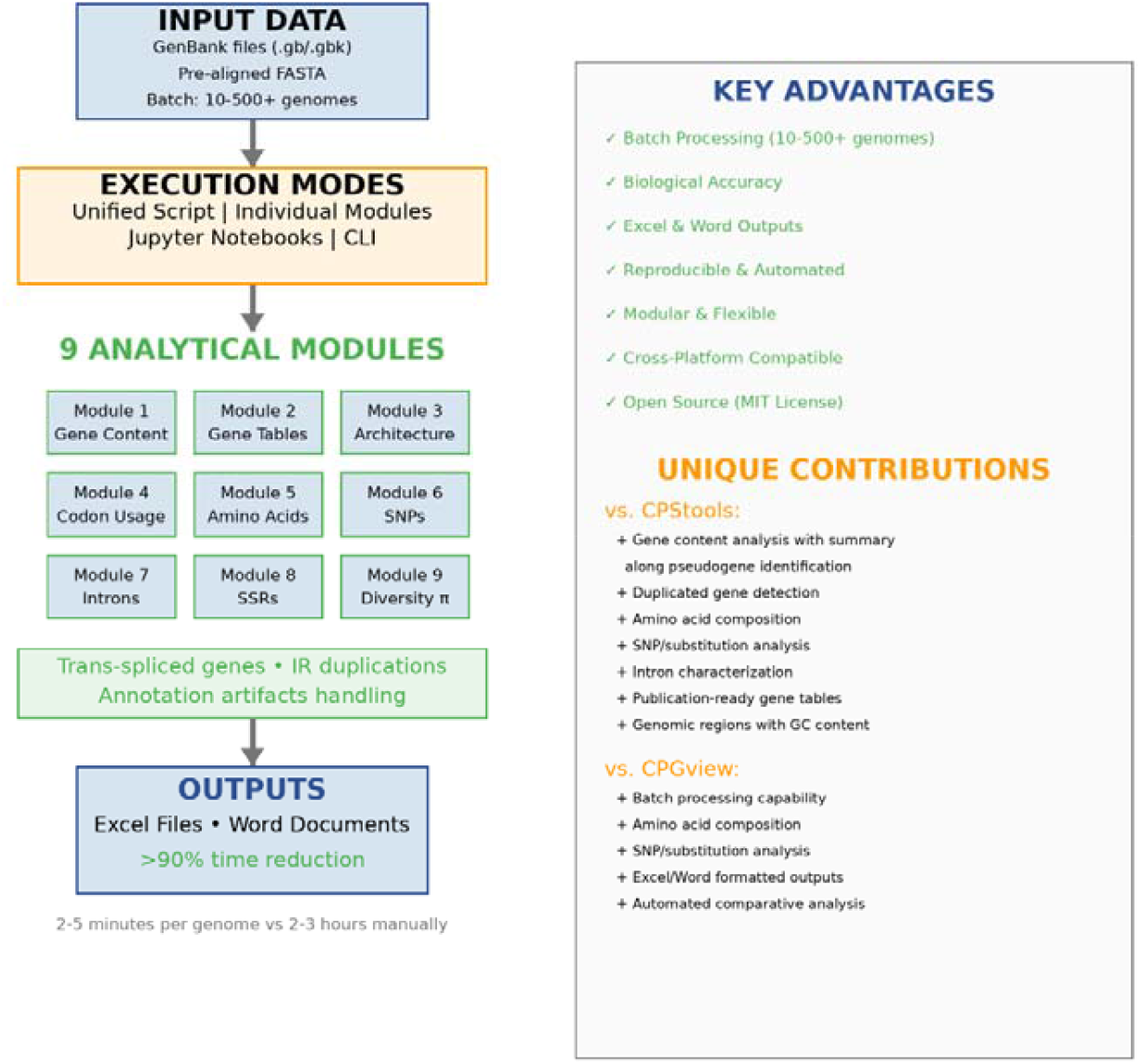
CGAS workflow architecture showing input data types, execution modes, nine analytical modules, and publication-ready outputs. The pipeline accepts GenBank files and pre-aligned FASTA sequences, processes them through nine integrated modules with biologically informed logic for trans-spliced genes and IR duplications, and generates timestamped Excel and Word outputs suitable for direct publication.

#### Unified Script Mode

Execute all modules sequentially with a single command (`python chloroplast_unified_analyzer.py`), ideal for comprehensive batch processing. We recommend this mode for ease and complete results, as the script does not stop if a module fails—it displays a message and continues with remaining modules.

#### Individual Script Mode

Run standalone Python scripts for specific analyses, enabling targeted investigations without complete pipeline execution.

#### Jupyter Notebook Mode

Interactive execution within Jupyter notebooks, facilitating exploratory analysis, result visualization, and educational demonstrations.

#### Command-Line Interface

Parameterized execution via command-line arguments for selective module activation and scripted automation after installation from GitHub.

The suite is fully cross-platform compatible, operating seamlessly on Windows, macOS, and Linux systems. Linux server deployment is particularly well-supported for batch job scheduling and integration into high-performance computing workflows. This flexibility ensures CGAS can be deployed in environments ranging from individual researcher workstations to institutional computing clusters. The complete workflow architecture is summarized in **Figure 1**.

### Input Data Requirements

The pipeline accepts two primary data types:

1. Annotated chloroplast genomes in GenBank format (.gb/.gbk/.gbff)
2. Pre-aligned FASTA files for SNP analysis, generated using standard alignment software

GenBank files are assumed to be well-annotated and conform to standard feature conventions (CDS, tRNA, rRNA, gene). The example input data used for validation are provided as **Supplementary Data File 2**. CGAS does not perform de novo annotation; instead, it focuses on downstream comparative analyses. The example output is provided as **Supplementary Date File 3**.

#### Module 1: Gene Content, Count, and Summary Statistics

This module parses all gene-related features from GenBank files to generate comprehensive gene inventories. Functional genes and pseudogenes are distinguished using feature qualifiers and annotation notes. IR-associated duplications are detected by repeated gene names with different genomic coordinates (typically >10 kb apart for IR duplications).

To prevent biologically misleading inflation of gene counts, CGAS collapses IR duplicates into single functional units for summary statistics while retaining duplication information for structural interpretation. Importantly, CGAS preserves gene names exactly as annotated in input GenBank files without attempting automatic standardization. This design choice serves two purposes: first, it maintains compatibility with downstream analyses requiring uniform annotations (e.g., nucleotide diversity calculations), where automatic name correction could disrupt gene alignment; second, it alerts researchers to annotation inconsistencies across genomes, facilitating manual curation when needed. While minor variations in gene nomenclature (e.g., *psbA* vs *psba, trnH* vs *trnH-GUG*) may occasionally appear across species, these are readily identifiable in output tables and can be corrected by researchers based on their specific analytical requirements. Output is provided as timestamped Excel files enabling direct cross-species comparison (**Table S1**).

#### Module 2: Gene Table Generation

CGAS automatically generates formatted Word documents containing gene tables suitable for immediate inclusion in manuscripts or supplementary materials. Genes are categorized by functional class, displayed in italic typeface, and annotated with standardized superscripts indicating IR duplication or intron presence.

A key methodological feature is the explicit handling of trans-spliced genes. The *rps12* gene, distributed across multiple genomic regions, is correctly counted as a single functional gene rather than as an IR-duplicated locus. Additionally, CGAS employs biologically informed gene classification: *ycf3* and *ycf4* are categorized as photosynthesis assembly genes based on their established functional roles (Nellaepalli et al., 2018), despite being annotated as open reading frames in some references. To enhance clarity and reflect current nomenclature, genes are presented with both systematic and functional names where applicable (e.g., *ycf3* (*pafI*), *ycf4* (*pafII*), *psbN* (*psb1*), *psbZ* (*lhbA*)). An explanatory note is embedded in each output document to ensure transparency regarding these conventions. Representative output is shown in **Table S2**, with complete results for all analyzed species available in **Supplementary Data File 3**.

#### Module 3: Comparative Genome Analysis

This module characterizes chloroplast genome architecture by identifying LSC, SSC, and IR boundaries and calculating region-specific GC content, including overall genome GC%, LSC GC%, SSC GC%, and IR GC%. In addition, GC content is computed separately for protein-coding genes, tRNAs, and rRNAs, facilitating fine-scale structural and compositional comparisons among taxa. The results are given in **Table S3**.

#### Module 4: Relative Synonymous Codon Usage Analysis (RSCU)

All protein-coding sequences are extracted, validated, and parsed into codon triplets using the plastid genetic code. CGAS calculates absolute codon frequencies and normalized RSCU values, generating species-by-codon matrices appropriate for comparative and evolutionary analyses. By processing all genomes in a directory automatically, this module eliminates the need for repetitive manual processing of individual files. The results are presented as **Table S4** for comparative genomic analysis.

#### Module 5: Amino Acid Composition Analysis

Protein-coding genes are translated and summarized to generate amino acid composition profiles across the chloroplast proteome. Relative abundances of all 20 standard amino acids are calculated and exported in comparative matrix format, supporting downstream analyses of translational bias and proteome evolution. The results are presented as **Table S5** for comparative genomic analysis for all species.

#### Module 6: Substitution Analysis

Using pre-aligned FASTA inputs, CGAS scans nucleotide positions to identify substitution types. Each SNP is classified as a transition or transversion, and summary statistics, including Ts/Tv ratios, are calculated. Alignment integrity is validated prior to analysis to prevent spurious SNP calls.

The module produces per-alignment reports and merged summaries, enabling efficient comparison across multiple datasets. This step requires alignment using MAFFT or any other alignment software. Alignment files should be named descriptively (e.g., `reference_vs_species.fasta` or `species.fasta` if a single species is used as reference) as these names appear in output tables. Here, we avoid generating file names from the species aligned in pairwise alignment, as in large-scale comparative genomics, researchers mostly use a single species as a reference and align other species to the reference genome. The results are given in **Table S6**.

#### Module 7: Intron Analysis

Intron-containing CDS and tRNA genes are identified using compound feature locations. CGAS records intron coordinates, lengths, and counts per gene, with safeguards to exclude annotation artifacts and trans-spliced structures that could bias intron statistics. The longest reported intron in chloroplast genomes is found in the *trnK-UUU* gene, which is up to 2,500 bp; therefore, introns >10,000 bp are excluded to prevent misidentification of *rps12* trans-splicing as an artificial intron of up to 30 kbp. However, the *rps12* part in IR contains an intron, and this part is correctly detected.

Outputs are organized into separate Excel sheets for CDS and tRNA introns, facilitating clear interpretation (**Table S7**).

#### Module 8: Simple Sequence Repeat (SSR) Analysis

CGAS identifies and classifies simple sequence repeats (microsatellites) across chloroplast genomes using configurable repeat thresholds. By default, the module detects mononucleotide repeats with ≥10 units, dinucleotide repeats with ≥5 units, and tri-through hexanucleotide repeats with ≥3-4 units, though these thresholds are customizable. Each detected SSR is systematically classified by three criteria: genomic region (LSC, SSC, or IR), motif type (mono-, di-, tri-, tetra-, penta-, or hexanucleotide), and precise genomic location.

The module employs a sophisticated location annotation system that identifies whether SSRs occur within protein-coding sequences, tRNA genes, rRNA genes, introns, or intergenic spacers (IGS). To prevent spurious annotations, the algorithm excludes trans-spliced genes from intron analysis by filtering features with intron lengths exceeding 10,000 bp, similar to the intron analysis module. For ambiguous cases where SSRs span multiple features, the module prioritizes tRNA genes over protein-coding genes and applies a 50% overlap threshold to assign definitive locations.

Outputs are organized into three complementary Excel files: a detailed table listing all SSRs with their coordinates, motifs, and annotations; a summary table aggregating SSR counts by species, region type, and motif category (**Table S8**), individual species sheets for focused examination (**Table S9**), and placement of all species data systematically with clear species heading in a single Excel sheet for publication as supplementary material (**Table S10**), as some journals allow data in one Excel sheet instead of placing data in multiple Excel sheets. Species names are formatted in italics throughout all outputs for publication-ready presentation.

#### Module 9: Nucleotide Diversity Analysis

CGAS automatically extracts genes, intergenic spacer regions, and intronic regions from multiple chloroplast GenBank files, aligns them using MAFFT, and calculates nucleotide diversity (π) to quantify evolutionary rates and selection pressures across different genomic compartments. This module requires MAFFT to be installed and accessible in the system path for sequence alignment.

The module first extracts sequences while implementing sophisticated gene name normalization to handle inconsistent nomenclature—including tRNA anticodon notation variants (*trnK-UUU* vs *trnK_UUU* vs *trnK(UUU)*), gene synonyms (*ycf3/pafI, psbN/pbf1, ycf4/pafII, clpP/clpP1, psbZ/lhbA*), and case inconsistencies across annotations. This normalization ensures accurate ortholog grouping across species despite annotation heterogeneity.

For each orthologous sequence set, CGAS performs multiple sequence alignment using MAFFT and calculates pairwise nucleotide diversity using standard population genetic formulae. The diversity metric π represents the average number of nucleotide differences per site between sequences, providing a quantitative measure of sequence conservation. Only polymorphic sites with unambiguous nucleotide calls are included in calculations to prevent alignment gaps and ambiguous bases from inflating diversity estimates.

The module handles complex genomic features including multi-exon genes with proper intron boundary detection, excludes trans-spliced genes that could confound intron analysis, and generates intergenic spacer coordinates by identifying gaps between annotated features.

Results are exported in multiple formats: a comprehensive summary file with diversity statistics for all regions organized by type (genes, introns, intergenic spacers) along with mean, median, minimum, and maximum π values for each region category, facilitating comparative analysis of evolutionary constraints across functional gene and non-coding regions (**Table S11**). Another important output file presents results with genomic coordinates (**Table S12**); and R-compatible tab-delimited files structured for direct visualization in ggplot2 workflows (**Tables S13 and S14**).

## Results and Practical Performance

CGAS was validated using ten publicly available chloroplast genomes from six angiosperm families. This input data is provided as **Supplementary Data File 2**, and the output data is provided as **Supplementary Data File 2**, as mentioned before. Across datasets, the pipeline produced results consistent with established tools while substantially reducing hands-on time. For example, the counts of amino acids, codon usage, and substitutions were checked with results of Geneious Prime (Kearse et al., 2012) and MEGA (Kumar et al., 2018). In addition, the gene content, their position in genome, and pseudogenization were also visualized in Geneious along with genome size and GC content of each genomic region. For validation of intron results, all results were carefully checked in Geneious for each gene to ensure correct parsing of intron. The SSR analyses were compared using MISA-web (Beier et al., 2017) and CPStools (Huang et al., 2024); and also visualized in Geneious. On standard desktop hardware, complete analysis of a typical angiosperm chloroplast genome required only a few minutes, with linear scaling observed as genome numbers increased. Importantly, biologically informed logic—particularly for trans-spliced genes and IR duplication— prevented common analytical errors observed in manual or semi-automated workflows.

## Discussion

CGAS addresses a critical methodological gap in chloroplast genomics by unifying commonly required analyses into a single, automated framework. In contrast to existing tools, CGAS is explicitly designed for comprehensive comparative analysis and publication-ready output generation, with particular emphasis on analyses not covered by recent platforms such as CPStools and CPGview—including amino acid frequency profiling, intron characterization, publication-ready gene content tables, and formatted outputs for genomic region and intron statistics. Its focus on directory-based batch processing, biological accuracy (particularly for trans-spliced genes and IR-mediated duplications), and standardized output formats makes it particularly suitable for large-scale multi-species comparative studies. Compared to manual workflows requiring 2-3 hours per genome to complete all analyses and consolidate results into a single comparative file, CGAS processes typical datasets of 10-20 genomes in 2-5 minutes, representing a >90% reduction in analysis time.

The modular architecture and multiple execution modes (unified script, module imports, Jupyter notebooks, command-line interface, and Linux server deployment) ensure CGAS can be deployed across diverse computational environments, from individual researcher workstations to high-performance computing clusters. This flexibility accommodates users ranging from wet-lab biologists seeking straightforward analysis to bioinformaticians requiring integration into complex automated pipelines. To enhance accessibility for researchers with limited computational expertise, we provide comprehensive video tutorials demonstrating complete analysis workflows in addition to detailed README documentation, significantly lowering the barrier to adoption.

CGAS operates on well-annotated input GenBank files, focusing its analytical capabilities on downstream comparative analyses that complement rather than duplicate existing annotation and alignment tools. This design choice enables specialized attention to publication-ready output generation and biologically informed handling of complex features—analytical capabilities that represent the primary novelty of the suite.

## Conclusions

The Chloroplast Genome Analysis Suite provides a reproducible, automation-driven solution for comparative chloroplast genome analysis. By integrating nine essential analytical modules into a single pipeline and generating publication-ready outputs, CGAS substantially streamlines chloroplast genomics workflows, reducing analysis time by >90% compared to fragmented manual approaches. The suite is freely available and extensible, offering a robust methodological foundation for future comparative and evolutionary studies.

## Supporting information

Supplementary Date File 1

Supplementary Date File 2

Supplementary Date File 3

## Availability and Requirements

**Project name:** Chloroplast Genome Analysis Suite (CGAS)

**Project home page:** https://github.com/abdullah30/Chloroplast-Genome-Analysis-Suite-CGAS

**Operating system:** Platform independent

**Programming language:** Python ≥3.8

**Dependencies:** Biopython (≥1.79), Pandas (≥1.3), OpenPyXL (≥3.0), python-docx (≥0.8.11), MAFFT (for nucleotide diversity module)

**License:** MIT

## Declarations

### Authors’ contributions

Abdullah conceived the study, developed the software, and prepared the manuscript. Rushan Yan contributed to software testing and validation. Xiaoxuan Tian contributed to methodological refinement and manuscript editing. All authors read and approved the final manuscript.

## Acknowledgments

We thank the open-source bioinformatics community for maintaining foundational Python libraries (Biopython, Pandas, NumPy, OpenPyXL) that enabled this work.

## Supplementary Materials

**Supplementary Data File 1:** Individual Python scripts and unified analyzer.

**Supplementary Data File 2:** Complete input data, including GenBank files of 10 genomes and three FASTA alignments.

**Supplementary Data File 3:** Example output data generated through CGAS.

**Table S1:** Gene content and count summary statistics for all species.

**Table S2:** Representative gene table with functional classification.

**Table S3:** Comparative genome architecture: LSC/SSC/IR lengths and region-specific GC content, along with the GC content of genes.

**Table S4:** Relative synonymous codon usage (RSCU) values for all species.

**Table S5:** Amino acid composition profiles across chloroplast proteomes.

**Table S6:** Substitutions analysis with Ts/Tv ratios.

**Table S7:** Intron characteristics for CDS and tRNA genes.

**Table S8:** Summary of simple sequence repeats (SSRs) by region and motif type.

**Table S9:** SSR details of each species in a separate sheet.

**Table S10:** Consolidated SSR data for all species in a single Excel sheet.

**Table S11:** Nucleotide diversity (π) summary statistics by genomic region type.

**Table S12:** Nucleotide diversity results with genomic coordinates.

**Table S13:** R-compatible nucleotide diversity data of genes.

**Table S14:** R-compatible nucleotide diversity data of non-coding regions.

