## Supplementary material for "CGAS (Chloroplast Genome Analysis Suite): An Automated Python Pipeline for Comprehensive Comparative Chloroplast Genomics": Table S2

**Table S2. Representative Gene table with gene functional classification**

| **Category for genes** | **Group of genes** | **Name of genes** | **Gene number** |
| --- | --- | --- | --- |
| Self-replication | Large subunit of ribosome | *rpl14*, *rpl16**, *rpl2*^, a^*, *rpl20*, *rpl22*, *rpl23^a^*, *rpl32*, *rpl33*, *rpl36* | 11 |
|  | Small subunit of ribosome | *rps11*, *rps12**, *rps14*, *rps15*, *rps16**, *rps18*, *rps19*, *rps2*, *rps3*, *rps4*, *rps7^a^*, *rps8* | 13 |
|  | DNA dependent RNA polymerase | *rpoA*, *rpoB*, *rpoC1**, *rpoC2* | 4 |
|  | rRNA genes | *rrn16^a^*, *rrn23^a^*, *rrn4.5^a^*, *rrn5^a^* | 8 |
|  | tRNA genes | *trnA-UGC*^, a^*, *trnC-GCA*, *trnD-GUC*, *trnE-UUC*, *trnF-GAA*, *trnG-UCC*, *trnH-GUG*, *trnI-CAU^a^*, *trnI-GAU*^, a^*, *trnK-UUU**, *trnL-CAA^a^*, *trnL-UAA**, *trnL-UAG*, *trnM-CAU*, *trnN-GUU^a^*, *trnP-UGG*, *trnQ-UUG*, *trnR-ACG^a^*, *trnR-UCU*, *trnS-GCU*, *trnS-GGA*, *trnS-UGA*, *trnT-UGU*, *trnV-GAC^a^*, *trnV-UAC**, *trnW-CCA*, *trnY-GUA*, *trnfM-CAU* | 35 |
| Photosynthesis | Photosystem Ⅰ | *psaA*, *psaB*, *psaC*, *psaI*, *psaJ* | 5 |
|  | Photosystem Ⅱ | *psbA*, *psbB*, *psbC*, *psbD*, *psbE*, *psbF*, *psbH*, *psbI*, *psbJ*, *psbK*, *psbL*, *psbM*, *psbN (pbf1)*, *psbT*, *psbZ (lhbA)* | 15 |
|  | NADPH dehydrogenase | *ndhA**, *ndhB*^, a^*, *ndhC*, *ndhD*, *ndhE*, *ndhF*, *ndhG*, *ndhH*, *ndhI*, *ndhJ*, *ndhK* | 12 |
|  | Cytochrome b/f complex | *petA*, *petB**, *petD**, *petG*, *petL*, *petN* | 6 |
|  | Subunits of ATP synthase | *atpA*, *atpB*, *atpE*, *atpF**, *atpH*, *atpI* | 6 |
|  | Large subunit of Rubisco | *rbcL* | 1 |
|  | Photosynthesis assembly genes | *ycf3 (pafI)**, *ycf4 (pafII)* | 2 |
| Other genes | Protease | *clpP** | 1 |
|  | Maturase | *matK* | 1 |
|  | Envelop membrane protein | *cemA* | 1 |
|  | Subunit of Acetyl-CoA-carboxylase | *accD* | 1 |
|  | C-type cytochrome synthesis gene | *ccsA* | 1 |
|  | Translation initiation factor | *infA* | 1 |
|  | Conserved open reading frames | *ycf1^a^*, *ycf2^a^* | 4 |
|  |  | **Total number of genes** | 128 |

**Note:** *, ^a^ indicate genes containing introns and duplicated genes in inverted repeat (IR) regions, respectively. The *rps12* gene is a trans-spliced gene and is not marked as duplicated despite appearing in multiple locations.
