## Supplementary Date File 1 for "CGAS (Chloroplast Genome Analysis Suite): An Automated Python Pipeline for Comprehensive Comparative Chloroplast Genomics": chloroplast_unified_analyzer.py

```
#!/usr/bin/env python3
"""
UNIFIED CHLOROPLAST GENOME ANALYSIS SUITE
==========================================

Version: 2.1.0 (Modular Import System)

This script imports and runs 9 analysis modules.
Place all individual script files in the same folder as this unified script.

Modules:
1. Genome Gene Count and Summary
2. Gene Table Generation (Word)
3. Comparative Genome Analysis
4. Codon Usage Analysis (RSCU)
5. Amino Acid Composition Analysis
6. SNP/Substitution Analysis (FASTA)
7. Intron Analysis
8. SSR Analysis
9. Nucleotide Diversity Analysis (requires MAFFT)

Author: Abdullah
Date: December 2025

Usage:
    python chloroplast_unified_analyzer.py

Folder Structure:
    /home/abdullah/script/unified/
    ├── chloroplast_unified_analyzer.py  (this file)
    ├── module1_gene_count.py
    ├── module2_gene_table.py
    ├── module3_comparative_analysis.py
    ├── module4_codon_usage.py
    ├── module5_amino_acid.py
    ├── module6_snp_analysis.py
    ├── module7_intron_extraction.py
    ├── module8_ssr_analysis.py
    └── module9_diversity_analysis.py

Requirements:
    pip install biopython pandas openpyxl numpy python-docx
    
    For Module 9: sudo apt-get install mafft (or brew install mafft on macOS)
"""

import os
import sys
import subprocess
import importlib.util
from pathlib import Path

print("="*80)
print("CHLOROPLAST GENOME UNIFIED ANALYSIS SUITE v2.1.0")
print("="*80)
print()

# Get script directory
SCRIPT_DIR = Path(__file__).parent.absolute()

# Detect available files in CURRENT WORKING DIRECTORY (not script directory)
WORK_DIR = Path.cwd()
gb_files = [f for f in os.listdir(WORK_DIR) if f.endswith(('.gb', '.gbk', '.genbank', '.gbff'))]
fasta_files = [f for f in os.listdir(WORK_DIR) if f.endswith(('.fasta', '.fa', '.fna', '.fas'))]

print(f"Working directory: {WORK_DIR}")
print(f"Script directory: {SCRIPT_DIR}")
print()
print("Files detected:")
print(f"  GenBank files: {len(gb_files)}")
print(f"  FASTA files: {len(fasta_files)}")
print()

# Check MAFFT availability for Module 9
MAFFT_AVAILABLE = False
try:
    subprocess.run(["mafft", "--version"], 
                  stdout=subprocess.DEVNULL, 
                  stderr=subprocess.DEVNULL, 
                  check=True)
    MAFFT_AVAILABLE = True
    print("✓ MAFFT detected - Module 9 will run")
except (subprocess.CalledProcessError, FileNotFoundError):
    print("⚠ MAFFT not found - Module 9 will be skipped")

print()

# ============================================================================
# Dynamic Module Importer
# ============================================================================

def import_module_from_file(module_name, file_path):
    """
    Dynamically import a module from a file path.
    
    Args:
        module_name: Name to give the module
        file_path: Path to the .py file
    
    Returns:
        Imported module object or None if import fails
    """
    try:
        spec = importlib.util.spec_from_file_location(module_name, file_path)
        if spec is None:
            print(f"  ✗ Could not load spec for {file_path}")
            return None
        
        module = importlib.util.module_from_spec(spec)
        sys.modules[module_name] = module
        spec.loader.exec_module(module)
        return module
    except Exception as e:
        print(f"  ✗ Error importing {file_path}: {e}")
        return None


# ============================================================================
# Module Configuration
# ============================================================================

MODULES = {
    1: {
        'name': 'Gene Count and Summary',
        'file': 'module1_gene_count.py',
        'main_function': 'main',
        'requires': 'genbank',
        'min_files': 1
    },
    2: {
        'name': 'Gene Table Generation',
        'file': 'module2_gene_table.py',
        'main_function': 'main',
        'requires': 'genbank',
        'min_files': 1
    },
    3: {
        'name': 'Comparative Genome Analysis',
        'file': 'module3_comparative_analysis.py',
        'main_function': 'main',
        'requires': 'genbank',
        'min_files': 1
    },
    4: {
        'name': 'Codon Usage Analysis (RSCU)',
        'file': 'module4_codon_usage.py',
        'main_function': 'main',
        'requires': 'genbank',
        'min_files': 1
    },
    5: {
        'name': 'Amino Acid Composition',
        'file': 'module5_amino_acid.py',
        'main_function': 'main',
        'requires': 'genbank',
        'min_files': 1
    },
    6: {
        'name': 'SNP/Substitution Analysis',
        'file': 'module6_snp_analysis.py',
        'main_function': 'main',
        'requires': 'fasta',
        'min_files': 1
    },
    7: {
        'name': 'Intron Extraction',
        'file': 'module7_intron_extraction.py',
        'main_function': 'main',
        'requires': 'genbank',
        'min_files': 1
    },
    8: {
        'name': 'SSR Analysis',
        'file': 'module8_ssr_analysis.py',
        'main_function': 'run_module8',
        'requires': 'genbank',
        'min_files': 1,
        'kwargs': {
            'gb_folder': '.',
            'output_folder': None,
            'thresholds': {1: 10, 2: 5, 3: 4, 4: 3, 5: 3, 6: 3}
        }
    },
    9: {
        'name': 'Nucleotide Diversity',
        'file': 'module9_diversity_analysis.py',
        'main_function': 'main',
        'requires': 'genbank',
        'min_files': 2,
        'needs_mafft': True,
        'set_argv': True  # Need to set sys.argv with GenBank file list
    }
}


# ============================================================================
# Module Execution Functions
# ============================================================================

def check_requirements(module_num, config):
    """Check if module requirements are met."""
    reasons = []
    
    # Check file requirements
    if config['requires'] == 'genbank':
        if len(gb_files) < config['min_files']:
            reasons.append(f"Requires {config['min_files']}+ GenBank files (found {len(gb_files)})")
    elif config['requires'] == 'fasta':
        if len(fasta_files) < config['min_files']:
            reasons.append(f"Requires {config['min_files']}+ FASTA files (found {len(fasta_files)})")
    
    # Check MAFFT requirement
    if config.get('needs_mafft', False) and not MAFFT_AVAILABLE:
        reasons.append("Requires MAFFT to be installed")
    
    return reasons


def run_module(module_num, config):
    """Run a single module."""
    print("\n" + "-"*80)
    print(f"Module {module_num}: {config['name']}")
    print("-"*80)
    
    # Check requirements
    issues = check_requirements(module_num, config)
    if issues:
        print(f"⚠ Skipping: {', '.join(issues)}")
        return 'skipped'
    
    # Check if file exists
    module_path = SCRIPT_DIR / config['file']
    if not module_path.exists():
        print(f"⚠ File not found: {config['file']}")
        print(f"  Expected location: {module_path}")
        return 'skipped'
    
    # Import module
    print(f"  Loading {config['file']}...")
    module = import_module_from_file(f"module{module_num}", str(module_path))
    
    if module is None:
        print(f"✗ Failed to import module")
        return 'failed'
    
    # Get main function
    main_func_name = config['main_function']
    if not hasattr(module, main_func_name):
        print(f"✗ Function '{main_func_name}' not found in module")
        return 'failed'
    
    main_func = getattr(module, main_func_name)
    
    try:
        # Handle special cases
        if config.get('set_argv', False):
            # Module needs sys.argv set (like module 9)
            old_argv = sys.argv
            sys.argv = ["script"] + gb_files
            try:
                main_func()
            finally:
                sys.argv = old_argv
        elif 'kwargs' in config:
            # Module accepts keyword arguments (like module 8)
            main_func(**config['kwargs'])
        else:
            # Standard main() call
            main_func()
        
        print(f"✓ Module {module_num} completed")
        return 'success'
        
    except KeyboardInterrupt:
        print(f"\n⚠ Module {module_num} interrupted by user")
        raise
    except Exception as e:
        print(f"✗ Module {module_num} error: {e}")
        import traceback
        traceback.print_exc()
        return 'failed'


# ============================================================================
# Main Execution Controller
# ============================================================================

def run_all_modules():
    """Execute all available modules in sequence."""
    print("\n" + "="*80)
    print("STARTING UNIFIED ANALYSIS")
    print("="*80)
    
    results = {
        'success': 0,
        'skipped': 0,
        'failed': 0
    }
    
    # Run modules in order
    for module_num in sorted(MODULES.keys()):
        config = MODULES[module_num]
        result = run_module(module_num, config)
        
        if result in results:
            results[result] += 1
    
    # Final Summary
    print("\n" + "="*80)
    print("ANALYSIS COMPLETE")
    print("="*80)
    print(f"✓ Modules completed: {results['success']}")
    print(f"⚠ Modules skipped: {results['skipped']}")
    print(f"✗ Modules failed: {results['failed']}")
    print("="*80)
    print()
    print("Output locations:")
    print(f"  • Current directory: {WORK_DIR}")
    print(f"  • Excel/Word files with timestamps")
    print(f"  • Module8_SSR_Analysis_* folder")
    print(f"  • Module9_Nucleotide_Diversity folder")
    print()
    
    if not MAFFT_AVAILABLE:
        print("💡 Tip: Install MAFFT to enable Module 9 (Nucleotide Diversity)")
        print("   Ubuntu/Debian: sudo apt-get install mafft")
        print("   macOS: brew install mafft")
    
    print("="*80)


# ============================================================================
# Entry Point
# ============================================================================

if __name__ == "__main__":
    try:
        run_all_modules()
    except KeyboardInterrupt:
        print("\n\n⚠ Analysis interrupted by user")
        sys.exit(0)
    except Exception as e:
        print(f"\n\n✗ FATAL ERROR: {e}")
        import traceback
        traceback.print_exc()
        sys.exit(1)
```
