## Supplementary Date File 1 for "CGAS (Chloroplast Genome Analysis Suite): An Automated Python Pipeline for Comprehensive Comparative Chloroplast Genomics": module1_gene_count.py

```
#!/usr/bin/env python3
"""
Chloroplast Genome Gene Comparative Analysis Tool
==================================================

This script performs comprehensive analysis of chloroplast genome annotations from
GenBank files. It identifies genes, classifies them as functional or pseudogenes, detects IR-mediated duplications,
and produces publication-ready Excel reports.

Author: Abdullah
Date: December 2025
Version: 1.0

Requirements:
    - Python 3.7+
    - biopython
    - pandas
    - openpyxl (for Excel writing)

Usage:
    Place this script in a folder containing GenBank (.gb, .gbk, .genbank) files
    and run:
        python chloroplast_analyzer.py
    
    Output will be generated as an Excel file with timestamp in the same directory.

Citation:
    If you use this tool in your research, please cite [appropriate reference]
"""

import os
import re
from collections import defaultdict
from datetime import datetime
from typing import List, Tuple, Dict, Set
from Bio import SeqIO
from Bio.SeqFeature import SeqFeature
import pandas as pd
from openpyxl.styles import Font


# ============================================================================
# CONFIGURATION
# ============================================================================

# Automatically use current working directory
WORKING_DIR = os.getcwd()

# Generate timestamped output filename for version control
TIMESTAMP = datetime.now().strftime("%Y%m%d_%H%M%S")
OUTPUT_FILE = os.path.join(WORKING_DIR, f"Chloroplast_Gene_Analysis_{TIMESTAMP}.xlsx")

# Gap tolerance for merging nearby gene features (bp)
GAP_TOLERANCE = 0

# GenBank file extensions to process
GENBANK_EXTENSIONS = ('.gb', '.gbk', '.genbank')


# ============================================================================
# UTILITY FUNCTIONS
# ============================================================================

def extract_species_name(filename: str) -> str:
    """
    Extract clean species name from GenBank filename.
    
    Removes file extensions (.gb, .gbk, .genbank) and standardizes the name
    for publication-quality output.
    
    Args:
        filename: Original GenBank filename
        
    Returns:
        Clean species name without extension
        
    Examples:
        'Solanum_lycopersicum.gb' -> 'Solanum_lycopersicum'
        'Species_name.gbk' -> 'Species_name'
    """
    # Remove common GenBank extensions
    name = filename
    for ext in GENBANK_EXTENSIONS:
        if name.lower().endswith(ext):
            name = name[:-len(ext)]
            break
    
    return name.strip()

def get_gene_name_from_qualifiers(qualifiers: Dict) -> str:
    """
    Extract gene name from GenBank feature qualifiers.
    
    Searches through common qualifier fields in priority order to identify
    the most appropriate gene name. This ensures consistent naming across
    different annotation styles.
    
    Args:
        qualifiers: Dictionary of GenBank feature qualifiers
        
    Returns:
        Gene name as string, or None if no suitable name found
        
    Priority order:
        1. gene - Standard gene symbol (e.g., 'rbcL', 'psbA')
        2. locus_tag - Systematic locus identifier
        3. product - Gene product description
        4. protein_id - Protein accession
        5. note - Additional annotations
    """
    for key in ("gene", "locus_tag", "product", "protein_id", "note"):
        if key in qualifiers and qualifiers[key]:
            value = qualifiers[key][0].strip()
            if value:
                # Replace spaces with underscores for consistent naming
                return value.replace(" ", "_")
    return None


def detect_inverted_repeat_regions(record: SeqIO.SeqRecord) -> Tuple[List[Tuple[int, int]], 
                                                                       List[Tuple[int, int]]]:
    """
    Identify inverted repeat regions (IRa and IRb) in chloroplast genome.
    
    Chloroplast genomes typically have two large inverted repeats (IRa and IRb)
    that separate the large and small single-copy regions. This function detects
    these regions from GenBank annotations.
    
    Args:
        record: BioPython SeqRecord object from GenBank file
        
    Returns:
        Tuple of two lists: (IRa_ranges, IRb_ranges)
        Each range is a tuple of (start, end) coordinates
        
    Notes:
        - Searches 'repeat_region' and 'misc_feature' annotations
        - Identifies IR regions by keywords: "inverted repeat", "ira", "irb"
        - Merges overlapping or adjacent annotations
    """
    ira_ranges = []
    irb_ranges = []
    
    # Search all features for IR annotations
    for feature in record.features:
        if feature.type.lower() in ("repeat_region", "misc_feature"):
            # Collect all descriptive text from multiple qualifier fields
            annotation_text = []
            for key in ("note", "product", "description"):
                if key in feature.qualifiers:
                    annotation_text.append(" ".join(feature.qualifiers[key]))
            
            combined_text = " ".join(annotation_text).lower()
            
            # Check if this feature describes an inverted repeat
            if "inverted repeat" in combined_text or "ira" in combined_text or "irb" in combined_text:
                span = (int(feature.location.start), int(feature.location.end))
                
                # Classify as IRa or IRb based on annotation text
                if "ira" in combined_text or "ir a" in combined_text:
                    ira_ranges.append(span)
                elif "irb" in combined_text or "ir b" in combined_text:
                    irb_ranges.append(span)
    
    # Merge overlapping or adjacent ranges
    return _merge_genomic_ranges(ira_ranges), _merge_genomic_ranges(irb_ranges)


def _merge_genomic_ranges(ranges: List[Tuple[int, int]]) -> List[Tuple[int, int]]:
    """
    Merge overlapping or adjacent genomic coordinate ranges.
    
    Args:
        ranges: List of (start, end) coordinate tuples
        
    Returns:
        List of merged (start, end) tuples with no overlaps
        
    Example:
        [(100, 200), (150, 250), (300, 400)] -> [(100, 250), (300, 400)]
    """
    if not ranges:
        return []
    
    # Sort ranges by start position
    sorted_ranges = sorted(ranges)
    merged = [list(sorted_ranges[0])]
    
    for start, end in sorted_ranges[1:]:
        # Check if current range overlaps or is adjacent to last merged range
        if start <= merged[-1][1] + 1:
            # Extend the last merged range
            merged[-1][1] = max(merged[-1][1], end)
        else:
            # Add as new separate range
            merged.append([start, end])
    
    return [(start, end) for start, end in merged]


def check_span_overlap(span1: Tuple[int, int], span2: Tuple[int, int]) -> bool:
    """
    Test if two genomic coordinate spans overlap.
    
    Args:
        span1: First span as (start, end)
        span2: Second span as (start, end)
        
    Returns:
        True if spans overlap, False otherwise
    """
    start1, end1 = span1
    start2, end2 = span2
    return not (end1 < start2 or end2 < start1)


def merge_feature_spans(spans: List[Tuple[int, int]], 
                        gap_tolerance: int = 0) -> List[Tuple[int, int]]:
    """
    Merge gene feature spans that are within gap_tolerance of each other.
    
    Some genes may be annotated as multiple features (e.g., exons). This function
    merges features that belong to the same gene based on proximity.
    
    Args:
        spans: List of (start, end) coordinate tuples
        gap_tolerance: Maximum gap (bp) to bridge when merging
        
    Returns:
        List of merged (start, end) tuples
    """
    if not spans:
        return []
    
    sorted_spans = sorted(spans)
    merged = [list(sorted_spans[0])]
    
    for start, end in sorted_spans[1:]:
        # Check if within gap tolerance of previous span
        if start <= merged[-1][1] + gap_tolerance + 1:
            merged[-1][1] = max(merged[-1][1], end)
        else:
            merged.append([start, end])
    
    return [(start, end) for start, end in merged]


def classify_gene_locus(features: List[SeqFeature]) -> str:
    """
    Classify a gene locus as functional or pseudogene.
    
    Classification logic:
        - Functional: Has CDS, tRNA, or rRNA with product annotation
        - Pseudogene: Explicitly marked as pseudo OR lacks protein-coding features
    
    Args:
        features: List of BioPython SeqFeature objects for this locus
        
    Returns:
        Classification string: "functional" or "pseudogene"
        
    Notes:
        - Prioritizes explicit pseudogene annotations
        - Considers both feature types and qualifier content
        - tRNA/rRNA require product annotation to be considered functional
    """
    has_cds = False
    has_trna = False
    has_rrna = False
    has_product = False
    explicit_pseudo = False
    
    for feature in features:
        qualifiers = feature.qualifiers
        
        # Check CDS (protein-coding) features
        if feature.type == "CDS":
            if "pseudo" in qualifiers:
                explicit_pseudo = True
            
            product_text = " ".join(
                qualifiers.get("product", []) + qualifiers.get("note", [])
            ).lower()
            
            if "pseudo" in product_text or "pseudogene" in product_text:
                explicit_pseudo = True
            else:
                has_cds = True
        
        # Check tRNA features
        if feature.type == "tRNA":
            has_trna = True
            product_text = " ".join(
                qualifiers.get("product", []) + qualifiers.get("note", [])
            ).strip()
            if product_text:
                has_product = True
        
        # Check rRNA features
        if feature.type == "rRNA":
            has_rrna = True
            product_text = " ".join(
                qualifiers.get("product", []) + qualifiers.get("note", [])
            ).strip()
            if product_text:
                has_product = True
        
        # Check for product annotation (any feature type)
        if "product" in qualifiers and qualifiers["product"]:
            product_text = " ".join(qualifiers["product"]).strip().lower()
            if product_text and "pseudo" not in product_text and "pseudogene" not in product_text:
                has_product = True
            elif "pseudo" in product_text or "pseudogene" in product_text:
                explicit_pseudo = True
        
        # Check note field for pseudogene annotations
        if "note" in qualifiers and qualifiers["note"]:
            note_text = " ".join(qualifiers["note"]).lower()
            if "pseudogene" in note_text or "pseudo" in note_text:
                explicit_pseudo = True
    
    # Classification decision
    if has_cds or has_trna or has_rrna or has_product:
        return "functional"
    return "pseudogene"


# ============================================================================
# MAIN ANALYSIS FUNCTIONS
# ============================================================================

def analyze_genbank_file(filepath: str) -> List[Dict]:
    """
    Analyze a single GenBank file for gene content and IR duplications.
    
    This is the main analysis function that:
        1. Detects IR regions (IRa and IRb)
        2. Identifies all genes and their features
        3. Classifies genes as functional or pseudogenes
        4. Determines copy number and IR duplication status
    
    Args:
        filepath: Path to GenBank file
        
    Returns:
        List of dictionaries, each containing analysis results for one gene:
            - Genome: Filename of source
            - Gene Name: Gene identifier
            - Copies: Number of copies detected
            - Functional / Pseudogene: Gene status
            - Duplicate Status: Detailed copy number and location info
    """
    # Parse GenBank file
    record = SeqIO.read(filepath, "genbank")
    
    # Detect inverted repeat regions
    ira_ranges, irb_ranges = detect_inverted_repeat_regions(record)
    
    # Collect all gene-related features
    gene_features = defaultdict(list)
    
    for feature in record.features:
        # Only process gene-related feature types
        if feature.type not in ("CDS", "tRNA", "rRNA", "gene"):
            continue
        
        # Get or generate gene name
        gene_name = get_gene_name_from_qualifiers(feature.qualifiers)
        if not gene_name:
            # Generate systematic name for unannotated ORFs
            start = int(feature.location.start)
            end = int(feature.location.end)
            strand = "-" if feature.location.strand == -1 else "+"
            gene_name = f"ORF_{start}_{end}_{strand}"
        
        gene_features[gene_name].append(feature)
    
    # Analyze each gene
    results = []
    for gene_name, features in sorted(gene_features.items()):
        # Extract genomic coordinates for all features of this gene
        span_feature_pairs = [
            (int(f.location.start), int(f.location.end), f) 
            for f in features
        ]
        spans = [(start, end) for start, end, _ in span_feature_pairs]
        
        # Merge nearby features into loci (handles multi-exon genes)
        locus_spans = merge_feature_spans(spans, gap_tolerance=GAP_TOLERANCE)
        
        # Analyze each locus separately
        locus_info_list = []
        for locus_span in locus_spans:
            locus_start, locus_end = locus_span
            
            # Get all features overlapping this locus
            locus_features = [
                feature for start, end, feature in span_feature_pairs
                if not (end < locus_start or start > locus_end)
            ]
            
            # Classify locus as functional or pseudogene
            classification = classify_gene_locus(locus_features)
            
            # Determine genomic region (IRa, IRb, or single copy)
            in_ira = any(check_span_overlap(locus_span, ir) for ir in ira_ranges)
            in_irb = any(check_span_overlap(locus_span, ir) for ir in irb_ranges)
            
            if in_ira and in_irb:
                region = "IR(both)"  # Rare: spans both IR regions
            elif in_ira:
                region = "IRa"
            elif in_irb:
                region = "IRb"
            else:
                region = "single_copy"
            
            locus_info_list.append({
                "span": locus_span,
                "classification": classification,
                "region": region
            })
        
        # Summarize gene-level statistics
        total_copies = len(locus_info_list)
        regions = {locus["region"] for locus in locus_info_list}
        is_ir_duplicated = ("IRa" in regions) and ("IRb" in regions)
        
        functional_copies = sum(
            1 for locus in locus_info_list 
            if locus["classification"] == "functional"
        )
        pseudogene_copies = sum(
            1 for locus in locus_info_list 
            if locus["classification"] == "pseudogene"
        )
        
        # Overall gene status (functional if any copy is functional)
        gene_status = "functional" if functional_copies > 0 else "pseudogene"
        
        # Generate detailed duplication status string
        if is_ir_duplicated:
            if functional_copies > 1 and pseudogene_copies == 0:
                dup_status = f"{functional_copies} functional copies (IRa+IRb)"
            elif functional_copies >= 1 and pseudogene_copies >= 1:
                dup_status = (f"{functional_copies} functional + "
                             f"{pseudogene_copies} pseudogene copies (IRa+IRb)")
            elif functional_copies == 0 and pseudogene_copies >= 1:
                dup_status = f"{pseudogene_copies} pseudogene copies (IRa+IRb)"
            else:
                dup_status = f"{total_copies} copies (IRa+IRb)"
        else:
            if total_copies == 1:
                dup_status = "single copy"
            else:
                if functional_copies >= 1 and pseudogene_copies == 0:
                    dup_status = f"{functional_copies} functional copies (not IR-duplicated)"
                elif functional_copies >= 1 and pseudogene_copies >= 1:
                    dup_status = (f"{functional_copies} functional + "
                                 f"{pseudogene_copies} pseudogene copies (not IR-duplicated)")
                else:
                    dup_status = f"{pseudogene_copies} pseudogene copies (not IR-duplicated)"
        
        # Store results for this gene
        results.append({
            "Genome": extract_species_name(os.path.basename(filepath)),
            "Gene Name": gene_name,
            "Copies": total_copies,
            "Functional / Pseudogene": gene_status,
            "Duplicate Status": dup_status
        })
    
    return results


def generate_summary_statistics(all_results: List[Dict], 
                                genome_gene_map: Dict[str, Set[str]]) -> pd.DataFrame:
    """
    Generate genome-level summary statistics.
    
    Calculates aggregate statistics for each genome:
        - Total unique genes
        - Number of functional genes
        - Number of pseudogenes
        - Number of IR-duplicated genes
    
    Args:
        all_results: List of all gene analysis results
        genome_gene_map: Dictionary mapping genome names to their gene sets
        
    Returns:
        DataFrame with summary statistics per genome
    """
    summary_data = []
    
    for genome_name, gene_set in genome_gene_map.items():
        genome_results = [r for r in all_results if r["Genome"] == genome_name]
        
        # Count unique functional and pseudogene genes
        functional_genes = set()
        pseudogene_genes = set()
        ir_duplicated_count = 0
        
        for result in genome_results:
            if result["Functional / Pseudogene"] == "functional":
                functional_genes.add(result["Gene Name"])
            elif result["Functional / Pseudogene"] == "pseudogene":
                pseudogene_genes.add(result["Gene Name"])
            
            # Count IR-duplicated genes
            if "IRa+IRb" in result["Duplicate Status"]:
                ir_duplicated_count += 1
        
        summary_data.append({
            "Genome": genome_name,
            "Total Genes": len(gene_set),
            "Functional Genes": len(functional_genes),
            "Pseudogene Genes": len(pseudogene_genes),
            "IR-duplicated Genes": ir_duplicated_count
        })
    
    return pd.DataFrame(
        summary_data,
        columns=["Genome", "Total Genes", "Functional Genes", 
                "Pseudogene Genes", "IR-duplicated Genes"]
    )


def identify_unique_genes(all_results: List[Dict], 
                         genome_gene_map: Dict[str, Set[str]]) -> Tuple[pd.DataFrame, pd.DataFrame]:
    """
    Identify genes with unique presence/absence patterns.
    
    Returns two types of unique genes:
    1. Genes present in only ONE genome (genome-specific)
    2. Genes missing from only ONE genome (nearly universal)
    
    Args:
        all_results: List of all gene analysis results
        genome_gene_map: Dictionary mapping genome names to their gene sets
        
    Returns:
        Tuple of two DataFrames: (genome_specific_genes, nearly_universal_genes)
    """
    genome_specific_data = []
    nearly_universal_data = []
    
    # Get all unique genes across all genomes
    all_genes = set()
    for genes in genome_gene_map.values():
        all_genes.update(genes)
    
    # For each gene, count how many genomes contain it
    gene_presence = defaultdict(list)
    for genome_name, gene_set in genome_gene_map.items():
        for gene_name in all_genes:
            if gene_name in gene_set:
                gene_presence[gene_name].append(genome_name)
    
    total_genomes = len(genome_gene_map)
    
    # Identify genome-specific genes (present in only 1 genome)
    for gene_name, genomes_with_gene in gene_presence.items():
        if len(genomes_with_gene) == 1:
            genome_name = genomes_with_gene[0]
            gene_result = next(
                r for r in all_results 
                if r["Genome"] == genome_name and r["Gene Name"] == gene_name
            )
            genome_specific_data.append(gene_result)
    
    # Identify nearly universal genes (missing from only 1 genome)
    if total_genomes > 2:  # Only meaningful with 3+ genomes
        for gene_name, genomes_with_gene in gene_presence.items():
            if len(genomes_with_gene) == total_genomes - 1:
                # Find which genome is missing this gene
                missing_genome = None
                for genome_name in genome_gene_map.keys():
                    if genome_name not in genomes_with_gene:
                        missing_genome = genome_name
                        break
                
                # Add entry for each genome that HAS the gene
                for genome_name in genomes_with_gene:
                    gene_result = next(
                        r for r in all_results 
                        if r["Genome"] == genome_name and r["Gene Name"] == gene_name
                    ).copy()
                    gene_result["Missing_From"] = missing_genome
                    nearly_universal_data.append(gene_result)
    
    df_specific = pd.DataFrame(
        genome_specific_data,
        columns=["Genome", "Gene Name", "Copies", 
                "Functional / Pseudogene", "Duplicate Status"]
    )
    
    df_nearly_universal = pd.DataFrame(
        nearly_universal_data,
        columns=["Genome", "Gene Name", "Copies", 
                "Functional / Pseudogene", "Duplicate Status", "Missing_From"]
    )
    
    return df_specific, df_nearly_universal


def apply_publication_formatting(writer: pd.ExcelWriter, sheet_name: str):
    """
    Apply publication-quality formatting to Excel worksheet.
    
    Formatting includes:
        - Auto-adjusted column widths
        - Frozen header row
        - Italic formatting for species names (Genome column)
        - Italic formatting for gene names (Gene Name column)
    
    Args:
        writer: pandas ExcelWriter object
        sheet_name: Name of the worksheet to format
    """
    worksheet = writer.sheets[sheet_name]
    italic_font = Font(italic=True)
    
    # Get column indices for species and gene names
    genome_col = None
    gene_col = None
    missing_col = None
    
    # Find column letters for Genome, Gene Name, and Missing_From
    for col_idx, column in enumerate(worksheet.iter_cols(1, worksheet.max_column, 1, 1), start=1):
        header_value = column[0].value
        if header_value == "Genome":
            genome_col = col_idx
        elif header_value == "Gene Name":
            gene_col = col_idx
        elif header_value == "Missing_From":
            missing_col = col_idx
    
    # Apply italic formatting to species names and gene names
    for row_idx in range(2, worksheet.max_row + 1):  # Start from row 2 (skip header)
        # Italicize species names in Genome column
        if genome_col:
            cell = worksheet.cell(row=row_idx, column=genome_col)
            cell.font = italic_font
        
        # Italicize gene names in Gene Name column
        if gene_col:
            cell = worksheet.cell(row=row_idx, column=gene_col)
            cell.font = italic_font
        
        # Italicize species names in Missing_From column (if exists)
        if missing_col:
            cell = worksheet.cell(row=row_idx, column=missing_col)
            if cell.value:  # Only if not empty
                cell.font = italic_font
    
    # Auto-adjust column widths
    for column in worksheet.columns:
        max_length = 0
        column_letter = column[0].column_letter
        
        for cell in column:
            try:
                if cell.value and len(str(cell.value)) > max_length:
                    max_length = len(str(cell.value))
            except:
                pass
        
        # Set width with reasonable limits
        adjusted_width = min(max_length + 3, 50)
        worksheet.column_dimensions[column_letter].width = adjusted_width
    
    # Freeze header row for easy scrolling
    worksheet.freeze_panes = worksheet['A2']


# ============================================================================
# MAIN EXECUTION
# ============================================================================

def main():
    """
    Main execution function.
    
    Workflow:
        1. Find all GenBank files in current directory
        2. Analyze each file
        3. Generate summary statistics
        4. Identify unique genes
        5. Write publication-ready Excel report
    """
    print("=" * 70)
    print("Chloroplast Genome Gene Comparative Analysis")
    print("=" * 70)
    print(f"\nWorking directory: {WORKING_DIR}")
    
    # Find all GenBank files
    genbank_files = sorted([
        f for f in os.listdir(WORKING_DIR)
        if f.lower().endswith(GENBANK_EXTENSIONS)
    ])
    
    if not genbank_files:
        print(f"\nERROR: No GenBank files found in {WORKING_DIR}")
        print(f"Supported extensions: {', '.join(GENBANK_EXTENSIONS)}")
        return
    
    print(f"\nFound {len(genbank_files)} GenBank file(s):")
    for filename in genbank_files:
        print(f"  - {filename}")
    
    # Process all files
    print("\n" + "-" * 70)
    print("Processing files...")
    print("-" * 70)
    
    all_results = []
    genome_gene_map = defaultdict(set)
    
    for genbank_file in genbank_files:
        filepath = os.path.join(WORKING_DIR, genbank_file)
        species_name = extract_species_name(genbank_file)
        print(f"\nAnalyzing: {species_name}")
        
        try:
            results = analyze_genbank_file(filepath)
            all_results.extend(results)
            genome_gene_map[species_name] = set(r["Gene Name"] for r in results)
            print(f"  ✓ Found {len(results)} genes")
        except Exception as e:
            print(f"  ✗ Error processing file: {str(e)}")
            continue
    
    if not all_results:
        print("\nERROR: No results generated. Check your GenBank files.")
        return
    
    # Generate analysis outputs
    print("\n" + "-" * 70)
    print("Generating analysis reports...")
    print("-" * 70)
    
    # 1. Complete gene table
    df_gene_table = pd.DataFrame(
        all_results,
        columns=["Genome", "Gene Name", "Copies", 
                "Functional / Pseudogene", "Duplicate Status"]
    )
    
    # 2. Summary statistics
    df_summary = generate_summary_statistics(all_results, genome_gene_map)
    
    # 3. Unique genes (genome-specific and nearly universal)
    df_genome_specific, df_nearly_universal = identify_unique_genes(all_results, genome_gene_map)
    
    # Write Excel file with publication-quality formatting
    print(f"\nWriting results to: {OUTPUT_FILE}")
    
    with pd.ExcelWriter(OUTPUT_FILE, engine='openpyxl') as writer:
        # Write each sheet
        df_gene_table.to_excel(writer, sheet_name="Gene_Table", index=False)
        df_summary.to_excel(writer, sheet_name="Summary", index=False)
        df_genome_specific.to_excel(writer, sheet_name="Genome_Specific_Genes", index=False)
        if not df_nearly_universal.empty:
            df_nearly_universal.to_excel(writer, sheet_name="Nearly_Universal_Genes", index=False)
        
        # Apply publication formatting to all sheets
        for sheet_name in writer.sheets:
            apply_publication_formatting(writer, sheet_name)
    
    # Print summary
    print("\n" + "=" * 70)
    print("ANALYSIS COMPLETE")
    print("=" * 70)
    print(f"\nTotal genomes analyzed: {len(genome_gene_map)}")
    print(f"Total genes identified: {len(set(r['Gene Name'] for r in all_results))}")
    print(f"Total gene records: {len(all_results)}")
    
    print("\nOutput file contains up to 4 sheets:")
    print("  1. Gene_Table: Complete gene catalog with duplication status")
    print("  2. Summary: Genome-level statistics")
    print("  3. Genome_Specific_Genes: Genes found in only ONE genome")
    if not df_nearly_universal.empty:
        print("  4. Nearly_Universal_Genes: Genes missing from only ONE genome")
    
    print(f"\nResults saved to: {OUTPUT_FILE}")
    print("\n" + "=" * 70)


if __name__ == "__main__":
    main()
```
