## Supplementary Date File 1 for "CGAS (Chloroplast Genome Analysis Suite): An Automated Python Pipeline for Comprehensive Comparative Chloroplast Genomics": module2_gene_table.py

```
import pandas as pd
from Bio import SeqIO
from docx import Document
from docx.shared import Pt, Inches, RGBColor
from docx.enum.text import WD_ALIGN_PARAGRAPH
from docx.oxml.ns import qn
from docx.oxml import OxmlElement
import os
import glob

# Mapping for showing both names
name_map = {
    "pafI": "ycf3 (pafI)",
    "pafII": "ycf4 (pafII)", 
    "lhbA": "psbZ (lhbA)",
    "pbf1": "psbN (pbf1)",
    "ycf3": "ycf3 (pafI)",
    "ycf4": "ycf4 (pafII)",
    "psbZ": "psbZ (lhbA)",
    "psbN": "psbN (pbf1)"
}

# Function to get display name
def get_display_name(gene_name):
    """Convert gene name to display format"""
    return name_map.get(gene_name, gene_name)

# Function to check if a gene or tRNA has introns
def has_introns(features, gene_name, feature_type="CDS"):
    gene_features = [
        f for f in features
        if f.type == feature_type and gene_name in f.qualifiers.get("gene", [])
    ]
    for feature in gene_features:
        if hasattr(feature.location, "parts") and len(feature.location.parts) > 1:
            return True
    return False

# Extract gene content
def extract_gene_content(genbank_file):
    gene_data = {
        "Category for genes": [],
        "Group of genes": [],
        "Name of genes": [],
        "Amount": []
    }

    intron_containing_cds = set()
    intron_containing_trna = set()

    for record in SeqIO.parse(genbank_file, "genbank"):
        gene_count = {}

        for feature in record.features:
            if feature.type == "gene":
                gene_name = feature.qualifiers.get("gene", [""])[0]

                if gene_name:
                    # Convert to display name
                    display_name = get_display_name(gene_name)

                    # Check introns on ORIGINAL gene name
                    if has_introns(record.features, gene_name, "CDS"):
                        intron_containing_cds.add(display_name)
                        display_name += "*"
                    elif has_introns(record.features, gene_name, "tRNA"):
                        intron_containing_trna.add(display_name)
                        display_name += "*"

                    # Count occurrences
                    gene_count[display_name] = gene_count.get(display_name, 0) + 1

        # Define gene categories
        categories = {
            "Self-replication": [
                ("Large subunit of ribosome", "rpl"),
                ("Small subunit of ribosome", "rps"),
                ("DNA dependent RNA polymerase", "rpo"),
                ("rRNA genes", "rrn"),
                ("tRNA genes", "trn")
            ],
            "Photosynthesis": [
                ("Photosystem Ⅰ", "psa"),
                ("Photosystem Ⅱ", "psb"),
                ("NADPH dehydrogenase", "ndh"),
                ("Cytochrome b/f complex", "pet"),
                ("Subunits of ATP synthase", "atp"),
                ("Large subunit of Rubisco", "rbc"),
                # ONLY ycf3 and ycf4 in Photosynthesis assembly genes
                ("Photosynthesis assembly genes", ["ycf3 (pafI)", "ycf4 (pafII)"])
            ],
            "Other genes": [
                ("Protease", "clp"),
                ("Maturase", "mat"),
                ("Envelop membrane protein", "cem"),
                ("Subunit of Acetyl-CoA-carboxylase", "acc"),
                ("C-type cytochrome synthesis gene", "ccs"),
                ("Translation initiation factor", "infA"),
                # Conserved open reading frames (excluding ycf3 and ycf4)
                ("Conserved open reading frames", lambda gene: gene.startswith("ycf") 
                 and not gene.startswith("ycf3") and not gene.startswith("ycf4"))
            ]
        }

        categorized_genes = set()

        for category, groups in categories.items():
            for group_name, pattern in groups:
                matched_genes = {}
                
                # Handle different pattern types
                if callable(pattern):
                    # Use callable function for matching
                    for k, v in gene_count.items():
                        k_no_star = k.replace("*", "")
                        if pattern(k_no_star):
                            matched_genes[k] = v
                elif isinstance(pattern, list):
                    # Exact match for list of specific gene names
                    for k, v in gene_count.items():
                        k_no_star = k.replace("*", "")
                        if k_no_star in pattern:
                            matched_genes[k] = v
                else:
                    # String prefix match
                    for k, v in gene_count.items():
                        k_no_star = k.replace("*", "")
                        if k_no_star.startswith(pattern):
                            matched_genes[k] = v
                
                if matched_genes:
                    categorized_genes.update(matched_genes.keys())
                    
                    sorted_genes = sorted(matched_genes.items())
                    
                    gene_data["Category for genes"].append(category)
                    gene_data["Group of genes"].append(group_name)
                    # Use superscript 'a' for duplicates instead of (2)
                    # Exclude rps12 from duplicate marking (trans-spliced gene)
                    gene_data["Name of genes"].append(
                        ", ".join([f"{k}^a" if v > 1 and "rps12" not in k.lower() else k for k, v in sorted_genes])
                    )
                    # Count rps12 as 1 gene (trans-spliced), others as their actual count
                    total_count = sum([1 if "rps12" in k.lower() else v for k, v in matched_genes.items()])
                    gene_data["Amount"].append(total_count)

        # Remaining genes (Excluding genes)
        excluding_genes = {k: v for k, v in gene_count.items()
                          if k not in categorized_genes}

        if excluding_genes:
            gene_data["Category for genes"].append("Excluding genes")
            gene_data["Group of genes"].append("Excluding genes")
            sorted_excluding = sorted(excluding_genes.items())
            # Exclude rps12 from duplicate marking (trans-spliced gene)
            gene_data["Name of genes"].append(
                ", ".join([f"{k}^a" if v > 1 and "rps12" not in k.lower() else k for k, v in sorted_excluding])
            )
            # Count rps12 as 1 gene (trans-spliced), others as their actual count
            total_count = sum([1 if "rps12" in k.lower() else v for k, v in excluding_genes.items()])
            gene_data["Amount"].append(total_count)

    # Add total row
    total_genes = sum(gene_data["Amount"])
    gene_data["Category for genes"].append("")
    gene_data["Group of genes"].append("")
    gene_data["Name of genes"].append("Total number of genes")
    gene_data["Amount"].append(total_genes)

    return pd.DataFrame(gene_data), {
        "CDS with introns": list(intron_containing_cds),
        "tRNA with introns": list(intron_containing_trna)
    }

# Function to add superscript
def add_superscript(run, text):
    """Add superscript formatting to a run"""
    run.font.superscript = True
    run.text = text

# Function to set cell borders
def set_cell_border(cell, **kwargs):
    """Set cell borders for professional table appearance"""
    tc = cell._element
    tcPr = tc.get_or_add_tcPr()
    
    for edge in ('top', 'left', 'bottom', 'right'):
        edge_tag = 'w:{}'.format(edge)
        edge_element = tcPr.find(qn(edge_tag))
        if edge_element is None:
            edge_element = OxmlElement(edge_tag)
            tcPr.append(edge_element)
        
        # Set border properties
        if edge in kwargs:
            for key, value in kwargs[edge].items():
                edge_element.set(qn('w:{}'.format(key)), str(value))

# Word formatting with superscript support
def add_italic_text_with_superscript(cell, text):
    """Add italic text with superscript 'a' for duplicates and * for introns"""
    cell.text = ""
    p = cell.add_paragraph()
    gene_names = text.split(", ")
    
    for i, gene_name in enumerate(gene_names):
        if i > 0:
            p.add_run(", ")
        
        # Check for both markers
        has_intron = "*" in gene_name
        has_duplicate = "^a" in gene_name
        
        # Remove markers to get base name
        base_name = gene_name.replace("*", "").replace("^a", "")
        
        # Add base name in italic
        run = p.add_run(base_name)
        run.italic = True
        
        # Add markers based on what's present
        if has_intron and has_duplicate:
            # Both markers: *, a
            asterisk = p.add_run("*")
            asterisk.italic = True
            comma_sup = p.add_run(", ")
            comma_sup.italic = True
            add_superscript(comma_sup, ", ")
            a_sup = p.add_run("a")
            a_sup.italic = True
            add_superscript(a_sup, "a")
        elif has_intron:
            # Only intron marker: *
            asterisk = p.add_run("*")
            asterisk.italic = True
        elif has_duplicate:
            # Only duplicate marker: a (superscript, no comma)
            a_sup = p.add_run("a")
            a_sup.italic = True
            add_superscript(a_sup, "a")

def create_table(df, output_file):
    doc = Document()
    
    # Set document margins for better layout
    sections = doc.sections
    for section in sections:
        section.top_margin = Inches(1)
        section.bottom_margin = Inches(1)
        section.left_margin = Inches(1)
        section.right_margin = Inches(1)
    
    # Add table title
    title = doc.add_paragraph()
    title_run = title.add_run("Table S1. Gene content of the chloroplast genome")
    title_run.bold = True
    title_run.font.size = Pt(12)
    title.alignment = WD_ALIGN_PARAGRAPH.LEFT
    
    # Create table
    table = doc.add_table(rows=1, cols=len(df.columns))
    table.style = 'Table Grid'
    
    # Format header row
    hdr = table.rows[0].cells
    header_labels = ["Category for genes", "Group of genes", "Name of genes", "Gene number"]
    
    for i, label in enumerate(header_labels):
        hdr[i].text = label
        # Bold header text
        for paragraph in hdr[i].paragraphs:
            for run in paragraph.runs:
                run.font.bold = True
                run.font.size = Pt(10)
            paragraph.alignment = WD_ALIGN_PARAGRAPH.CENTER
        # Center align header cells vertically
        tc = hdr[i]._element
        tcPr = tc.get_or_add_tcPr()
        tcVAlign = OxmlElement('w:vAlign')
        tcVAlign.set(qn('w:val'), 'center')
        tcPr.append(tcVAlign)
    
    # Store rows by category for merging
    category_rows = {}
    current_category = None
    
    for idx, row in df.iterrows():
        cells = table.add_row().cells
        row_index = len(table.rows) - 1
        
        category = str(row["Category for genes"])
        
        # Track rows for each category
        if category and category != "":
            if category not in category_rows:
                category_rows[category] = []
            category_rows[category].append(row_index)
            
            # Only write category name in first occurrence
            if category != current_category:
                cells[0].text = category
                current_category = category
            else:
                cells[0].text = ""  # Leave empty for subsequent rows
        else:
            cells[0].text = ""
            current_category = None
        
        cells[1].text = str(row["Group of genes"])
        
        if row["Name of genes"] == "Total number of genes":
            cells[2].text = row["Name of genes"]
            # Bold the total row
            for paragraph in cells[2].paragraphs:
                for run in paragraph.runs:
                    run.font.bold = True
            for paragraph in cells[3].paragraphs:
                for run in paragraph.runs:
                    run.font.bold = True
        else:
            add_italic_text_with_superscript(cells[2], row["Name of genes"])
        
        cells[3].text = str(row["Amount"])
        
        # Set font size for all cells
        for cell in cells:
            for paragraph in cell.paragraphs:
                for run in paragraph.runs:
                    run.font.size = Pt(10)
    
    # Merge cells for each category and center align
    for category, row_indices in category_rows.items():
        if len(row_indices) > 1:
            # Get the first cell
            first_cell = table.rows[row_indices[0]].cells[0]
            
            # Merge with all subsequent cells in this category
            for i in range(1, len(row_indices)):
                first_cell.merge(table.rows[row_indices[i]].cells[0])
            
            # Center align the merged cell
            first_cell.paragraphs[0].alignment = WD_ALIGN_PARAGRAPH.CENTER
            # Also set vertical alignment to center
            tc = first_cell._element
            tcPr = tc.get_or_add_tcPr()
            tcVAlign = OxmlElement('w:vAlign')
            tcVAlign.set(qn('w:val'), 'center')
            tcPr.append(tcVAlign)
    
    # Set column widths for better appearance
    widths = [Inches(1.5), Inches(2.2), Inches(3.5), Inches(0.8)]
    for row in table.rows:
        for idx, width in enumerate(widths):
            row.cells[idx].width = width
    
    # Add notes at the end
    doc.add_paragraph()
    notes = doc.add_paragraph()
    notes_run = notes.add_run("Note: ")
    notes_run.bold = True
    notes_run.font.size = Pt(10)
    
    text_run = notes.add_run("*, ")
    text_run.font.size = Pt(10)
    
    sup_run = notes.add_run("a")
    sup_run.font.size = Pt(10)
    add_superscript(sup_run, "a")
    
    final_text = notes.add_run(" indicate genes containing introns and duplicated genes in inverted repeat (IR) regions, respectively. The ")
    final_text.font.size = Pt(10)
    
    # Add italic rps12
    rps12_run = notes.add_run("rps12")
    rps12_run.font.size = Pt(10)
    rps12_run.italic = True
    
    # Continue with rest of note
    trans_text = notes.add_run(" gene is a trans-spliced gene and is not marked as duplicated despite appearing in multiple locations.")
    trans_text.font.size = Pt(10)

    doc.save(output_file)
    print(f"✓ Saved: {output_file}")

# Process a single GenBank file
def process_genbank_file(genbank_file):
    """Process a single GenBank file and create its Word document"""
    # Extract filename without extension for output naming
    base_name = os.path.splitext(os.path.basename(genbank_file))[0]
    output_file = f"Table_{base_name}.docx"
    
    print(f"\nProcessing: {genbank_file}")
    
    try:
        df, introns = extract_gene_content(genbank_file)
        create_table(df, output_file)
        
        print(f"  CDS with introns: {introns['CDS with introns']}")
        print(f"  tRNA with introns: {introns['tRNA with introns']}")
        
        return True
    except Exception as e:
        print(f"✗ Error processing {genbank_file}: {str(e)}")
        return False

# MAIN
def main():
    # Find all .gb files in the current directory
    gb_files = glob.glob("*.gb")
    
    if not gb_files:
        print("No .gb files found in the current directory!")
        return
    
    print(f"Found {len(gb_files)} GenBank file(s):")
    for f in gb_files:
        print(f"  - {f}")
    
    print("\n" + "="*60)
    
    # Process each file
    success_count = 0
    for gb_file in gb_files:
        if process_genbank_file(gb_file):
            success_count += 1
    
    print("\n" + "="*60)
    print(f"\nSummary: Successfully processed {success_count}/{len(gb_files)} file(s)")

if __name__ == "__main__":
    main()
```
