## Supplementary Date File 1 for "CGAS (Chloroplast Genome Analysis Suite): An Automated Python Pipeline for Comprehensive Comparative Chloroplast Genomics": module3_comparative_analysis.py

```
import os
import pandas as pd
from Bio import SeqIO
from openpyxl import load_workbook
from openpyxl.styles import Font, Alignment, PatternFill, Border, Side
from openpyxl.utils import get_column_letter

# Function to calculate GC content with rounding to two decimal places
def calculate_gc(sequence):
    total_bases = len(sequence)
    gc_count = sequence.count('G') + sequence.count('C')
    return round((gc_count / total_bases) * 100, 2) if total_bases > 0 else 0

# Initialize list to store results
results = []

# Use current working directory
folder_path = os.getcwd()

# Loop through .gb and .gbf files in the folder
for filename in os.listdir(folder_path):
    if filename.endswith('.gb') or filename.endswith('.gbf'):
        record_path = os.path.join(folder_path, filename)
        record = SeqIO.read(record_path, 'genbank')
        
        # Basic genome information
        species = record.annotations['organism']
        genome_length = len(record.seq)
        accession_number = record.annotations.get('accessions', ['N/A'])[0]
        
        # Initialize variables for LSC and SSC regions
        lsc_feature = ssc_feature = None
        
        # Search for LSC and SSC in feature notes
        for feature in record.features:
            if 'note' in feature.qualifiers:
                note_value = feature.qualifiers['note'][0].lower()
                if any(keyword in note_value for keyword in ["large single copy (lsc)", "large single copy region (lsc)", "lsc"]):
                    lsc_feature = feature
                elif any(keyword in note_value for keyword in ["small single copy region (ssc)", "small single copy (ssc)", "ssc"]):
                    ssc_feature = feature
        
        # Calculate exact lengths of LSC and SSC
        lsc_length = len(record.seq[lsc_feature.location.start:lsc_feature.location.end]) if lsc_feature else 0
        ssc_length = len(record.seq[ssc_feature.location.start:ssc_feature.location.end]) if ssc_feature else 0
        
        # Calculate IR length
        ir_length = (genome_length - lsc_length - ssc_length) // 2

        # Calculate GC content
        total_gc = calculate_gc(record.seq)
        lsc_gc = calculate_gc(record.seq[lsc_feature.location.start:lsc_feature.location.end]) if lsc_feature else 0
        ssc_gc = calculate_gc(record.seq[ssc_feature.location.start:ssc_feature.location.end]) if ssc_feature else 0

        # IR = everything except LSC and SSC
        if lsc_feature and ssc_feature:
            ir_seq = (
                record.seq[:lsc_feature.location.start] +
                record.seq[lsc_feature.location.end:ssc_feature.location.start] +
                record.seq[ssc_feature.location.end:]
            )
            ir_gc = calculate_gc(ir_seq)
        else:
            ir_gc = 0
        
        # Extract sequences for tRNA, rRNA, CDS
        tRNA_seq = rRNA_seq = CDS_seq = ""

        for feature in record.features:
            parts = []
            if hasattr(feature.location, "parts"):
                parts = feature.location.parts
            else:
                parts = [feature.location]

            if feature.type == 'CDS':
                for part in parts:
                    CDS_seq += part.extract(record.seq)
            elif feature.type == 'tRNA':
                for part in parts:
                    tRNA_seq += part.extract(record.seq)
            elif feature.type == 'rRNA':
                for part in parts:
                    rRNA_seq += part.extract(record.seq)

        # GC content for gene groups
        tRNA_gc = calculate_gc(tRNA_seq)
        rRNA_gc = calculate_gc(rRNA_seq)
        CDS_gc = calculate_gc(CDS_seq)
        
        # Store results as NUMBERS (not formatted strings)
        results.append([
            species, genome_length, lsc_length, ssc_length, ir_length, 
            total_gc, lsc_gc, ssc_gc, ir_gc, 
            tRNA_gc, rRNA_gc, CDS_gc, accession_number
        ])

# Create output table with new column structure
df = pd.DataFrame(results, columns=[
    "Species", "Complete", "LSC", "SSC", "IR",
    "Complete_GC", "LSC_GC", "SSC_GC", "IR_GC",
    "tRNA", "rRNA", "CDS", "Accession Number"
])

# Save to Excel
output_file = 'genome_analysis_publication_quality.xlsx'
df.to_excel(output_file, index=False, engine='openpyxl')

# Apply publication-quality formatting
wb = load_workbook(output_file)
ws = wb.active

# Define styles
header_font = Font(name='Arial', size=11, bold=True, color='FFFFFF')
subheader_font = Font(name='Arial', size=10, bold=True, color='FFFFFF')
header_fill = PatternFill(start_color='4472C4', end_color='4472C4', fill_type='solid')
header_alignment = Alignment(horizontal='center', vertical='center', wrap_text=True)

cell_font = Font(name='Arial', size=10)
species_font = Font(name='Arial', size=10, italic=True)
cell_alignment = Alignment(horizontal='center', vertical='center')
species_alignment = Alignment(horizontal='left', vertical='center')

thin_border = Border(
    left=Side(style='thin', color='000000'),
    right=Side(style='thin', color='000000'),
    top=Side(style='thin', color='000000'),
    bottom=Side(style='thin', color='000000')
)

# Insert a new row at the top for main headers
ws.insert_rows(1)

# Merge cells for main headers
ws.merge_cells('A1:A2')  # Species
ws.merge_cells('B1:E1')  # Genome Length (bp)
ws.merge_cells('F1:I1')  # GC (%)
ws.merge_cells('J1:L1')  # GC (%)
ws.merge_cells('M1:M2')  # Accession Number

# Set main headers (row 1)
ws['A1'] = 'Species'
ws['B1'] = 'Genome Length (bp)'
ws['F1'] = 'GC (%)'
ws['J1'] = 'GC (%)'
ws['M1'] = 'Accession Number'

# Set subheaders (row 2)
ws['B2'] = 'Complete'
ws['C2'] = 'LSC'
ws['D2'] = 'SSC'
ws['E2'] = 'IR'
ws['F2'] = 'Complete'
ws['G2'] = 'LSC'
ws['H2'] = 'SSC'
ws['I2'] = 'IR'
ws['J2'] = 'tRNA'
ws['K2'] = 'rRNA'
ws['L2'] = 'CDS'

# Format all header cells
for row in [1, 2]:
    for col in range(1, 14):
        cell = ws.cell(row=row, column=col)
        cell.font = header_font if row == 1 else subheader_font
        cell.fill = header_fill
        cell.alignment = header_alignment
        cell.border = thin_border

# Format data cells (starting from row 3)
for row_num in range(3, len(df) + 3):
    for col_num in range(1, 14):
        cell = ws.cell(row=row_num, column=col_num)
        cell.border = thin_border
        
        # Species column - italic and left-aligned
        if col_num == 1:
            cell.font = species_font
            cell.alignment = species_alignment
        # Genome length columns (B-E: Complete, LSC, SSC, IR)
        elif col_num in [2, 3, 4, 5]:
            cell.font = cell_font
            cell.alignment = Alignment(horizontal='right', vertical='center')
            cell.number_format = '#,##0'
        # GC percentage columns (F-L)
        elif col_num in [6, 7, 8, 9, 10, 11, 12]:
            cell.font = cell_font
            cell.alignment = cell_alignment
            cell.number_format = '0.00'
        # Accession number
        else:
            cell.font = cell_font
            cell.alignment = cell_alignment

# Adjust column widths
column_widths = {
    'A': 35,  # Species
    'B': 12,  # Complete
    'C': 12,  # LSC
    'D': 12,  # SSC
    'E': 12,  # IR
    'F': 12,  # Complete GC
    'G': 12,  # LSC GC
    'H': 12,  # SSC GC
    'I': 12,  # IR GC
    'J': 12,  # tRNA GC
    'K': 12,  # rRNA GC
    'L': 12,  # CDS GC
    'M': 16   # Accession Number
}

for col, width in column_widths.items():
    ws.column_dimensions[col].width = width

# Set row heights
ws.row_dimensions[1].height = 25
ws.row_dimensions[2].height = 25

# Freeze the header rows
ws.freeze_panes = 'A3'

# Add footnote
footnote_row = len(df) + 4
ws.merge_cells(f'A{footnote_row}:M{footnote_row}')
footnote_cell = ws.cell(row=footnote_row, column=1)
footnote_cell.value = ('GC: guanine-cytosine content; LSC: large single-copy region; SSC: small single-copy region; '
                       'IR: inverted repeat region; Complete: complete chloroplast genome; '
                       'tRNA: transfer RNA; rRNA: ribosomal RNA; CDS: protein-coding sequences.')
footnote_cell.font = Font(name='Arial', size=9, italic=True)
footnote_cell.alignment = Alignment(horizontal='left', vertical='top', wrap_text=True)
ws.row_dimensions[footnote_row].height = 30

# Save formatted workbook
wb.save(output_file)

print(f"✓ Publication-quality table created: {output_file}")
print(f"✓ Location: {folder_path}")
print(f"✓ Total genomes analyzed: {len(results)}")
print(f"✓ Format: Two-level headers with grouped columns")
print(f"✓ Footnote added with abbreviation definitions")
```
