## Supplementary Date File 1 for "CGAS (Chloroplast Genome Analysis Suite): An Automated Python Pipeline for Comprehensive Comparative Chloroplast Genomics": module4_codon_usage.py

```
#!/usr/bin/env python3
"""
Relative Synonymous Codon Usage (RSCU) Analysis Pipeline
=========================================================

This script performs comprehensive codon usage analysis on GenBank files by:
1. Extracting coding sequences (CDS) from GenBank format files
2. Calculating Relative Synonymous Codon Usage (RSCU) values
3. Generating individual RSCU reports for each genome
4. Creating a merged comparative analysis across all genomes

Author: Bioinformatics Analysis Tool
Version: 1.0
Date: 2025

Dependencies:
    - biopython (Bio)
    - pandas
    - openpyxl (for Excel file handling)

Usage:
    Place GenBank files (.gb, .gbf, .gbk) in the working directory and run:
    python codon_usage_analysis.py

Output:
    - Individual RSCU files: [genome_name]_RSCU.xlsx
    - Merged analysis: Merged_RSCU_Analysis.xlsx
"""

import os
import sys
from pathlib import Path
from typing import List, Dict, Tuple
from collections import Counter, defaultdict

try:
    from Bio import SeqIO
    import pandas as pd
except ImportError as e:
    print(f"Error: Required package not installed: {e}")
    print("Please install required packages using:")
    print("pip install biopython pandas openpyxl")
    sys.exit(1)


# ============================================================================
# CODON USAGE TABLE
# ============================================================================
# Standard genetic code: mapping of amino acids to their synonymous codons
# Based on the universal genetic code (NCBI transl_table=1)

SYNONYMOUS_CODONS = {
    'Ala': ['GCT', 'GCC', 'GCA', 'GCG'],
    'Arg': ['CGT', 'CGC', 'CGA', 'CGG', 'AGA', 'AGG'],
    'Asn': ['AAT', 'AAC'],
    'Asp': ['GAT', 'GAC'],
    'Cys': ['TGT', 'TGC'],
    'Gln': ['CAA', 'CAG'],
    'Glu': ['GAA', 'GAG'],
    'Gly': ['GGT', 'GGC', 'GGA', 'GGG'],
    'His': ['CAT', 'CAC'],
    'Ile': ['ATT', 'ATC', 'ATA'],
    'Leu': ['TTA', 'TTG', 'CTT', 'CTC', 'CTA', 'CTG'],
    'Lys': ['AAA', 'AAG'],
    'Met': ['ATG'],  # Start codon (non-degenerate)
    'Phe': ['TTT', 'TTC'],
    'Pro': ['CCT', 'CCC', 'CCA', 'CCG'],
    'Ser': ['TCT', 'TCC', 'TCA', 'TCG', 'AGT', 'AGC'],
    'Thr': ['ACT', 'ACC', 'ACA', 'ACG'],
    'Trp': ['TGG'],  # Non-degenerate
    'Tyr': ['TAT', 'TAC'],
    'Val': ['GTT', 'GTC', 'GTA', 'GTG'],
    'Stop': ['TAA', 'TAG', 'TGA']
}

# Full amino acid names for publication-quality output
AMINO_ACID_NAMES = {
    'Ala': 'Alanine',
    'Arg': 'Arginine',
    'Asn': 'Asparagine',
    'Asp': 'Aspartate',
    'Cys': 'Cysteine',
    'Gln': 'Glutamine',
    'Glu': 'Glutamate',
    'Gly': 'Glycine',
    'His': 'Histidine',
    'Ile': 'Isoleucine',
    'Leu': 'Leucine',
    'Lys': 'Lysine',
    'Met': 'Methionine',
    'Phe': 'Phenylalanine',
    'Pro': 'Proline',
    'Ser': 'Serine',
    'Thr': 'Threonine',
    'Trp': 'Tryptophan',
    'Tyr': 'Tyrosine',
    'Val': 'Valine',
    'Stop': 'Stop'
}

# Define the order of amino acids for publication (alphabetical by 3-letter code)
AA_ORDER = ['Ala', 'Arg', 'Asn', 'Asp', 'Cys', 'Gln', 'Glu', 'Gly', 'His', 
            'Ile', 'Leu', 'Lys', 'Met', 'Phe', 'Pro', 'Ser', 'Thr', 'Trp', 
            'Tyr', 'Val', 'Stop']


# ============================================================================
# CODON EXTRACTION AND COUNTING
# ============================================================================

def extract_coding_sequences(genbank_file: str) -> List[str]:
    """
    Extract all coding sequences (CDS) from a GenBank file.
    
    Parameters:
    -----------
    genbank_file : str
        Path to the GenBank format file
        
    Returns:
    --------
    List[str]
        List of coding sequences as uppercase strings
        
    Raises:
    -------
    FileNotFoundError
        If the GenBank file doesn't exist
    ValueError
        If the file cannot be parsed as GenBank format
    """
    try:
        genome_record = SeqIO.read(genbank_file, "genbank")
    except FileNotFoundError:
        raise FileNotFoundError(f"GenBank file not found: {genbank_file}")
    except Exception as e:
        raise ValueError(f"Error parsing GenBank file {genbank_file}: {e}")
    
    coding_sequences = []
    cds_count = 0
    
    # Iterate through all features in the genome
    for feature in genome_record.features:
        if feature.type == "CDS":
            cds_count += 1
            # Extract the sequence and remove any whitespace
            seq = str(feature.extract(genome_record.seq))
            seq = seq.replace("\n", "").replace(" ", "").upper()
            coding_sequences.append(seq)
    
    print(f"  - Extracted {cds_count} coding sequences")
    return coding_sequences


def count_codons(coding_sequences: List[str]) -> Tuple[Counter, List[str]]:
    """
    Count the occurrence of each codon across all coding sequences.
    
    Parameters:
    -----------
    coding_sequences : List[str]
        List of coding sequences
        
    Returns:
    --------
    Tuple[Counter, List[str]]
        Tuple of (codon counts, list of unusual/ambiguous codons found)
        
    Notes:
    ------
    - Only complete codons (3 nucleotides) are counted
    - Partial codons at sequence ends are ignored
    - Detects ambiguous nucleotides (N, R, Y, etc.)
    - Detects non-standard nucleotides
    """
    # Valid nucleotides in standard genetic code
    valid_nucleotides = set('ATCG')
    
    # Concatenate all CDS into a single sequence
    all_cds_sequence = "".join(coding_sequences)
    
    # Count codons in triplets and detect unusual ones
    codon_counts = Counter()
    unusual_codons = []
    
    for i in range(0, len(all_cds_sequence), 3):
        codon = all_cds_sequence[i:i+3]
        
        # Ensure complete codon
        if len(codon) == 3:
            codon_counts[codon] += 1
            
            # Check for unusual nucleotides
            codon_nucleotides = set(codon)
            if not codon_nucleotides.issubset(valid_nucleotides):
                if codon not in unusual_codons:
                    unusual_codons.append(codon)
    
    total_codons = sum(codon_counts.values())
    print(f"  - Total codons counted: {total_codons:,}")
    
    if unusual_codons:
        print(f"  ⚠ WARNING: Found {len(unusual_codons)} unusual codon(s): {', '.join(unusual_codons[:10])}")
        if len(unusual_codons) > 10:
            print(f"           (and {len(unusual_codons) - 10} more...)")
    
    return codon_counts, unusual_codons


# ============================================================================
# RSCU CALCULATION
# ============================================================================

def calculate_rscu(codon_counts: Counter, 
                   synonymous_codons: Dict[str, List[str]],
                   unusual_codons: List[str] = None) -> Tuple[pd.DataFrame, pd.DataFrame]:
    """
    Calculate Relative Synonymous Codon Usage (RSCU) values.
    
    RSCU is defined as the ratio of the observed frequency of a codon to the
    expected frequency if all synonymous codons for an amino acid were used
    equally.
    
    RSCU = (Observed codon count) / (Expected codon count)
    
    Where:
        Expected count = (Total AA count) / (Number of synonymous codons)
    
    RSCU values interpretation:
        - RSCU > 1.0: Codon is used more frequently than expected
        - RSCU = 1.0: Codon is used at the expected frequency
        - RSCU < 1.0: Codon is used less frequently than expected
    
    Parameters:
    -----------
    codon_counts : Counter
        Dictionary of codon counts
    synonymous_codons : Dict[str, List[str]]
        Mapping of amino acids to their synonymous codons
    unusual_codons : List[str], optional
        List of unusual/ambiguous codons detected
        
    Returns:
    --------
    Tuple[pd.DataFrame, pd.DataFrame]
        Tuple of (RSCU DataFrame, unusual codons DataFrame)
    """
    # Get all standard codons
    all_standard_codons = set()
    for codons in synonymous_codons.values():
        all_standard_codons.update(codons)
    
    # Calculate total counts for each amino acid
    aa_totals = defaultdict(int)
    for aa, codons in synonymous_codons.items():
        for codon in codons:
            aa_totals[aa] += codon_counts[codon]
    
    # Calculate RSCU for each codon in publication order
    rscu_data = []
    for aa in AA_ORDER:
        codons = synonymous_codons[aa]
        num_synonymous = len(codons)  # Degeneracy of amino acid
        full_name = AMINO_ACID_NAMES[aa]
        
        for idx, codon in enumerate(codons):
            observed_count = codon_counts[codon]
            
            # Calculate expected count (uniform distribution)
            expected_count = aa_totals[aa] / num_synonymous if aa_totals[aa] > 0 else 0
            
            # Calculate RSCU
            if expected_count > 0:
                rscu = observed_count / expected_count
            else:
                rscu = 0.0
            
            # For publication format: show AA name only for first codon
            if idx == 0:
                rscu_data.append({
                    'AA': aa,
                    'Amino_Acid': full_name,
                    'Codon': codon,
                    'Count': observed_count,
                    'RSCU': round(rscu, 4)
                })
            else:
                rscu_data.append({
                    'AA': '',  # Empty for subsequent codons
                    'Amino_Acid': '',
                    'Codon': codon,
                    'Count': observed_count,
                    'RSCU': round(rscu, 4)
                })
    
    # Create DataFrame
    df_rscu = pd.DataFrame(rscu_data)
    
    # Handle unusual codons (those not in standard genetic code)
    unusual_data = []
    if unusual_codons:
        for codon in unusual_codons:
            if codon not in all_standard_codons:
                unusual_data.append({
                    'Codon': codon,
                    'Count': codon_counts[codon],
                    'Note': 'Contains ambiguous/non-standard nucleotides'
                })
    
    # Also check for observed codons not in standard list
    for codon, count in codon_counts.items():
        if codon not in all_standard_codons and codon not in [u['Codon'] for u in unusual_data]:
            unusual_data.append({
                'Codon': codon,
                'Count': count,
                'Note': 'Not in standard genetic code table'
            })
    
    df_unusual = pd.DataFrame(unusual_data) if unusual_data else pd.DataFrame()
    
    if not df_unusual.empty:
        total_unusual = df_unusual['Count'].sum()
        print(f"  ⚠ WARNING: {len(df_unusual)} unusual codon type(s) with {total_unusual:,} total occurrences")
    
    return df_rscu, df_unusual


# ============================================================================
# FILE PROCESSING
# ============================================================================

def process_single_genbank_file(file_path: str) -> Tuple[str, pd.DataFrame]:
    """
    Process a single GenBank file and calculate RSCU values.
    
    Parameters:
    -----------
    file_path : str
        Path to the GenBank file
        
    Returns:
    --------
    Tuple[str, pd.DataFrame]
        Tuple of (output filename, RSCU DataFrame)
    """
    print(f"\nProcessing: {file_path}")
    
    # Extract coding sequences
    coding_sequences = extract_coding_sequences(file_path)
    
    if not coding_sequences:
        print(f"  WARNING: No coding sequences found in {file_path}")
        return None, None
    
    # Count codons and detect unusual ones
    codon_counts, unusual_codons = count_codons(coding_sequences)
    
    # Calculate RSCU
    df_rscu, df_unusual = calculate_rscu(codon_counts, SYNONYMOUS_CODONS, unusual_codons)
    
    # Generate output filename
    base_name = Path(file_path).stem
    output_file = f"{base_name}_RSCU.xlsx"
    
    # Save individual RSCU file with multiple sheets
    with pd.ExcelWriter(output_file, engine='openpyxl') as writer:
        # Main RSCU analysis (all 64 standard codons)
        df_rscu.to_excel(writer, sheet_name='RSCU_Analysis', index=False)
        
        # If unusual codons found, add them to a separate sheet
        if not df_unusual.empty:
            df_unusual.to_excel(writer, sheet_name='Unusual_Codons', index=False)
            print(f"  ⚠ Unusual codons saved in separate sheet")
        
        # Add summary statistics sheet
        summary_data = {
            'Metric': [
                'Total CDS sequences',
                'Total codons analyzed',
                'Standard codons',
                'Unusual codons detected',
                'Unusual codon occurrences'
            ],
            'Value': [
                len(coding_sequences),
                sum(codon_counts.values()),
                64,
                len(df_unusual),
                df_unusual['Count'].sum() if not df_unusual.empty else 0
            ]
        }
        df_summary = pd.DataFrame(summary_data)
        df_summary.to_excel(writer, sheet_name='Summary', index=False)
    
    print(f"  ✓ Saved: {output_file}")
    
    return output_file, df_rscu


def find_genbank_files(directory: str = None) -> List[str]:
    """
    Find all GenBank files in the specified directory.
    
    Parameters:
    -----------
    directory : str, optional
        Directory to search (default: current working directory)
        
    Returns:
    --------
    List[str]
        List of GenBank file paths
    """
    if directory is None:
        directory = os.getcwd()
    
    # Supported GenBank file extensions
    genbank_extensions = ('.gb', '.gbf', '.gbk', '.genbank')
    
    genbank_files = [
        os.path.join(directory, f) 
        for f in os.listdir(directory) 
        if f.lower().endswith(genbank_extensions)
    ]
    
    return sorted(genbank_files)


# ============================================================================
# MERGE RSCU FILES
# ============================================================================

def merge_rscu_files(rscu_files: List[str], output_file: str = 'Merged_RSCU_Analysis.xlsx'):
    """
    Merge individual RSCU files into a single comparative analysis file.
    
    Creates a merged Excel file where each column represents RSCU values from
    a different genome, allowing for easy comparison of codon usage patterns.
    The amino acid cells are vertically merged to span all synonymous codons.
    
    Parameters:
    -----------
    rscu_files : List[str]
        List of RSCU Excel file paths to merge
    output_file : str
        Name of the output merged file
    """
    print(f"\n{'='*70}")
    print("MERGING RSCU FILES")
    print(f"{'='*70}")
    
    if not rscu_files:
        print("No RSCU files to merge.")
        return
    
    # Read all RSCU files and collect data
    all_data = []
    genome_names = []
    
    for filename in rscu_files:
        print(f"  - Adding: {filename}")
        
        try:
            # Read the RSCU file
            data = pd.read_excel(filename, sheet_name='RSCU_Analysis')
            
            # Get the genome name from filename
            genome_name = Path(filename).stem.replace('_RSCU', '')
            genome_names.append(genome_name)
            
            # Store the data
            all_data.append(data)
            
        except Exception as e:
            print(f"  WARNING: Error reading {filename}: {e}")
            continue
    
    if not all_data:
        print("  ERROR: No data to merge.")
        return
    
    # Create the merged structure
    # Start with Codon column from first file
    base_data = all_data[0][['AA', 'Amino_Acid', 'Codon']].copy()
    
    # Keep amino acid name only on first occurrence
    display_amino_acid = []
    for idx, row in base_data.iterrows():
        if row['AA']:  # Non-empty AA code means first codon of new amino acid
            display_amino_acid.append(row['Amino_Acid'])
        else:
            display_amino_acid.append('')
    
    # Build the final merged dataframe
    merged_data = pd.DataFrame()
    merged_data['Amino_Acid'] = display_amino_acid
    merged_data['Codon'] = base_data['Codon']
    
    # Add RSCU columns from each genome
    for genome_name, data in zip(genome_names, all_data):
        merged_data[genome_name] = data['RSCU'].values
    
    # Save merged file with proper formatting
    with pd.ExcelWriter(output_file, engine='openpyxl') as writer:
        merged_data.to_excel(writer, sheet_name='Merged_RSCU', index=False)
        
        # Get the worksheet to apply formatting
        workbook = writer.book
        worksheet = writer.sheets['Merged_RSCU']
        
        # Apply formatting for publication quality
        from openpyxl.styles import Font, Alignment, PatternFill, Border, Side
        
        # Header formatting
        header_fill = PatternFill(start_color='366092', end_color='366092', fill_type='solid')
        header_font = Font(bold=True, color='FFFFFF', size=11)
        
        for cell in worksheet[1]:
            cell.fill = header_fill
            cell.font = header_font
            cell.alignment = Alignment(horizontal='center', vertical='center')
        
        # Make species name headers italic (columns C onwards, excluding "Amino_Acid" and "Codon")
        for col_idx in range(3, len(genome_names) + 3):
            header_cell = worksheet.cell(row=1, column=col_idx)
            header_cell.font = Font(bold=True, italic=True, color='FFFFFF', size=11)
        
        # Data formatting
        thin_border = Border(
            left=Side(style='thin'),
            right=Side(style='thin'),
            top=Side(style='thin'),
            bottom=Side(style='thin')
        )
        
        # Track amino acid groups for merging cells
        merge_groups = []
        current_aa = None
        start_row = None
        
        # First pass: identify merge ranges and format cells
        
        # Format data rows
        for row_idx, row in enumerate(worksheet.iter_rows(min_row=2, max_row=worksheet.max_row), start=2):
            amino_acid_value = row[0].value
            
            # Track amino acid groups for cell merging
            if amino_acid_value and str(amino_acid_value).strip():
                # Save previous group if exists
                if current_aa is not None and start_row is not None:
                    end_row = row_idx - 1
                    if end_row > start_row:  # Only merge if more than one row
                        merge_groups.append((start_row, end_row))
                
                # Start new group
                current_aa = amino_acid_value
                start_row = row_idx
            
            # Format all cells in the row
            for col_idx, cell in enumerate(row):
                cell.border = thin_border
                
                # First column (Amino_Acid): left align and make bold + italic
                if col_idx == 0:
                    cell.alignment = Alignment(horizontal='center', vertical='center')
                    if cell.value and str(cell.value).strip():
                        cell.font = Font(bold=True, italic=True, size=10)
                
                # Second column (Codon): center align
                elif col_idx == 1:
                    cell.alignment = Alignment(horizontal='center', vertical='center')
                
                # RSCU value columns: format to 4 decimal places (NO italic for numbers)
                else:
                    cell.alignment = Alignment(horizontal='center', vertical='center')
                    if cell.value is not None:
                        try:
                            cell.number_format = '0.0000'
                            # Don't apply italic to RSCU numbers
                        except:
                            pass
        
        # Don't forget the last group
        if current_aa is not None and start_row is not None:
            end_row = worksheet.max_row
            if end_row > start_row:  # Only merge if more than one row
                merge_groups.append((start_row, end_row))
        
        # Second pass: merge cells for amino acids with multiple codons
        for start_row, end_row in merge_groups:
            # Merge cells in the Amino_Acid column (column A)
            worksheet.merge_cells(f'A{start_row}:A{end_row}')
            
            # Apply formatting to merged cell
            merged_cell = worksheet[f'A{start_row}']
            merged_cell.alignment = Alignment(horizontal='center', vertical='center')
            merged_cell.font = Font(bold=True, italic=True, size=10)
            merged_cell.border = thin_border
        
        # Adjust column widths
        worksheet.column_dimensions['A'].width = 15  # Amino_Acid
        worksheet.column_dimensions['B'].width = 8   # Codon
        
        # Set width for genome columns (make them wider for species names)
        for col_idx in range(3, len(genome_names) + 3):
            col_letter = worksheet.cell(row=1, column=col_idx).column_letter
            worksheet.column_dimensions[col_letter].width = 14
        
        # Freeze panes (freeze first row and first two columns)
        worksheet.freeze_panes = 'C2'
    
    print(f"\n  ✓ Merged analysis saved: {output_file}")
    print(f"  - Total genomes compared: {len(genome_names)}")
    print(f"  - Total codons: {len(merged_data)}")
    print(f"  - Format: Publication-ready with vertically merged amino acid cells")
    print(f"  - Species names formatted in italic")


# ============================================================================
# MAIN PIPELINE
# ============================================================================

def main():
    """
    Main pipeline execution function.
    
    Workflow:
    1. Find all GenBank files in working directory
    2. Process each file to calculate RSCU
    3. Merge all RSCU files into comparative analysis
    """
    print(f"\n{'='*70}")
    print("CODON USAGE ANALYSIS PIPELINE")
    print(f"{'='*70}")
    
    # Get working directory
    working_dir = os.getcwd()
    print(f"\nWorking directory: {working_dir}")
    
    # Find GenBank files
    genbank_files = find_genbank_files(working_dir)
    
    if not genbank_files:
        print("\n❌ ERROR: No GenBank files found!")
        print("Please place GenBank files (.gb, .gbf, .gbk) in the working directory.")
        return
    
    print(f"\nFound {len(genbank_files)} GenBank file(s):")
    for gf in genbank_files:
        print(f"  - {os.path.basename(gf)}")
    
    # Process each GenBank file
    print(f"\n{'='*70}")
    print("CALCULATING RSCU VALUES")
    print(f"{'='*70}")
    
    rscu_files = []
    for gb_file in genbank_files:
        try:
            output_file, _ = process_single_genbank_file(gb_file)
            if output_file:
                rscu_files.append(output_file)
        except Exception as e:
            print(f"  ❌ ERROR processing {gb_file}: {e}")
            continue
    
    # Merge RSCU files
    if rscu_files:
        merge_rscu_files(rscu_files)
    
    # Summary
    print(f"\n{'='*70}")
    print("ANALYSIS COMPLETE")
    print(f"{'='*70}")
    print(f"✓ Successfully processed: {len(rscu_files)} genome(s)")
    print(f"✓ Individual RSCU files: {len(rscu_files)}")
    print(f"✓ Merged analysis file: Merged_RSCU_Analysis.xlsx")
    print(f"\nAll output files saved in: {working_dir}")
    print(f"{'='*70}\n")


# ============================================================================
# SCRIPT ENTRY POINT
# ============================================================================

if __name__ == "__main__":
    try:
        main()
    except KeyboardInterrupt:
        print("\n\n⚠ Analysis interrupted by user.")
        sys.exit(0)
    except Exception as e:
        print(f"\n\n❌ FATAL ERROR: {e}")
        import traceback
        traceback.print_exc()
        sys.exit(1)
```
