## Supplementary Date File 1 for "CGAS (Chloroplast Genome Analysis Suite): An Automated Python Pipeline for Comprehensive Comparative Chloroplast Genomics": module5_amino_acid.py

```
#!/usr/bin/env python3
"""
Amino Acid Composition Analysis Pipeline
=========================================

This script performs comprehensive amino acid composition analysis on GenBank files by:
1. Extracting and translating coding sequences (CDS) from GenBank format files
2. Calculating amino acid composition percentages
3. Generating individual composition reports for each genome
4. Creating a merged comparative analysis across all genomes

Author: Bioinformatics Analysis Tool
Version: 1.0
Date: 2025

Dependencies:
    - biopython (Bio)
    - pandas
    - openpyxl (for Excel file handling)

Usage:
    Place GenBank files (.gb, .gbf, .gbk) in the working directory and run:
    python amino_acid_analysis.py

Output:
    - Individual composition files: [genome_name]_AminoAcid.xlsx
    - Merged analysis: Merged_AminoAcid_Analysis.xlsx
"""

import os
import sys
from pathlib import Path
from typing import List, Dict, Tuple
from collections import Counter

try:
    from Bio import SeqIO
    from Bio.Seq import Seq
    from Bio.SeqUtils import ProtParam
    import pandas as pd
except ImportError as e:
    print(f"Error: Required package not installed: {e}")
    print("Please install required packages using:")
    print("pip install biopython pandas openpyxl")
    sys.exit(1)


# ============================================================================
# AMINO ACID INFORMATION
# ============================================================================

# Full amino acid names mapping
AMINO_ACID_NAMES = {
    'A': 'Alanine',
    'R': 'Arginine',
    'N': 'Asparagine',
    'D': 'Aspartate',
    'C': 'Cysteine',
    'Q': 'Glutamine',
    'E': 'Glutamate',
    'G': 'Glycine',
    'H': 'Histidine',
    'I': 'Isoleucine',
    'L': 'Leucine',
    'K': 'Lysine',
    'M': 'Methionine',
    'F': 'Phenylalanine',
    'P': 'Proline',
    'S': 'Serine',
    'T': 'Threonine',
    'W': 'Tryptophan',
    'Y': 'Tyrosine',
    'V': 'Valine'
}

# Amino acid order for publication (alphabetical)
AA_ORDER = ['A', 'R', 'N', 'D', 'C', 'Q', 'E', 'G', 'H', 'I', 
            'L', 'K', 'M', 'F', 'P', 'S', 'T', 'W', 'Y', 'V']


# ============================================================================
# SEQUENCE EXTRACTION AND TRANSLATION
# ============================================================================

def get_organism_name(genbank_file: str) -> str:
    """
    Extract organism name from GenBank file annotations.
    
    Parameters:
    -----------
    genbank_file : str
        Path to the GenBank file
        
    Returns:
    --------
    str
        Organism name with spaces replaced by underscores
    """
    try:
        record = SeqIO.read(genbank_file, "genbank")
        organism = record.annotations.get("organism", "Unknown")
        # Replace spaces with underscores for file naming
        return organism.replace(" ", "_")
    except Exception as e:
        print(f"  WARNING: Error reading organism name: {e}")
        return "Unknown"


def extract_and_translate_cds(genbank_file: str) -> Tuple[str, int]:
    """
    Extract all CDS features and translate them to amino acid sequences.
    
    Parameters:
    -----------
    genbank_file : str
        Path to the GenBank file
        
    Returns:
    --------
    Tuple[str, int]
        Combined amino acid sequence and number of CDS features processed
        
    Notes:
    ------
    - Sequences are translated using the standard genetic code
    - Translation stops at the first stop codon
    - Stop codons are excluded from the final sequence
    """
    all_amino_acid_sequences = []
    cds_count = 0
    
    try:
        record = SeqIO.read(genbank_file, "genbank")
        
        for feature in record.features:
            if feature.type == "CDS":
                try:
                    # Extract CDS sequence
                    cds_seq = feature.extract(record.seq)
                    
                    # Translate to amino acids (stop at first stop codon)
                    amino_acid_sequence = cds_seq.translate(to_stop=True)
                    
                    all_amino_acid_sequences.append(str(amino_acid_sequence))
                    cds_count += 1
                    
                except Exception as e:
                    print(f"  WARNING: Error translating CDS: {e}")
                    continue
        
        print(f"  - Extracted and translated {cds_count} coding sequences")
        
    except Exception as e:
        print(f"  ERROR: Failed to process file: {e}")
        return "", 0
    
    # Combine all amino acid sequences
    combined_sequence = "".join(all_amino_acid_sequences)
    
    return combined_sequence, cds_count


# ============================================================================
# AMINO ACID COMPOSITION ANALYSIS
# ============================================================================

def analyze_amino_acid_composition(amino_acid_sequence: str) -> pd.DataFrame:
    """
    Analyze amino acid composition and calculate percentages.
    
    Uses Biopython's ProteinAnalysis module to calculate the percentage
    composition of each amino acid in the protein sequence.
    
    Parameters:
    -----------
    amino_acid_sequence : str
        Combined amino acid sequence from all CDS
        
    Returns:
    --------
    pd.DataFrame
        DataFrame with columns: AA_Code, Amino_Acid, Percentage
        
    Notes:
    ------
    - Percentages represent the fraction of each amino acid
    - Values are multiplied by 100 for percentage format
    - Results are sorted alphabetically by amino acid code
    """
    if not amino_acid_sequence:
        print("  WARNING: Empty amino acid sequence")
        return pd.DataFrame()
    
    # Use Biopython's ProteinAnalysis
    protein_analysis = ProtParam.ProteinAnalysis(amino_acid_sequence)
    composition = protein_analysis.amino_acids_percent
    
    # Convert to structured data
    composition_data = []
    for aa_code in AA_ORDER:
        if aa_code in composition:
            percentage = composition[aa_code]  # Already in percentage (0-100)
            composition_data.append({
                'AA_Code': aa_code,
                'Amino_Acid': AMINO_ACID_NAMES[aa_code],
                'Percentage': round(percentage, 4)
            })
    
    df = pd.DataFrame(composition_data)
    
    total_aa = len(amino_acid_sequence)
    print(f"  - Total amino acids analyzed: {total_aa:,}")
    
    return df


# ============================================================================
# FILE PROCESSING
# ============================================================================

def process_single_genbank_file(file_path: str) -> Tuple[str, pd.DataFrame]:
    """
    Process a single GenBank file and calculate amino acid composition.
    
    Parameters:
    -----------
    file_path : str
        Path to the GenBank file
        
    Returns:
    --------
    Tuple[str, pd.DataFrame]
        Tuple of (output filename, composition DataFrame)
    """
    print(f"\nProcessing: {file_path}")
    
    # Get organism name
    organism_name = get_organism_name(file_path)
    print(f"  - Organism: {organism_name}")
    
    # Extract and translate CDS
    amino_acid_sequence, cds_count = extract_and_translate_cds(file_path)
    
    if not amino_acid_sequence:
        print(f"  WARNING: No amino acid sequences found in {file_path}")
        return None, None
    
    # Analyze composition
    df_composition = analyze_amino_acid_composition(amino_acid_sequence)
    
    if df_composition.empty:
        return None, None
    
    # Generate output filename
    base_name = Path(file_path).stem
    output_file = f"{base_name}_AminoAcid.xlsx"
    
    # Save individual composition file
    with pd.ExcelWriter(output_file, engine='openpyxl') as writer:
        # Main composition sheet
        df_composition.to_excel(writer, sheet_name='AA_Composition', index=False)
        
        # Summary statistics sheet
        summary_data = {
            'Metric': [
                'Organism',
                'Total CDS sequences',
                'Total amino acids',
                'Number of AA types'
            ],
            'Value': [
                organism_name.replace('_', ' '),
                cds_count,
                len(amino_acid_sequence),
                len(df_composition)
            ]
        }
        df_summary = pd.DataFrame(summary_data)
        df_summary.to_excel(writer, sheet_name='Summary', index=False)
    
    print(f"  ✓ Saved: {output_file}")
    
    return output_file, df_composition


def find_genbank_files(directory: str = None) -> List[str]:
    """
    Find all GenBank files in the specified directory.
    
    Parameters:
    -----------
    directory : str, optional
        Directory to search (default: current working directory)
        
    Returns:
    --------
    List[str]
        List of GenBank file paths
    """
    if directory is None:
        directory = os.getcwd()
    
    # Supported GenBank file extensions
    genbank_extensions = ('.gb', '.gbf', '.gbk', '.genbank')
    
    genbank_files = [
        os.path.join(directory, f) 
        for f in os.listdir(directory) 
        if f.lower().endswith(genbank_extensions)
    ]
    
    return sorted(genbank_files)


# ============================================================================
# MERGE COMPOSITION FILES
# ============================================================================

def merge_amino_acid_files(aa_files: List[str], output_file: str = 'Merged_AminoAcid_Analysis.xlsx'):
    """
    Merge individual amino acid composition files into a single comparative analysis file.
    
    Creates a merged Excel file where each column represents composition percentages
    from a different genome, with publication-quality formatting.
    
    Parameters:
    -----------
    aa_files : List[str]
        List of amino acid composition Excel file paths to merge
    output_file : str
        Name of the output merged file
    """
    print(f"\n{'='*70}")
    print("MERGING AMINO ACID COMPOSITION FILES")
    print(f"{'='*70}")
    
    if not aa_files:
        print("No amino acid files to merge.")
        return
    
    # Read all composition files and collect data
    all_data = []
    genome_names = []
    
    for filename in aa_files:
        print(f"  - Adding: {filename}")
        
        try:
            # Read the composition file
            data = pd.read_excel(filename, sheet_name='AA_Composition')
            
            # Get the genome name from filename
            genome_name = Path(filename).stem.replace('_AminoAcid', '')
            genome_names.append(genome_name)
            
            # Store the data
            all_data.append(data)
            
        except Exception as e:
            print(f"  WARNING: Error reading {filename}: {e}")
            continue
    
    if not all_data:
        print("  ERROR: No data to merge.")
        return
    
    # Create the merged structure
    base_data = all_data[0][['AA_Code', 'Amino_Acid']].copy()
    
    # Build the final merged dataframe
    merged_data = pd.DataFrame()
    merged_data['Amino_Acid'] = base_data['Amino_Acid']
    
    # Add percentage columns from each genome
    for genome_name, data in zip(genome_names, all_data):
        merged_data[genome_name] = data['Percentage'].values
    
    # Save merged file with proper formatting
    with pd.ExcelWriter(output_file, engine='openpyxl') as writer:
        merged_data.to_excel(writer, sheet_name='Merged_AA_Composition', index=False)
        
        # Get the worksheet to apply formatting
        workbook = writer.book
        worksheet = writer.sheets['Merged_AA_Composition']
        
        # Apply formatting for publication quality
        from openpyxl.styles import Font, Alignment, PatternFill, Border, Side
        
        # Header formatting
        header_fill = PatternFill(start_color='366092', end_color='366092', fill_type='solid')
        
        for cell in worksheet[1]:
            cell.fill = header_fill
            cell.font = Font(bold=True, color='FFFFFF', size=11)
            cell.alignment = Alignment(horizontal='center', vertical='center')
        
        # Make species name headers italic (columns B onwards)
        for col_idx in range(2, len(genome_names) + 2):
            header_cell = worksheet.cell(row=1, column=col_idx)
            header_cell.font = Font(bold=True, italic=True, color='FFFFFF', size=11)
        
        # Data formatting
        thin_border = Border(
            left=Side(style='thin'),
            right=Side(style='thin'),
            top=Side(style='thin'),
            bottom=Side(style='thin')
        )
        
        # Format data rows
        for row_idx, row in enumerate(worksheet.iter_rows(min_row=2, max_row=worksheet.max_row), start=2):
            for col_idx, cell in enumerate(row):
                cell.border = thin_border
                
                # First column (Amino_Acid): bold + italic, center aligned
                if col_idx == 0:
                    cell.alignment = Alignment(horizontal='center', vertical='center')
                    if cell.value:
                        cell.font = Font(bold=True, italic=True, size=10)
                
                # Percentage columns: format to 4 decimal places, center aligned
                else:
                    cell.alignment = Alignment(horizontal='center', vertical='center')
                    if cell.value is not None:
                        try:
                            cell.number_format = '0.0000'
                        except:
                            pass
        
        # Adjust column widths
        worksheet.column_dimensions['A'].width = 15  # Amino_Acid
        
        # Set width for genome columns
        for col_idx in range(2, len(genome_names) + 2):
            col_letter = worksheet.cell(row=1, column=col_idx).column_letter
            worksheet.column_dimensions[col_letter].width = 14
        
        # Freeze panes (freeze first row and first column)
        worksheet.freeze_panes = 'B2'
    
    print(f"\n  ✓ Merged analysis saved: {output_file}")
    print(f"  - Total genomes compared: {len(genome_names)}")
    print(f"  - Total amino acids: {len(merged_data)}")
    print(f"  - Format: Publication-ready with italic species names")


# ============================================================================
# MAIN PIPELINE
# ============================================================================

def main():
    """
    Main pipeline execution function.
    
    Workflow:
    1. Find all GenBank files in working directory
    2. Process each file to calculate amino acid composition
    3. Merge all composition files into comparative analysis
    """
    print(f"\n{'='*70}")
    print("AMINO ACID COMPOSITION ANALYSIS PIPELINE")
    print(f"{'='*70}")
    
    # Get working directory
    working_dir = os.getcwd()
    print(f"\nWorking directory: {working_dir}")
    
    # Find GenBank files
    genbank_files = find_genbank_files(working_dir)
    
    if not genbank_files:
        print("\n❌ ERROR: No GenBank files found!")
        print("Please place GenBank files (.gb, .gbf, .gbk) in the working directory.")
        return
    
    print(f"\nFound {len(genbank_files)} GenBank file(s):")
    for gf in genbank_files:
        print(f"  - {os.path.basename(gf)}")
    
    # Process each GenBank file
    print(f"\n{'='*70}")
    print("CALCULATING AMINO ACID COMPOSITION")
    print(f"{'='*70}")
    
    aa_files = []
    for gb_file in genbank_files:
        try:
            output_file, _ = process_single_genbank_file(gb_file)
            if output_file:
                aa_files.append(output_file)
        except Exception as e:
            print(f"  ❌ ERROR processing {gb_file}: {e}")
            continue
    
    # Merge composition files
    if aa_files:
        merge_amino_acid_files(aa_files)
    
    # Summary
    print(f"\n{'='*70}")
    print("ANALYSIS COMPLETE")
    print(f"{'='*70}")
    print(f"✓ Successfully processed: {len(aa_files)} genome(s)")
    print(f"✓ Individual composition files: {len(aa_files)}")
    print(f"✓ Merged analysis file: Merged_AminoAcid_Analysis.xlsx")
    print(f"\nAll output files saved in: {working_dir}")
    print(f"{'='*70}\n")


# ============================================================================
# SCRIPT ENTRY POINT
# ============================================================================

if __name__ == "__main__":
    try:
        main()
    except KeyboardInterrupt:
        print("\n\n⚠ Analysis interrupted by user.")
        sys.exit(0)
    except Exception as e:
        print(f"\n\n❌ FATAL ERROR: {e}")
        import traceback
        traceback.print_exc()
        sys.exit(1)
```
