## Supplementary Date File 1 for "CGAS (Chloroplast Genome Analysis Suite): An Automated Python Pipeline for Comprehensive Comparative Chloroplast Genomics": module6_snp_analysis.py

```
#!/usr/bin/env python3
"""
Nucleotide Substitution Analysis Pipeline
==========================================

This script performs comprehensive nucleotide substitution analysis on FASTA alignment files by:
1. Comparing pairwise aligned sequences from FASTA files
2. Identifying and counting all types of nucleotide substitutions
3. Calculating transition (Ts) and transversion (Tv) counts and ratios
4. Recording positions of each substitution type
5. Generating individual Excel reports for each alignment
6. Creating a merged comparative analysis across all alignments

Author: Bioinformatics Analysis Tool
Version: 1.0
Date: 2025

Dependencies:
    - openpyxl (for Excel file handling)

Usage:
    Place FASTA alignment files (.fasta, .fa, .fna) in the working directory and run:
    python substitution_analysis.py

Output:
    - Individual substitution files: [alignment_name]_Substitutions.xlsx
    - Merged analysis: Merged_Substitution_Analysis.xlsx

Notes:
    - FASTA files must contain exactly 2 sequences (reference and query)
    - Sequences must be pre-aligned and of equal length
    - Gaps (-) are ignored in the analysis
"""

import os
import sys
from pathlib import Path
from typing import Dict, List, Tuple

try:
    import openpyxl
    from openpyxl.styles import Font, Alignment, PatternFill, Border, Side
except ImportError as e:
    print(f"Error: Required package not installed: {e}")
    print("Please install required packages using:")
    print("pip install openpyxl")
    sys.exit(1)


# ============================================================================
# SUBSTITUTION TYPE DEFINITIONS
# ============================================================================

# All possible nucleotide substitutions
SUBSTITUTION_TYPES = [
    'A_to_G', 'G_to_A', 'T_to_G', 'G_to_T',
    'A_to_C', 'C_to_A', 'C_to_T', 'T_to_C',
    'G_to_C', 'C_to_G', 'A_to_T', 'T_to_A'
]

# Transition substitutions (purine-purine or pyrimidine-pyrimidine)
TRANSITIONS = ['A_to_G', 'G_to_A', 'C_to_T', 'T_to_C']

# Transversion substitutions (purine-pyrimidine or pyrimidine-purine)
TRANSVERSIONS = ['T_to_G', 'G_to_T', 'A_to_C', 'C_to_A', 
                 'G_to_C', 'C_to_G', 'A_to_T', 'T_to_A']


# ============================================================================
# SEQUENCE LOADING
# ============================================================================

def load_fasta_sequences(fasta_file: str) -> Tuple[List[str], List[str]]:
    """
    Load sequences from a FASTA file.
    
    Parameters:
    -----------
    fasta_file : str
        Path to the FASTA file
        
    Returns:
    --------
    Tuple[List[str], List[str]]
        Tuple of (sequence_names, sequences)
        
    Raises:
    -------
    ValueError
        If file doesn't contain exactly 2 sequences
        If sequences are not of equal length
    """
    sequence_names = []
    sequences = []
    current_sequence = ''
    current_name = ''
    
    try:
        with open(fasta_file, 'r') as file:
            for line in file:
                line = line.strip()
                
                if line.startswith('>'):
                    # Save previous sequence
                    if current_sequence:
                        sequences.append(current_sequence.upper())
                        sequence_names.append(current_name)
                    
                    # Start new sequence
                    current_name = line[1:].strip()
                    current_sequence = ''
                else:
                    current_sequence += line
            
            # Add last sequence
            if current_sequence:
                sequences.append(current_sequence.upper())
                sequence_names.append(current_name)
    
    except Exception as e:
        raise ValueError(f"Error reading FASTA file: {e}")
    
    # Validate sequences
    if len(sequences) != 2:
        raise ValueError(f"Expected 2 sequences, found {len(sequences)}. "
                        "This script requires pairwise alignments.")
    
    if len(sequences[0]) != len(sequences[1]):
        raise ValueError(f"Sequences have different lengths: {len(sequences[0])} vs {len(sequences[1])}. "
                        "Sequences must be pre-aligned.")
    
    return sequence_names, sequences


# ============================================================================
# SUBSTITUTION ANALYSIS
# ============================================================================

def analyze_substitutions(sequences: List[str]) -> Dict:
    """
    Analyze nucleotide substitutions between two aligned sequences.
    
    Parameters:
    -----------
    sequences : List[str]
        List containing exactly 2 aligned sequences
        
    Returns:
    --------
    Dict
        Dictionary containing:
        - substitutions: count of each substitution type
        - positions: list of positions for each substitution type
        - ts_count: total transition count
        - tv_count: total transversion count
        - ts_tv_ratio: transition/transversion ratio
        - total_compared: number of positions compared
        - identical: number of identical positions
    """
    # Initialize counters
    substitutions = {sub_type: 0 for sub_type in SUBSTITUTION_TYPES}
    positions = {sub_type: [] for sub_type in SUBSTITUTION_TYPES}
    
    ref_seq = sequences[0]
    query_seq = sequences[1]
    
    total_compared = 0
    identical = 0
    
    # Compare sequences position by position
    for i in range(len(ref_seq)):
        ref_base = ref_seq[i]
        query_base = query_seq[i]
        
        # Skip gaps
        if ref_base == '-' or query_base == '-':
            continue
        
        # Skip non-standard bases
        if ref_base not in 'ATGC' or query_base not in 'ATGC':
            continue
        
        total_compared += 1
        
        if ref_base == query_base:
            identical += 1
            continue
        
        # Record substitution
        substitution = f"{ref_base}_to_{query_base}"
        if substitution in substitutions:
            substitutions[substitution] += 1
            positions[substitution].append(i + 1)  # 1-based position
    
    # Calculate transition and transversion counts
    ts_count = sum(substitutions[sub] for sub in TRANSITIONS)
    tv_count = sum(substitutions[sub] for sub in TRANSVERSIONS)
    
    # Calculate Ts/Tv ratio
    ts_tv_ratio = ts_count / tv_count if tv_count > 0 else float('inf')
    
    return {
        'substitutions': substitutions,
        'positions': positions,
        'ts_count': ts_count,
        'tv_count': tv_count,
        'ts_tv_ratio': ts_tv_ratio,
        'total_compared': total_compared,
        'identical': identical
    }


# ============================================================================
# EXCEL OUTPUT
# ============================================================================

def create_substitution_excel(analysis_results: Dict, sequence_names: List[str], 
                             output_file: str, alignment_length: int):
    """
    Create an Excel file with substitution analysis results.
    
    Parameters:
    -----------
    analysis_results : Dict
        Results from analyze_substitutions()
    sequence_names : List[str]
        Names of the two sequences analyzed
    output_file : str
        Path for the output Excel file
    alignment_length : int
        Total length of the alignment
    """
    workbook = openpyxl.Workbook()
    
    # Remove default sheet
    if 'Sheet' in workbook.sheetnames:
        workbook.remove(workbook['Sheet'])
    
    # ===== Sheet 1: Substitution Details =====
    sheet_details = workbook.create_sheet('Substitution_Details', 0)
    
    # Headers
    headers = ['Substitution', 'Count', 'Percentage', 'Positions']
    sheet_details.append(headers)
    
    # Format headers
    header_fill = PatternFill(start_color='366092', end_color='366092', fill_type='solid')
    for cell in sheet_details[1]:
        cell.fill = header_fill
        cell.font = Font(bold=True, color='FFFFFF', size=11)
        cell.alignment = Alignment(horizontal='center', vertical='center')
    
    # Add substitution data
    total_substitutions = sum(analysis_results['substitutions'].values())
    
    for sub_type in SUBSTITUTION_TYPES:
        count = analysis_results['substitutions'][sub_type]
        percentage = (count / total_substitutions * 100) if total_substitutions > 0 else 0
        positions = analysis_results['positions'][sub_type]
        positions_str = ', '.join(map(str, positions)) if positions else 'None'
        
        row = [sub_type, count, round(percentage, 4), positions_str]
        sheet_details.append(row)
    
    # Add spacing
    sheet_details.append([])
    
    # Add summary statistics
    sheet_details.append(['Category', 'Value', '', ''])
    summary_row = sheet_details.max_row
    sheet_details[summary_row][0].font = Font(bold=True)
    sheet_details[summary_row][1].font = Font(bold=True)
    
    sheet_details.append(['Transitions (Ts)', analysis_results['ts_count'], '', ''])
    sheet_details.append(['Transversions (Tv)', analysis_results['tv_count'], '', ''])
    sheet_details.append(['Ts/Tv Ratio', round(analysis_results['ts_tv_ratio'], 4), '', ''])
    sheet_details.append(['Total Substitutions', total_substitutions, '', ''])
    
    # Format percentage column
    for row in range(2, len(SUBSTITUTION_TYPES) + 2):
        cell = sheet_details.cell(row=row, column=3)
        cell.number_format = '0.0000'
        cell.alignment = Alignment(horizontal='center')
    
    # Adjust column widths
    sheet_details.column_dimensions['A'].width = 15
    sheet_details.column_dimensions['B'].width = 10
    sheet_details.column_dimensions['C'].width = 12
    sheet_details.column_dimensions['D'].width = 50
    
    # ===== Sheet 2: Summary =====
    sheet_summary = workbook.create_sheet('Summary', 1)
    
    summary_data = [
        ['Metric', 'Value'],
        ['Reference Sequence', sequence_names[0]],
        ['Query Sequence', sequence_names[1]],
        ['Alignment Length', alignment_length],
        ['Positions Compared', analysis_results['total_compared']],
        ['Identical Positions', analysis_results['identical']],
        ['Total Substitutions', total_substitutions],
        ['Identity (%)', round(analysis_results['identical'] / analysis_results['total_compared'] * 100, 2) 
         if analysis_results['total_compared'] > 0 else 0],
        ['Substitution Rate (%)', round(total_substitutions / analysis_results['total_compared'] * 100, 2)
         if analysis_results['total_compared'] > 0 else 0],
        ['Transitions (Ts)', analysis_results['ts_count']],
        ['Transversions (Tv)', analysis_results['tv_count']],
        ['Ts/Tv Ratio', round(analysis_results['ts_tv_ratio'], 4)]
    ]
    
    for row_data in summary_data:
        sheet_summary.append(row_data)
    
    # Format summary sheet
    for cell in sheet_summary[1]:
        cell.fill = header_fill
        cell.font = Font(bold=True, color='FFFFFF', size=11)
        cell.alignment = Alignment(horizontal='center')
    
    for row in range(2, sheet_summary.max_row + 1):
        sheet_summary.cell(row=row, column=1).font = Font(bold=True)
    
    sheet_summary.column_dimensions['A'].width = 25
    sheet_summary.column_dimensions['B'].width = 40
    
    # Save workbook
    workbook.save(output_file)


# ============================================================================
# FILE PROCESSING
# ============================================================================

def process_single_fasta_file(fasta_file: str) -> Tuple[str, Dict]:
    """
    Process a single FASTA file and analyze substitutions.
    
    Parameters:
    -----------
    fasta_file : str
        Path to the FASTA file
        
    Returns:
    --------
    Tuple[str, Dict]
        Tuple of (output filename, analysis results)
    """
    print(f"\nProcessing: {fasta_file}")
    
    try:
        # Load sequences
        sequence_names, sequences = load_fasta_sequences(fasta_file)
        print(f"  - Reference: {sequence_names[0]}")
        print(f"  - Query: {sequence_names[1]}")
        print(f"  - Alignment length: {len(sequences[0]):,} bp")
        
        # Analyze substitutions
        results = analyze_substitutions(sequences)
        
        total_subs = sum(results['substitutions'].values())
        print(f"  - Total substitutions: {total_subs:,}")
        print(f"  - Transitions: {results['ts_count']:,}")
        print(f"  - Transversions: {results['tv_count']:,}")
        print(f"  - Ts/Tv ratio: {results['ts_tv_ratio']:.4f}")
        
        # Generate output filename
        base_name = Path(fasta_file).stem
        output_file = f"{base_name}_Substitutions.xlsx"
        
        # Create Excel file
        create_substitution_excel(results, sequence_names, output_file, len(sequences[0]))
        
        print(f"  ✓ Saved: {output_file}")
        
        return output_file, results
        
    except Exception as e:
        print(f"  ❌ ERROR: {e}")
        return None, None


def find_fasta_files(directory: str = None) -> List[str]:
    """
    Find all FASTA files in the specified directory.
    
    Parameters:
    -----------
    directory : str, optional
        Directory to search (default: current working directory)
        
    Returns:
    --------
    List[str]
        List of FASTA file paths
    """
    if directory is None:
        directory = os.getcwd()
    
    # Supported FASTA file extensions
    fasta_extensions = ('.fasta', '.fa', '.fna', '.faa')
    
    fasta_files = [
        os.path.join(directory, f) 
        for f in os.listdir(directory) 
        if f.lower().endswith(fasta_extensions)
    ]
    
    return sorted(fasta_files)


# ============================================================================
# MERGE RESULTS
# ============================================================================

def merge_substitution_files(sub_files: List[str], output_file: str = 'Merged_Substitution_Analysis.xlsx'):
    """
    Merge individual substitution analysis files into a single comparative analysis file.
    
    Parameters:
    -----------
    sub_files : List[str]
        List of substitution Excel file paths to merge
    output_file : str
        Name of the output merged file
    """
    print(f"\n{'='*70}")
    print("MERGING SUBSTITUTION ANALYSIS FILES")
    print(f"{'='*70}")
    
    if not sub_files:
        print("No substitution files to merge.")
        return
    
    # Collect data from all files
    all_data = {}
    file_names = []
    
    for filename in sub_files:
        print(f"  - Adding: {filename}")
        
        try:
            workbook = openpyxl.load_workbook(filename)
            sheet = workbook['Substitution_Details']
            
            # Get file identifier
            file_id = Path(filename).stem.replace('_Substitutions', '')
            file_names.append(file_id)
            
            # Extract substitution counts
            data = {}
            for row in range(2, 14):  # Rows 2-13 contain substitution data
                sub_type = sheet.cell(row=row, column=1).value
                count = sheet.cell(row=row, column=2).value
                data[sub_type] = count
            
            # Extract Ts, Tv, and ratio
            for row in range(15, sheet.max_row + 1):
                category = sheet.cell(row=row, column=1).value
                value = sheet.cell(row=row, column=2).value
                if category in ['Transitions (Ts)', 'Transversions (Tv)', 'Ts/Tv Ratio']:
                    data[category] = value
            
            all_data[file_id] = data
            workbook.close()
            
        except Exception as e:
            print(f"  WARNING: Error reading {filename}: {e}")
            continue
    
    if not all_data:
        print("  ERROR: No data to merge.")
        return
    
    # Create merged workbook
    merged_wb = openpyxl.Workbook()
    merged_sheet = merged_wb.active
    merged_sheet.title = 'Merged_Substitutions'
    
    # Build header row
    header = ['Substitution_Type'] + file_names
    merged_sheet.append(header)
    
    # Add substitution data
    for sub_type in SUBSTITUTION_TYPES:
        row_data = [sub_type]
        for file_id in file_names:
            row_data.append(all_data[file_id].get(sub_type, 0))
        merged_sheet.append(row_data)
    
    # Add spacing
    merged_sheet.append([])
    
    # Add summary statistics
    for category in ['Transitions (Ts)', 'Transversions (Tv)', 'Ts/Tv Ratio']:
        row_data = [category]
        for file_id in file_names:
            row_data.append(all_data[file_id].get(category, 0))
        merged_sheet.append(row_data)
    
    # Format the merged sheet
    header_fill = PatternFill(start_color='366092', end_color='366092', fill_type='solid')
    
    for cell in merged_sheet[1]:
        cell.fill = header_fill
        cell.font = Font(bold=True, color='FFFFFF', size=11)
        cell.alignment = Alignment(horizontal='center', vertical='center')
    
    # Format data cells
    for row in merged_sheet.iter_rows(min_row=2, max_row=merged_sheet.max_row):
        row[0].font = Font(bold=True)
        row[0].alignment = Alignment(horizontal='left')
        
        for cell in row[1:]:
            cell.alignment = Alignment(horizontal='center')
    
    # Adjust column widths
    merged_sheet.column_dimensions['A'].width = 20
    for col_idx in range(2, len(file_names) + 2):
        col_letter = merged_sheet.cell(row=1, column=col_idx).column_letter
        merged_sheet.column_dimensions[col_letter].width = 15
    
    # Freeze panes
    merged_sheet.freeze_panes = 'B2'
    
    # Save merged file
    merged_wb.save(output_file)
    
    print(f"\n  ✓ Merged analysis saved: {output_file}")
    print(f"  - Total alignments compared: {len(file_names)}")


# ============================================================================
# MAIN PIPELINE
# ============================================================================

def main():
    """
    Main pipeline execution function.
    
    Workflow:
    1. Find all FASTA files in working directory
    2. Process each file to analyze substitutions
    3. Merge all results into comparative analysis
    """
    print(f"\n{'='*70}")
    print("NUCLEOTIDE SUBSTITUTION ANALYSIS PIPELINE")
    print(f"{'='*70}")
    
    # Get working directory
    working_dir = os.getcwd()
    print(f"\nWorking directory: {working_dir}")
    
    # Find FASTA files
    fasta_files = find_fasta_files(working_dir)
    
    if not fasta_files:
        print("\n❌ ERROR: No FASTA files found!")
        print("Please place FASTA alignment files (.fasta, .fa, .fna) in the working directory.")
        return
    
    print(f"\nFound {len(fasta_files)} FASTA file(s):")
    for ff in fasta_files:
        print(f"  - {os.path.basename(ff)}")
    
    # Process each FASTA file
    print(f"\n{'='*70}")
    print("ANALYZING NUCLEOTIDE SUBSTITUTIONS")
    print(f"{'='*70}")
    
    sub_files = []
    for fasta_file in fasta_files:
        try:
            output_file, _ = process_single_fasta_file(fasta_file)
            if output_file:
                sub_files.append(output_file)
        except Exception as e:
            print(f"  ❌ ERROR processing {fasta_file}: {e}")
            continue
    
    # Merge results
    if sub_files:
        merge_substitution_files(sub_files)
    
    # Summary
    print(f"\n{'='*70}")
    print("ANALYSIS COMPLETE")
    print(f"{'='*70}")
    print(f"✓ Successfully processed: {len(sub_files)} alignment(s)")
    print(f"✓ Individual substitution files: {len(sub_files)}")
    if sub_files:
        print(f"✓ Merged analysis file: Merged_Substitution_Analysis.xlsx")
    print(f"\nAll output files saved in: {working_dir}")
    print(f"{'='*70}\n")


# ============================================================================
# SCRIPT ENTRY POINT
# ============================================================================

if __name__ == "__main__":
    try:
        main()
    except KeyboardInterrupt:
        print("\n\n⚠ Analysis interrupted by user.")
        sys.exit(0)
    except Exception as e:
        print(f"\n\n❌ FATAL ERROR: {e}")
        import traceback
        traceback.print_exc()
        sys.exit(1)
```
