## Supplementary Date File 1 for "CGAS (Chloroplast Genome Analysis Suite): An Automated Python Pipeline for Comprehensive Comparative Chloroplast Genomics": module7_intron_extraction.py

```
#!/usr/bin/env python3
"""
Gene and tRNA Intron Extraction from GenBank Files

This script extracts intron positions and lengths from both gene (CDS/gene features)
and tRNA features in GenBank format files. Results are saved in a single Excel file
with separate sheets for gene introns and tRNA introns.

Author: [Your Name]
Date: December 2025
Version: 2.0

Requirements:
    - Python 3.7+
    - biopython
    - pandas
    - openpyxl

Installation:
    pip install biopython pandas openpyxl

Usage:
    python extract_introns.py

Output:
    intron_data.xlsx - Excel file with two sheets:
        - Sheet 1: Gene Introns (CDS/gene features)
        - Sheet 2: tRNA Introns (tRNA features)
"""

import os
import sys
import logging
from datetime import datetime
from typing import List, Tuple
import pandas as pd
from Bio import SeqIO
from Bio.SeqFeature import CompoundLocation

# Configure logging
logging.basicConfig(
    level=logging.INFO,
    format='%(asctime)s - %(levelname)s - %(message)s',
    handlers=[
        logging.FileHandler('intron_extraction.log', encoding='utf-8'),
        logging.StreamHandler(sys.stdout)
    ]
)
logger = logging.getLogger(__name__)

# Handle Windows console encoding
if sys.platform == 'win32':
    try:
        if hasattr(sys.stdout, 'reconfigure'):
            sys.stdout.reconfigure(encoding='utf-8')
    except (AttributeError, OSError):
        pass

# Global constants
MAX_INTRON_LENGTH = 15000
OUTPUT_FILENAME = "intron_data.xlsx"
VALID_EXTENSIONS = ('.gb', '.gbff', '.genbank')


def extract_gene_introns_from_genbank(gb_file: str) -> List[List]:
    """
    Extract gene/CDS intron information from a GenBank file.
    
    Args:
        gb_file (str): Path to GenBank format file
        
    Returns:
        List[List]: List of gene records with intron data
    """
    gene_introns = []
    
    for record in SeqIO.parse(gb_file, "genbank"):
        accession = record.id
        species = record.annotations.get('organism', 'Unknown species')
        
        for feature in record.features:
            if feature.type in ['CDS', 'gene']:
                gene = feature.qualifiers.get('gene', ['Unknown'])[0]
                
                if isinstance(feature.location, CompoundLocation):
                    exons = sorted(feature.location.parts, key=lambda x: x.start)
                    intron_info = []
                    
                    for i in range(len(exons) - 1):
                        intron_start = int(exons[i].end) + 1
                        intron_end = int(exons[i + 1].start)
                        intron_length = intron_end - intron_start + 1
                        
                        if 0 < intron_length <= MAX_INTRON_LENGTH:
                            intron_info.extend([
                                f"Intron {i + 1}",
                                intron_start,
                                intron_end,
                                intron_length
                            ])
                    
                    if intron_info:
                        gene_data = [accession, species, gene] + intron_info
                        gene_introns.append(gene_data)
    
    return gene_introns


def extract_tRNA_introns_from_genbank(gb_file: str) -> List[List]:
    """
    Extract tRNA intron information from a GenBank file.
    
    Args:
        gb_file (str): Path to GenBank format file
        
    Returns:
        List[List]: List of tRNA records with intron data
    """
    trna_introns = []
    
    for record in SeqIO.parse(gb_file, "genbank"):
        accession = record.id
        species = record.annotations.get('organism', 'Unknown species')
        
        for feature in record.features:
            if feature.type == 'tRNA':
                gene = feature.qualifiers.get('gene', feature.qualifiers.get('product', ['Unknown']))[0]
                
                if isinstance(feature.location, CompoundLocation):
                    exons = sorted(feature.location.parts, key=lambda x: x.start)
                    intron_info = []
                    
                    for i in range(len(exons) - 1):
                        intron_start = exons[i].end + 1
                        intron_end = exons[i + 1].start - 1
                        intron_length = int(intron_end - intron_start + 1)
                        
                        if intron_length > 0 and intron_length <= MAX_INTRON_LENGTH:
                            intron_info.extend([
                                f"Intron {i + 1}",
                                int(intron_start),
                                int(intron_end),
                                intron_length
                            ])
                    
                    if intron_info:
                        gene_data = [accession, species, gene] + intron_info
                        trna_introns.append(gene_data)
    
    return trna_introns


def create_combined_excel_output(gene_data: List[List], trna_data: List[List], 
                                 output_excel: str) -> None:
    """
    Create Excel output with separate sheets for gene and tRNA introns.
    
    Args:
        gene_data (List[List]): Gene intron data rows
        trna_data (List[List]): tRNA intron data rows
        output_excel (str): Output Excel filename
    """
    with pd.ExcelWriter(output_excel, engine='openpyxl') as writer:
        from openpyxl.styles import Font, Alignment
        
        # Define fonts
        header_font = Font(bold=True)
        header_alignment = Alignment(horizontal='center')
        italic_font = Font(italic=True)
        regular_font = Font(italic=False)
        center_alignment = Alignment(horizontal='center')
        left_alignment = Alignment(horizontal='left')
        
        # Create Gene Introns sheet
        if gene_data:
            max_introns = max((len(row) - 3) // 4 for row in gene_data)
            columns = ["Accession", "Species", "Gene"]
            for i in range(1, max_introns + 1):
                columns += [f"Intron {i} #", f"Intron {i} Start", f"Intron {i} End", f"Intron {i} Length"]
            
            for row in gene_data:
                while len(row) < len(columns):
                    row.extend(["", "", "", ""])
            
            df_gene = pd.DataFrame(gene_data, columns=columns)
            df_gene.to_excel(writer, index=False, sheet_name='Gene Introns')
            
            worksheet = writer.sheets['Gene Introns']
            
            # Format header row
            for cell in worksheet[1]:
                cell.font = header_font
                cell.alignment = header_alignment
            
            # Format data rows
            for row in worksheet.iter_rows(min_row=2, max_row=worksheet.max_row):
                # Column A: Accession (regular, left-aligned)
                row[0].font = regular_font
                row[0].alignment = left_alignment
                
                # Column B: Species (italic, left-aligned)
                row[1].font = italic_font
                row[1].alignment = left_alignment
                
                # Column C: Gene (italic, left-aligned)
                row[2].font = italic_font
                row[2].alignment = left_alignment
                
                # Remaining columns: numbers (regular, centered)
                for cell in row[3:]:
                    cell.font = regular_font
                    cell.alignment = center_alignment
            
            # Auto-adjust column widths
            for column in worksheet.columns:
                max_length = 0
                column_letter = column[0].column_letter
                for cell in column:
                    if cell.value:
                        max_length = max(max_length, len(str(cell.value)))
                worksheet.column_dimensions[column_letter].width = min(max_length + 3, 50)
        
        # Create tRNA Introns sheet
        if trna_data:
            max_introns = max((len(row) - 3) // 4 for row in trna_data)
            columns = ["Accession", "Species", "tRNA Gene"]
            for i in range(1, max_introns + 1):
                columns += [f"Intron {i} #", f"Intron {i} Start", f"Intron {i} End", f"Intron {i} Length"]
            
            for row in trna_data:
                while len(row) < len(columns):
                    row.extend(["", "", "", ""])
            
            df_trna = pd.DataFrame(trna_data, columns=columns)
            df_trna.to_excel(writer, index=False, sheet_name='tRNA Introns')
            
            worksheet = writer.sheets['tRNA Introns']
            
            # Format header row
            for cell in worksheet[1]:
                cell.font = header_font
                cell.alignment = header_alignment
            
            # Format data rows
            for row in worksheet.iter_rows(min_row=2, max_row=worksheet.max_row):
                # Column A: Accession (regular, left-aligned)
                row[0].font = regular_font
                row[0].alignment = left_alignment
                
                # Column B: Species (italic, left-aligned)
                row[1].font = italic_font
                row[1].alignment = left_alignment
                
                # Column C: tRNA Gene (italic, left-aligned)
                row[2].font = italic_font
                row[2].alignment = left_alignment
                
                # Remaining columns: numbers (regular, centered)
                for cell in row[3:]:
                    cell.font = regular_font
                    cell.alignment = center_alignment
            
            # Auto-adjust column widths
            for column in worksheet.columns:
                max_length = 0
                column_letter = column[0].column_letter
                for cell in column:
                    if cell.value:
                        max_length = max(max_length, len(str(cell.value)))
                worksheet.column_dimensions[column_letter].width = min(max_length + 3, 50)


def process_all_genbank_files(output: str = OUTPUT_FILENAME) -> None:
    """
    Process all GenBank files in the current working directory.
    
    Args:
        output (str): Output Excel filename
    """
    logger.info("="*70)
    logger.info("INTRON EXTRACTION ANALYSIS")
    logger.info("="*70)
    
    all_gene_introns = []
    all_trna_introns = []
    current_dir = os.getcwd()
    
    gb_files = [f for f in os.listdir(current_dir) if f.endswith(VALID_EXTENSIONS)]
    
    if not gb_files:
        logger.error(f"No GenBank files found in {current_dir}")
        raise FileNotFoundError("No GenBank files (.gb, .gbff, .genbank) found")
    
    logger.info(f"Found {len(gb_files)} GenBank file(s)")
    logger.info("-"*70)
    
    # Process each file
    for idx, gb_file in enumerate(gb_files, 1):
        logger.info(f"[{idx}/{len(gb_files)}] Processing: {gb_file}")
        try:
            gene_rows = extract_gene_introns_from_genbank(gb_file)
            trna_rows = extract_tRNA_introns_from_genbank(gb_file)
            all_gene_introns.extend(gene_rows)
            all_trna_introns.extend(trna_rows)
            logger.info(f"  Genes: {len(gene_rows)}, tRNAs: {len(trna_rows)}")
        except Exception as e:
            logger.warning(f"  Skipping {gb_file}: {str(e)}")
            continue
    
    logger.info("-"*70)
    
    if not all_gene_introns and not all_trna_introns:
        raise ValueError("No introns found in any file")
    
    # Create Excel file
    logger.info(f"Creating: {output}")
    create_combined_excel_output(all_gene_introns, all_trna_introns, output)
    
    logger.info("="*70)
    logger.info(f"SUCCESS: {output} created")
    logger.info(f"  Gene introns: {len(all_gene_introns)}")
    logger.info(f"  tRNA introns: {len(all_trna_introns)}")
    logger.info("="*70)


def main():
    """Main execution function."""
    try:
        process_all_genbank_files()
    except Exception as e:
        logger.error(f"Error: {str(e)}")
        sys.exit(1)


if __name__ == "__main__":
    main()
```
