## Supplementary Date File 1 for "CGAS (Chloroplast Genome Analysis Suite): An Automated Python Pipeline for Comprehensive Comparative Chloroplast Genomics": module8_ssr_analysis.py

```
#!/usr/bin/env python3
"""
Comprehensive Chloroplast SSR Analyzer
Processes GenBank files to identify SSRs and classify them by:
1. Genomic region (LSC/SSC/IR)
2. Motif type (Mono/Di/Tri/Tetra/Penta/Hexa)
3. Genomic location (gene names, intergenic spacers)
"""

import os
import sys
import pandas as pd
from Bio import SeqIO
from collections import defaultdict
import re

class ChloroplastSSRAnalyzer:
    """Main analyzer class for SSR detection and classification"""
    
    def __init__(self, gb_folder, output_folder="ssr_analysis_results"):
        self.gb_folder = gb_folder
        self.output_folder = output_folder
        self.thresholds = {1: 10, 2: 5, 3: 4, 4: 3, 5: 3, 6: 3}
        os.makedirs(output_folder, exist_ok=True)
        
    def find_ssrs(self, sequence):
        """Detect SSRs in sequence based on motif thresholds"""
        ssrs = []
        seq_upper = str(sequence).upper()
        seq_len = len(seq_upper)
        
        # Track positions already in SSRs to avoid overlaps
        covered = set()
        
        for motif_len in range(1, 7):  # 1-6 bp motifs
            min_repeats = self.thresholds[motif_len]
            
            # Slide through sequence
            i = 0
            while i < seq_len - motif_len * min_repeats + 1:
                # Skip if position already covered
                if i in covered:
                    i += 1
                    continue
                
                # Get potential motif
                motif = seq_upper[i:i + motif_len]
                
                # Skip if contains non-ATCG bases
                if not all(base in 'ATCG' for base in motif):
                    i += 1
                    continue
                
                # Count consecutive repeats of this motif
                repeat_count = 0
                pos = i
                while pos + motif_len <= seq_len and seq_upper[pos:pos + motif_len] == motif:
                    repeat_count += 1
                    pos += motif_len
                
                # If meets threshold, record SSR
                if repeat_count >= min_repeats:
                    ssr_start = i + 1  # 1-based
                    ssr_end = pos      # 1-based, inclusive
                    ssr_length = pos - i
                    
                    # Mark these positions as covered
                    for p in range(i, pos):
                        covered.add(p)
                    
                    ssrs.append((motif, ssr_length, ssr_start, ssr_end, repeat_count))
                    
                    # Jump past this SSR
                    i = pos
                else:
                    i += 1
        
        return sorted(ssrs, key=lambda x: x[2])
    
    def extract_regions(self, record):
        """Extract LSC, SSC, IR boundaries from GenBank record"""
        seq_len = len(record.seq)
        ir_features = []
        
        for feat in record.features:
            if feat.type != "repeat_region":
                continue
            
            note_text = " ".join(feat.qualifiers.get("note", []))
            rpt_type = " ".join(feat.qualifiers.get("rpt_type", []))
            combined = (note_text + " " + rpt_type).lower()
            
            if any(term in combined for term in ["inverted", "ir", "inverted repeat"]):
                try:
                    ir_start = int(feat.location.start) + 1
                    ir_end = int(feat.location.end)
                    ir_features.append((ir_start, ir_end))
                except:
                    continue
        
        if len(ir_features) >= 2:
            ir_features = sorted(ir_features, key=lambda x: x[0])
            irb_start, irb_end = ir_features[0]
            ira_start, ira_end = ir_features[1]
            
            lsc_start = 1
            lsc_end = irb_start - 1
            ssc_start = irb_end + 1
            ssc_end = ira_start - 1
            
            return {
                'LSC': (lsc_start, lsc_end),
                'SSC': (ssc_start, ssc_end),
                'IRb': (irb_start, irb_end),
                'IRa': (ira_start, ira_end)
            }
        elif len(ir_features) == 1:
            ir_start, ir_end = ir_features[0]
            lsc_start = 1
            lsc_end = ir_start - 1
            ssc_start = ir_end + 1
            ssc_end = seq_len
            
            return {
                'LSC': (lsc_start, lsc_end),
                'SSC': (ssc_start, ssc_end),
                'IR': (ir_start, ir_end)
            }
        else:
            return {'LSC': (1, seq_len)}
    
    def classify_region(self, pos, regions):
        """Classify SSR position into genomic region"""
        for region_name, (start, end) in regions.items():
            if start <= pos <= end:
                if region_name in ['IRb', 'IRa']:
                    return 'IR'
                return region_name
        return 'Unknown'
    
    def classify_motif(self, motif):
        """Classify SSR by motif length"""
        motif_types = {
            1: "Mono", 2: "Di", 3: "Tri",
            4: "Tetra", 5: "Penta", 6: "Hexa"
        }
        return motif_types.get(len(motif), "Other")
    
    def get_genomic_location(self, ssr_start, ssr_end, record):
        """Determine precise genomic location of SSR"""
        
        # Maximum intron length - anything longer is likely trans-spliced
        MAX_INTRON_LENGTH = 10000
        
        # Collect all features with their actual parts
        all_features = []
        
        for feat in record.features:
            if feat.type in ["gene", "tRNA", "rRNA", "CDS"]:
                gene_name = feat.qualifiers.get("gene", [""])[0]
                if not gene_name:
                    continue
                
                # Get overall span
                f_start = int(feat.location.start) + 1
                f_end = int(feat.location.end)
                
                # Get individual parts if split
                parts = []
                if hasattr(feat.location, 'parts'):
                    for part in feat.location.parts:
                        parts.append((int(part.start) + 1, int(part.end)))
                else:
                    parts = [(f_start, f_end)]
                
                # Calculate intron length if multi-part
                is_trans_spliced = False
                if len(parts) > 1:
                    # Check distance between parts
                    sorted_parts = sorted(parts, key=lambda x: x[0])
                    for i in range(len(sorted_parts) - 1):
                        intron_length = sorted_parts[i+1][0] - sorted_parts[i][1] - 1
                        if intron_length > MAX_INTRON_LENGTH:
                            is_trans_spliced = True
                            break
                
                # Also check for explicit trans_splicing qualifier
                if 'trans_splicing' in feat.qualifiers:
                    is_trans_spliced = True
                
                all_features.append({
                    'type': feat.type,
                    'name': gene_name,
                    'start': f_start,
                    'end': f_end,
                    'parts': parts,
                    'feature': feat,
                    'is_trans_spliced': is_trans_spliced
                })
        
        # Helper: check if SSR is in any part of a feature
        def is_in_parts(ssr_start, ssr_end, parts):
            for part_start, part_end in parts:
                if ssr_start >= part_start and ssr_end <= part_end:
                    return True
            return False
        
        # Priority 1: Check tRNAs
        trnas = [f for f in all_features if f['type'] == 'tRNA' and 
                 not f.get('is_trans_spliced', False) and
                 ssr_start >= f['start'] and ssr_end <= f['end']]
        
        if trnas:
            trna = trnas[0]
            if len(trna['parts']) > 1:
                # tRNA has intron
                if not is_in_parts(ssr_start, ssr_end, trna['parts']):
                    # SSR is in tRNA intron
                    # Check for embedded gene (like matK in trnK)
                    genes = [f for f in all_features if f['type'] == 'gene' and 
                            not f.get('is_trans_spliced', False) and
                            f['name'] != trna['name'] and
                            ssr_start >= f['start'] and ssr_end <= f['end']]
                    
                    if genes:
                        # Format: trnK-UUU_2-matK
                        return f"{trna['name']}_2-{genes[0]['name']}", "Intron"
                    else:
                        # Format: trnI-GAU_1-trnI-GAU_2 (showing intron between exons)
                        return f"{trna['name']}_1-{trna['name']}_2", "Intron"
            
            return trna['name'], "tRNA"
        
        # Priority 2: Check rRNAs
        rrnas = [f for f in all_features if f['type'] == 'rRNA' and 
                 ssr_start >= f['start'] and ssr_end <= f['end']]
        if rrnas:
            return rrnas[0]['name'], "rRNA"
        
        # Priority 3: Check genes (non-trans-spliced only)
        genes = [f for f in all_features if f['type'] == 'gene' and 
                 not f.get('is_trans_spliced', False) and
                 ssr_start >= f['start'] and ssr_end <= f['end']]
        
        if genes:
            gene = genes[0]
            gene_name = gene['name']
            
            # Find CDS for this gene
            cdss = [f for f in all_features if f['type'] == 'CDS' and 
                   f['name'] == gene_name and not f.get('is_trans_spliced', False)]
            
            if cdss:
                cds = cdss[0]
                
                # Check if SSR is in CDS exon
                if is_in_parts(ssr_start, ssr_end, cds['parts']):
                    # SSR is in exon - check if multi-part CDS
                    if len(cds['parts']) > 1:
                        # Find which part
                        for i, (pstart, pend) in enumerate(cds['parts'], 1):
                            if ssr_start >= pstart and ssr_end <= pend:
                                return f"{gene_name}_{i}", "Gene"
                    return gene_name, "Gene"
                
                # SSR is in gene but not in CDS = intron
                if len(cds['parts']) > 1:
                    # Multi-part CDS - check which intron
                    sorted_parts = sorted(cds['parts'], key=lambda x: x[0])
                    for i in range(len(sorted_parts) - 1):
                        intron_start = sorted_parts[i][1] + 1
                        intron_end = sorted_parts[i+1][0] - 1
                        if ssr_start >= intron_start and ssr_end <= intron_end:
                            # Format: gene_2-gene_1 (higher number first for reverse strand)
                            # Check strand orientation
                            if sorted_parts[0][0] > sorted_parts[1][0]:  # reverse strand
                                return f"{gene_name}_{i+2}-{gene_name}_{i+1}", "Intron"
                            else:
                                return f"{gene_name}_{i+1}-{gene_name}_{i+2}", "Intron"
                    
                    return f"{gene_name}_1-{gene_name}_2", "Intron"
                else:
                    return gene_name, "Intron"
            else:
                # Gene has no CDS
                return gene_name, "Gene"
        
        # Priority 3b: Check if in trans-spliced CDS parts only (not gene span)
        # For trans-spliced genes, only check if SSR is directly in one of the CDS parts
        for cds_feat in [f for f in all_features if f['type'] == 'CDS' and f.get('is_trans_spliced', False)]:
            if is_in_parts(ssr_start, ssr_end, cds_feat['parts']):
                # Find which part
                for i, (pstart, pend) in enumerate(cds_feat['parts'], 1):
                    if ssr_start >= pstart and ssr_end <= pend:
                        if len(cds_feat['parts']) > 1:
                            return f"{cds_feat['name']}_{i}", "Gene"
                        else:
                            return cds_feat['name'], "Gene"
        
        # Priority 4: Intergenic
        flanking = self.find_flanking_genes_with_parts(ssr_start, ssr_end, all_features)
        if flanking:
            return flanking, "IGS"
        
        return "intergenic", "IGS"
    
    def find_flanking_genes_with_parts(self, ssr_start, ssr_end, all_features):
        """Find flanking genes for IGS, with part numbers if applicable"""
        genes_before = []
        genes_after = []
        
        for feat in all_features:
            if feat['type'] not in ['gene', 'tRNA', 'rRNA']:
                continue
            
            # Skip trans-spliced genes
            if feat.get('is_trans_spliced', False):
                continue
            
            f_start = feat['start']
            f_end = feat['end']
            gene_name = feat['name']
            
            if f_end < ssr_start:
                # Find CDS to check if multi-part
                cdss = [f for f in all_features if f['type'] == 'CDS' and f['name'] == gene_name]
                if cdss and len(cdss[0]['parts']) > 1:
                    # Multi-part gene - find which part is closest
                    last_part_num = len(cdss[0]['parts'])
                    genes_before.append((f_end, f"{gene_name}_{last_part_num}"))
                else:
                    genes_before.append((f_end, gene_name))
            
            elif f_start > ssr_end:
                # Find CDS to check if multi-part
                cdss = [f for f in all_features if f['type'] == 'CDS' and f['name'] == gene_name]
                if cdss and len(cdss[0]['parts']) > 1:
                    # Multi-part gene - use first part
                    genes_after.append((f_start, f"{gene_name}_1"))
                elif feat['type'] == 'tRNA' and len(feat['parts']) > 1:
                    # Multi-part tRNA
                    genes_after.append((f_start, f"{gene_name}_2"))
                else:
                    genes_after.append((f_start, gene_name))
        
        if genes_before and genes_after:
            upstream = max(genes_before, key=lambda x: x[0])[1]
            downstream = min(genes_after, key=lambda x: x[0])[1]
            return f"{upstream}-{downstream}"
        
        return None
    
    def process_genbank_file(self, gb_file):
        """Process a single GenBank file"""
        try:
            record = SeqIO.read(gb_file, "genbank")
        except Exception as e:
            print(f"ERROR reading {gb_file}: {e}")
            return None
        
        species = os.path.splitext(os.path.basename(gb_file))[0]
        print(f"Processing {species}...")
        
        regions = self.extract_regions(record)
        ssrs = self.find_ssrs(record.seq)
        
        ssr_data = []
        for motif, length, start, end, repeat_count in ssrs:
            region = self.classify_region(start, regions)
            motif_type = self.classify_motif(motif)
            location, loc_type = self.get_genomic_location(start, end, record)
            
            ssr_data.append({
                'Species': species,
                'Type': motif,
                'Motif_Type': motif_type,
                'Repeat_Count': repeat_count,
                'Length': length,
                'Start': start,
                'End': end,
                'Region': region,
                'Location': location,
                'Location_Type': loc_type
            })
        
        return pd.DataFrame(ssr_data)
    
    def format_location_for_display(self, location, loc_type):
        """
        Format location string for publication display
        Returns: formatted text with gene names that should be italic
        """
        if loc_type == "Intron":
            # Handle different intron formats
            if "_2-" in location and location.count("_") == 2:
                # Format: trnK-UUU_2-matK -> trnK-UUU intron (for tRNA with embedded gene)
                parts = location.split("_2-")
                if len(parts) == 2:
                    gene1 = parts[0]
                    gene2 = parts[1]
                    # This is tRNA intron with embedded gene
                    return f"{gene1} intron"
            elif "_1-" in location and "_2" in location:
                # Format: trnK-UUU_1-trnK-UUU_2 or rps16_1-rps16_2 -> gene intron
                gene_name = location.split("_")[0]
                return f"{gene_name} intron"
            elif "-" in location:
                # Format: gene_N-gene_M -> gene intron
                gene_name = location.split("_")[0] if "_" in location else location.split("-")[0]
                return f"{gene_name} intron"
            # Fallback
            return f"{location} intron"
        
        return location
    
    def create_formatted_excel(self, all_data, output_file):
        """Create publication-quality Excel file with formatting"""
        from openpyxl import Workbook
        from openpyxl.styles import Font, Alignment, PatternFill, Border, Side
        from openpyxl.utils import get_column_letter
        
        wb = Workbook()
        ws = wb.active
        ws.title = "All SSRs"
        
        # Headers
        headers = ['Type', 'Motif_Type', 'Repeat_Count', 'Length', 'Start', 'End', 'Region', 'Location', 'Location_Type']
        
        # Write headers
        for col, header in enumerate(headers, 1):
            cell = ws.cell(1, col, header)
            cell.font = Font(bold=True)
            cell.alignment = Alignment(horizontal='center', vertical='center')
        
        row = 2
        
        # Sort by species, then by Start position
        all_data = all_data.sort_values(['Species', 'Start'])
        
        for species in sorted(all_data['Species'].unique()):
            species_data = all_data[all_data['Species'] == species].sort_values('Start')
            
            # Add species row (merged across all columns)
            ws.merge_cells(f'A{row}:I{row}')
            species_cell = ws.cell(row, 1, species.replace("_", " "))
            species_cell.font = Font(italic=True, bold=True, size=11)
            species_cell.alignment = Alignment(horizontal='center', vertical='center')
            species_cell.fill = PatternFill(start_color='E8E8E8', end_color='E8E8E8', fill_type='solid')
            row += 1
            
            # Add data rows for this species
            for idx, data_row in species_data.iterrows():
                # Format location for display
                location_display = self.format_location_for_display(
                    data_row['Location'], data_row['Location_Type']
                )
                
                # Write data
                ws.cell(row, 1, data_row['Type'])
                ws.cell(row, 2, data_row['Motif_Type'])
                ws.cell(row, 3, data_row['Repeat_Count'])
                ws.cell(row, 4, data_row['Length'])
                ws.cell(row, 5, data_row['Start'])
                ws.cell(row, 6, data_row['End'])
                ws.cell(row, 7, data_row['Region'])
                
                # Location cell - italicize if it's a gene name
                location_cell = ws.cell(row, 8, location_display)
                if data_row['Location_Type'] in ['Gene', 'Intron', 'tRNA', 'rRNA', 'IGS']:
                    location_cell.font = Font(italic=True)
                
                ws.cell(row, 9, data_row['Location_Type'])
                
                row += 1
        
        # Adjust column widths
        ws.column_dimensions['A'].width = 10
        ws.column_dimensions['B'].width = 12
        ws.column_dimensions['C'].width = 14
        ws.column_dimensions['D'].width = 10
        ws.column_dimensions['E'].width = 10
        ws.column_dimensions['F'].width = 10
        ws.column_dimensions['G'].width = 10
        ws.column_dimensions['H'].width = 25
        ws.column_dimensions['I'].width = 15
        
        wb.save(output_file)
    
    def create_individual_species_sheets(self, all_data, output_file):
        """Create Excel workbook with one sheet per species"""
        from openpyxl import Workbook
        from openpyxl.styles import Font, Alignment, PatternFill
        
        wb = Workbook()
        wb.remove(wb.active)  # Remove default sheet
        
        for species in sorted(all_data['Species'].unique()):
            species_data = all_data[all_data['Species'] == species].sort_values('Start')
            
            # Create sheet (max 31 chars)
            sheet_name = species.replace("_", " ")[:31]
            ws = wb.create_sheet(sheet_name)
            
            # Add species name as title (merged row)
            ws.merge_cells('A1:I1')
            title_cell = ws.cell(1, 1, species.replace("_", " "))
            title_cell.font = Font(italic=True, bold=True, size=14)
            title_cell.alignment = Alignment(horizontal='center')
            title_cell.fill = PatternFill(start_color='E8E8E8', end_color='E8E8E8', fill_type='solid')
            
            # Headers
            headers = ['Type', 'Motif_Type', 'Repeat_Count', 'Length', 'Start', 'End', 'Region', 'Location', 'Location_Type']
            for col, header in enumerate(headers, 1):
                cell = ws.cell(3, col, header)
                cell.font = Font(bold=True)
                cell.alignment = Alignment(horizontal='center')
            
            # Data
            row = 4
            for idx, data_row in species_data.iterrows():
                location_display = self.format_location_for_display(
                    data_row['Location'], data_row['Location_Type']
                )
                
                ws.cell(row, 1, data_row['Type'])
                ws.cell(row, 2, data_row['Motif_Type'])
                ws.cell(row, 3, data_row['Repeat_Count'])
                ws.cell(row, 4, data_row['Length'])
                ws.cell(row, 5, data_row['Start'])
                ws.cell(row, 6, data_row['End'])
                ws.cell(row, 7, data_row['Region'])
                
                location_cell = ws.cell(row, 8, location_display)
                if data_row['Location_Type'] in ['Gene', 'Intron', 'tRNA', 'rRNA', 'IGS']:
                    location_cell.font = Font(italic=True)
                
                ws.cell(row, 9, data_row['Location_Type'])
                
                row += 1
            
            # Adjust column widths
            ws.column_dimensions['A'].width = 10
            ws.column_dimensions['B'].width = 12
            ws.column_dimensions['C'].width = 14
            ws.column_dimensions['D'].width = 10
            ws.column_dimensions['E'].width = 10
            ws.column_dimensions['F'].width = 10
            ws.column_dimensions['G'].width = 10
            ws.column_dimensions['H'].width = 25
            ws.column_dimensions['I'].width = 15
        
        wb.save(output_file)
    
    def create_summary_tables(self, all_data):
        """Create summary tables for all species"""
        from openpyxl import Workbook
        from openpyxl.styles import Font, Alignment
        from openpyxl.utils import get_column_letter
        
        summary_data = []
        
        for species in all_data['Species'].unique():
            species_data = all_data[all_data['Species'] == species]
            
            region_counts = species_data['Region'].value_counts()
            motif_counts = species_data['Motif_Type'].value_counts()
            loc_counts = species_data['Location_Type'].value_counts()
            
            summary = {
                'Species': species.replace("_", " "),
                'Total_SSRs': len(species_data),
                'LSC': region_counts.get('LSC', 0),
                'SSC': region_counts.get('SSC', 0),
                'IR': region_counts.get('IR', 0),
                'Mono': motif_counts.get('Mono', 0),
                'Di': motif_counts.get('Di', 0),
                'Tri': motif_counts.get('Tri', 0),
                'Tetra': motif_counts.get('Tetra', 0),
                'Penta': motif_counts.get('Penta', 0),
                'Hexa': motif_counts.get('Hexa', 0),
                'Gene': loc_counts.get('Gene', 0),
                'tRNA': loc_counts.get('tRNA', 0),
                'rRNA': loc_counts.get('rRNA', 0),
                'Intron': loc_counts.get('Intron', 0),
                'IGS': loc_counts.get('IGS', 0)
            }
            summary_data.append(summary)
        
        summary_df = pd.DataFrame(summary_data)
        
        # Create formatted Excel with italic species names
        wb = Workbook()
        ws = wb.active
        ws.title = "Summary"
        
        # Write headers
        headers = list(summary_df.columns)
        for col, header in enumerate(headers, 1):
            cell = ws.cell(1, col, header)
            cell.font = Font(bold=True)
            cell.alignment = Alignment(horizontal='center')
        
        # Write data with italic species names
        for row_idx, (idx, row_data) in enumerate(summary_df.iterrows(), 2):
            for col_idx, value in enumerate(row_data, 1):
                cell = ws.cell(row_idx, col_idx, value)
                # Italicize species column
                if col_idx == 1:
                    cell.font = Font(italic=True)
        
        # Adjust column widths
        ws.column_dimensions['A'].width = 30
        for col in range(2, len(headers) + 1):
            ws.column_dimensions[get_column_letter(col)].width = 12
        
        return wb
    
    def run_analysis(self):
        """Main analysis pipeline"""
        gb_files = [
            os.path.join(self.gb_folder, f)
            for f in os.listdir(self.gb_folder)
            if f.endswith('.gb') or f.endswith('.gbk')
        ]
        
        if not gb_files:
            print("No GenBank files found!")
            return
        
        all_ssr_data = []
        
        for gb_file in sorted(gb_files):
            df = self.process_genbank_file(gb_file)
            if df is not None and not df.empty:
                all_ssr_data.append(df)
        
        if not all_ssr_data:
            print("No SSRs detected in any files!")
            return
        
        combined_df = pd.concat(all_ssr_data, ignore_index=True)
        
        # Create formatted Excel with merged species cells
        detailed_file = os.path.join(self.output_folder, "all_ssrs_detailed.xlsx")
        print(f"Creating formatted Excel file...")
        self.create_formatted_excel(combined_df, detailed_file)
        print(f"✅ Detailed SSR data saved to: {detailed_file}")
        
        # Create summary table
        summary_file = os.path.join(self.output_folder, "ssr_summary.xlsx")
        summary_wb = self.create_summary_tables(combined_df)
        summary_wb.save(summary_file)
        print(f"✅ Summary table saved to: {summary_file}")
        
        # Create individual species sheets
        species_file = os.path.join(self.output_folder, "ssr_by_species.xlsx")
        self.create_individual_species_sheets(combined_df, species_file)
        print(f"✅ Individual species data saved to: {species_file}")
        
        print(f"\nAnalysis complete! Results in: {self.output_folder}/")
        print(f"Total SSRs detected: {len(combined_df)}")
        print(f"Species analyzed: {len(combined_df['Species'].unique())}")


def main(gb_folder=None, output_folder="ssr_analysis_results"):
    """
    Main function that can be called from command line or imported
    
    Args:
        gb_folder: Path to folder containing GenBank files (default: current directory)
        output_folder: Path to output folder (default: ssr_analysis_results)
    """
    import argparse
    
    # Check if we're in Jupyter/IPython
    try:
        get_ipython()
        in_jupyter = True
    except NameError:
        in_jupyter = False
    
    # If called from command line (not Jupyter)
    if not in_jupyter and gb_folder is None and len(sys.argv) >= 2:
        parser = argparse.ArgumentParser(
            description='Chloroplast SSR Analyzer - Detect and classify SSRs in chloroplast genomes',
            formatter_class=argparse.RawDescriptionHelpFormatter,
            epilog='''
Examples:
  # Basic usage
  python chloroplast_ssr_analyzer.py /path/to/genbank/
  
  # Custom output folder
  python chloroplast_ssr_analyzer.py /path/to/genbank/ /path/to/output/
  
  # Custom thresholds (format: mono,di,tri,tetra,penta,hexa)
  python chloroplast_ssr_analyzer.py /path/to/genbank/ -t 12,6,5,4,4,4
  
  # Less stringent thresholds
  python chloroplast_ssr_analyzer.py /path/to/genbank/ -t 8,4,3,3,3,3
            '''
        )
        parser.add_argument('genbank_folder', help='Path to folder containing GenBank (.gb/.gbk) files')
        parser.add_argument('output_folder', nargs='?', default='ssr_analysis_results',
                          help='Path to output folder (default: ssr_analysis_results)')
        parser.add_argument('-t', '--thresholds', type=str, default='10,5,4,3,3,3',
                          help='SSR detection thresholds as comma-separated values: mono,di,tri,tetra,penta,hexa (default: 10,5,4,3,3,3)')
        
        args = parser.parse_args()
        gb_folder = args.genbank_folder
        output_folder = args.output_folder
        
        # Parse thresholds
        try:
            threshold_values = [int(x.strip()) for x in args.thresholds.split(',')]
            if len(threshold_values) != 6:
                print("ERROR: Thresholds must have exactly 6 values (mono,di,tri,tetra,penta,hexa)")
                return None
            custom_thresholds = {i+1: threshold_values[i] for i in range(6)}
        except ValueError:
            print("ERROR: Invalid threshold format. Use comma-separated integers (e.g., 10,5,4,3,3,3)")
            return None
    else:
        custom_thresholds = None
    
    # Default to current directory if no folder specified
    if gb_folder is None:
        gb_folder = "."
        print("No folder specified, using current directory")
    
    if not os.path.isdir(gb_folder):
        print(f"ERROR: Directory '{gb_folder}' does not exist!")
        return None
    
    analyzer = ChloroplastSSRAnalyzer(gb_folder, output_folder)
    
    # Apply custom thresholds if provided
    if custom_thresholds:
        analyzer.thresholds = custom_thresholds
        print(f"Using custom thresholds: {custom_thresholds}")
    
    analyzer.run_analysis()
    return analyzer


# ============================================================================
# PIPELINE ENTRY POINT (USED BY UNIFIED ANALYZER)
# ============================================================================

def run_module8(gb_folder='.',
                output_folder=None,
                thresholds=None):
    """
    Unified pipeline wrapper for Module 8: SSR Analysis

    Parameters
    ----------
    gb_folder : str
        Folder containing GenBank files
    output_folder : str or None
        Output directory (None = auto-named)
    thresholds : dict or None
        SSR thresholds {1:mono, 2:di, ..., 6:hexa}
    """

    # Auto-create module-specific output folder
    if output_folder is None:
        output_folder = "Module8_SSR_Analysis"

    analyzer = ChloroplastSSRAnalyzer(
        gb_folder=gb_folder,
        output_folder=output_folder
    )

    if thresholds is not None:
        analyzer.thresholds = thresholds

    analyzer.run_analysis()
    return analyzer
```
