## Supplementary Date File 1 for "CGAS (Chloroplast Genome Analysis Suite): An Automated Python Pipeline for Comprehensive Comparative Chloroplast Genomics": module9_diversity_analysis.py

```
#!/usr/bin/env python3
"""
GenBank Gene and Intergenic Region Analysis Pipeline
Extracts genes, introns, and intergenic spacers from GenBank files, aligns them with MAFFT,
and calculates nucleotide diversity for each region.
"""

import os
import sys
import subprocess
from pathlib import Path
from Bio import SeqIO, AlignIO
from Bio.Seq import Seq
from Bio.SeqRecord import SeqRecord
from collections import defaultdict
import numpy as np
import re

def normalize_gene_name(gene_name):
    """
    Normalize gene names to handle different naming conventions:
    - tRNA: trnN-GUU, trnN_GUU, trnN_guu, trnN(GUU) -> trnK-UUU (uppercase)
    - rRNA: normalize case
    - Gene synonyms: ycf3/pafI/paf1, ycf4/pafII, psbN/psb1, psbZ/lhbA, clpP/clpP1/clp1, etc.
    """
    if not gene_name:
        return gene_name
    
    # Gene synonym mapping - maps alternative names to the primary/standard name
    gene_synonyms = {
        # ycf genes
        'pafi': 'ycf3', 'paf1': 'ycf3', 'pafI': 'ycf3',
        'pafii': 'ycf4', 'paf2': 'ycf4', 'pafII': 'ycf4',
        
        # psb genes
        'psb1': 'psbN', 'psbTn': 'psbN',
        'lhba': 'psbZ', 'lhbA': 'psbZ',
        
        # clp genes
        'clp1': 'clpP', 'clpp1': 'clpP', 'clpP1': 'clpP',
        
        # rpo genes
        'rpoa': 'rpoA', 'rpob': 'rpoB', 'rpoc1': 'rpoC1', 'rpoc2': 'rpoC2',
        
        # rpl genes (ribosomal proteins large subunit)
        'rpl2': 'rpl2', 'rpl14': 'rpl14', 'rpl16': 'rpl16', 'rpl20': 'rpl20',
        'rpl22': 'rpl22', 'rpl23': 'rpl23', 'rpl32': 'rpl32', 'rpl33': 'rpl33',
        'rpl36': 'rpl36',
        
        # rps genes (ribosomal proteins small subunit)
        'rps2': 'rps2', 'rps3': 'rps3', 'rps4': 'rps4', 'rps7': 'rps7',
        'rps8': 'rps8', 'rps11': 'rps11', 'rps12': 'rps12', 'rps14': 'rps14',
        'rps15': 'rps15', 'rps16': 'rps16', 'rps18': 'rps18', 'rps19': 'rps19',
        
        # inf gene
        'infa': 'infA',
        
        # mat gene
        'matk': 'matK',
        
        # cem gene
        'cema': 'cemA',
        
        # acc gene
        'accd': 'accD',
        
        # ndh genes
        'ndha': 'ndhA', 'ndhb': 'ndhB', 'ndhc': 'ndhC', 'ndhd': 'ndhD',
        'ndhe': 'ndhE', 'ndhf': 'ndhF', 'ndhg': 'ndhG', 'ndhh': 'ndhH',
        'ndhi': 'ndhI', 'ndhj': 'ndhJ', 'ndhk': 'ndhK',
        
        # pet genes
        'peta': 'petA', 'petb': 'petB', 'petd': 'petD', 'petg': 'petG',
        'petl': 'petL', 'petn': 'petN',
        
        # psb genes (photosystem II)
        'psba': 'psbA', 'psbb': 'psbB', 'psbc': 'psbC', 'psbd': 'psbD',
        'psbe': 'psbE', 'psbf': 'psbF', 'psbh': 'psbH', 'psbi': 'psbI',
        'psbj': 'psbJ', 'psbk': 'psbK', 'psbl': 'psbL', 'psbm': 'psbM',
        'psbt': 'psbT', 'psbz': 'psbZ',
        
        # psa genes (photosystem I)
        'psaa': 'psaA', 'psab': 'psaB', 'psac': 'psaC', 'psai': 'psaI',
        'psaj': 'psaJ', 'psam': 'psaM',
        
        # atp genes
        'atpa': 'atpA', 'atpb': 'atpB', 'atpe': 'atpE', 'atpf': 'atpF',
        'atph': 'atpH', 'atpi': 'atpI',
        
        # rbc genes
        'rbcl': 'rbcL',
        
        # ccs gene
        'ccsa': 'ccsA',
        
        # ycf genes
        'ycf1': 'ycf1', 'ycf2': 'ycf2', 'ycf3': 'ycf3', 'ycf4': 'ycf4',
    }
    
    # Handle tRNA naming conventions for ALL tRNA genes
    if gene_name.lower().startswith('trn'):
        # Extract the amino acid and anticodon
        # Patterns: trnN-GUU, trnN_GUU, trnN(GUU), trnN_guu, trnN-guu, etc.
        match = re.match(r'(trn[A-Z])[_\-\(]?([A-Z]{3})\)?', gene_name, re.IGNORECASE)
        if match:
            amino_acid = match.group(1)  # Keep original case from match
            # Ensure proper capitalization: trnK not trnk
            amino_acid = 'trn' + amino_acid[3].upper()
            anticodon = match.group(2).upper()   # e.g., 'GUU'
            return f"{amino_acid}-{anticodon}"
        else:
            # Just trnN without anticodon (e.g., trnH, trnN)
            match = re.match(r'(trn[A-Z])', gene_name, re.IGNORECASE)
            if match:
                amino_acid = match.group(1)
                return 'trn' + amino_acid[3].upper()  # trnH not trnh
    
    # Handle rRNA naming conventions
    if gene_name.lower().startswith('rrn'):
        # Normalize to lowercase
        return gene_name.lower()
    
    # Check if gene has a synonym and use the standard name
    gene_lower = gene_name.lower()
    if gene_lower in gene_synonyms:
        return gene_synonyms[gene_lower]
    
    # For other genes, return as is (keep original case)
    return gene_name

def extract_features_from_genbank(genbank_files, output_dir):
    """
    Extract genes, introns, and intergenic regions from multiple GenBank files.
    Handles tRNA naming conventions, tracks strand direction, and normalizes names.
    
    Returns:
        genes_dict: {gene_name: [SeqRecord1, SeqRecord2, ...]}
        intron_dict: {intron_name: [SeqRecord1, SeqRecord2, ...]}
        intergenic_dict: {spacer_name: [SeqRecord1, SeqRecord2, ...]}
        ambiguous_genes: set of genes without clear identification
        gene_without_anticodon: set of tRNA genes without anticodon information
        synonym_mapping: dict showing which original names were mapped to standard names
    """
    genes_dict = defaultdict(list)
    intron_dict = defaultdict(list)
    intergenic_dict = defaultdict(list)
    ambiguous_genes = set()
    gene_without_anticodon = set()
    synonym_mapping = defaultdict(set)
    
    for gb_file in genbank_files:
        print(f"Processing {gb_file}...")
        
        for record in SeqIO.parse(gb_file, "genbank"):
            organism_id = record.id
            sequence = record.seq
            
            # Extract all relevant features
            all_features = [f for f in record.features if f.type in ["gene", "CDS", "tRNA", "rRNA"]]
            
            # Group CDS/tRNA/rRNA features by gene name
            features_by_gene = defaultdict(list)
            
            for feature in all_features:
                if feature.type not in ["CDS", "tRNA", "rRNA"]:
                    continue
                
                # Get gene name
                gene_name = None
                if "gene" in feature.qualifiers:
                    gene_name = feature.qualifiers["gene"][0]
                elif "product" in feature.qualifiers:
                    gene_name = feature.qualifiers["product"][0]
                elif "locus_tag" in feature.qualifiers:
                    gene_name = feature.qualifiers["locus_tag"][0]
                else:
                    gene_name = f"unknown_{feature.location.start}"
                    ambiguous_genes.add(gene_name)
                
                # Store original name
                original_name = gene_name
                
                # Check if tRNA without anticodon
                if gene_name.lower().startswith('trn'):
                    has_anticodon_in_name = '-' in gene_name or '_' in gene_name or '(' in gene_name
                    has_anticodon_qualifier = 'anticodon' in feature.qualifiers
                    
                    if not has_anticodon_in_name and not has_anticodon_qualifier:
                        gene_without_anticodon.add(f"{gene_name} (in {organism_id} at position {feature.location.start}-{feature.location.end})")
                
                # Normalize gene name
                normalized_name = normalize_gene_name(gene_name)
                
                # Track synonyms
                if original_name != normalized_name:
                    synonym_mapping[normalized_name].add(original_name)
                
                # Group by normalized gene name and position (to handle IR copies)
                # Use position as part of key to keep IR copies separate
                key = (normalized_name, feature.location.start)
                features_by_gene[key].append(feature)
            
            # Now process each gene group
            for (gene_name, gene_start), gene_features in features_by_gene.items():
                # Extract gene sequence from the first feature (they should all be the same gene)
                first_feature = gene_features[0]
                gene_seq = first_feature.location.extract(sequence)
                strand = first_feature.location.strand
                
                # Create unique ID based on position to distinguish IR copies
                unique_id = f"{organism_id}_pos{gene_start}"
                
                seq_record = SeqRecord(
                    gene_seq,
                    id=unique_id,
                    description=f"{organism_id}_{gene_name}"
                )
                
                genes_dict[gene_name].append(seq_record)
                
                # ========================================
                # Intron extraction logic
                # ========================================
                # Combine all exon parts from all features for this gene
                # (important for genes split across multiple CDS features)
                exon_parts = []
                
                for feature in gene_features:
                    if hasattr(feature.location, "parts"):
                        exon_parts.extend(feature.location.parts)
                    else:
                        exon_parts.append(feature.location)
                
                # Sort exons by genomic position
                exon_parts.sort(key=lambda x: int(x.start))
                
                # Genes without introns (only 1 exon)
                if len(exon_parts) < 2:
                    continue
                
                # Skip trans-spliced genes (e.g., rps12 spanning LSC to IR)
                # But allow extraction of introns WITHIN a region for rps12
                gene_span = int(exon_parts[-1].end) - int(exon_parts[0].start)
                
                # For rps12: special handling
                # It has parts in LSC and IR with huge gap - but IR portion has real intron
                if gene_name.lower() == "rps12":
                    # Don't skip entirely, but filter introns later
                    # Extract introns, but only if they're < 10,000 bp (within same region)
                    pass
                elif gene_span > 10000:
                    # For other genes, skip if span > 10,000 bp
                    print(f"    Skipping trans-spliced gene {gene_name} (span: {gene_span} bp)")
                    continue
                
                # Extract introns between exons
                for i in range(len(exon_parts) - 1):
                    intron_start = int(exon_parts[i].end)
                    intron_end = int(exon_parts[i + 1].start)
                    
                    # Skip invalid introns
                    if intron_end <= intron_start:
                        continue
                    
                    intron_length = intron_end - intron_start
                    
                    # Skip trans-spliced "introns" (e.g., gap between LSC and IR in rps12)
                    if intron_length > 10000:
                        print(f"    Skipping trans-spliced intron in {gene_name} ({intron_length} bp)")
                        continue
                    
                    intron_seq = sequence[intron_start:intron_end]
                    
                    # Handle strand orientation
                    if strand == -1:
                        intron_seq = intron_seq.reverse_complement()
                    
                    # Name the intron
                    intron_name = f"{gene_name}_intron{i+1}"
                    
                    # Create unique ID including position to distinguish IR copies
                    unique_intron_id = f"{organism_id}_pos{gene_start}_intron{i+1}"
                    
                    intron_record = SeqRecord(
                        intron_seq,
                        id=unique_intron_id,
                        description=f"{organism_id}_{intron_name}"
                    )
                    
                    intron_dict[intron_name].append(intron_record)
                    print(f"    Extracted {intron_name}: {intron_start+1}..{intron_end} ({intron_length} bp)")
            
            # Extract intergenic regions
            # Use only CDS/tRNA/rRNA features
            coding_features = [f for f in all_features if f.type in ["CDS", "tRNA", "rRNA"]]
            
            if len(coding_features) > 1:
                for i in range(len(coding_features) - 1):
                    current_feature = coding_features[i]
                    next_feature = coding_features[i + 1]
                    
                    # Skip if either feature doesn't have gene name
                    if "gene" not in current_feature.qualifiers or "gene" not in next_feature.qualifiers:
                        continue
                    
                    # Get normalized gene names
                    gene1_raw = current_feature.qualifiers.get("gene", [f"gene{i}"])[0]
                    gene2_raw = next_feature.qualifiers.get("gene", [f"gene{i+1}"])[0]
                    
                    gene1 = normalize_gene_name(gene1_raw)
                    gene2 = normalize_gene_name(gene2_raw)
                    
                    # Create spacer name: gene1-gene2
                    spacer_name = f"{gene1}-{gene2}"
                    
                    # Extract intergenic sequence
                    start = int(current_feature.location.end)
                    end = int(next_feature.location.start)
                    
                    if start < end:  # Valid intergenic region
                        intergenic_seq = sequence[start:end]
                        
                        # Only include if substantial length (> 10 bp)
                        if len(intergenic_seq) > 10:
                            # Determine orientation based on flanking genes
                            strand1 = current_feature.location.strand
                            strand2 = next_feature.location.strand
                            
                            # If both genes on minus strand, reverse complement
                            if strand1 == -1 and strand2 == -1:
                                intergenic_seq = intergenic_seq.reverse_complement()
                            
                            # Use position to create unique ID for IR copies
                            unique_id = f"{organism_id}_pos{start}"
                            
                            seq_record = SeqRecord(
                                intergenic_seq,
                                id=unique_id,
                                description=f"{organism_id}_{spacer_name}"
                            )
                            
                            intergenic_dict[spacer_name].append(seq_record)
    
    return genes_dict, intron_dict, intergenic_dict, ambiguous_genes, gene_without_anticodon, synonym_mapping

def get_feature_order(genbank_file):
    """
    Extract genomic order of features (genes, introns, intergenic spacers).
    Returns dictionaries mapping feature names to their genomic positions.
    """
    gene_positions = {}
    intron_positions = {}
    intergenic_positions = {}
    
    for record in SeqIO.parse(genbank_file, "genbank"):
        all_features = [f for f in record.features if f.type in ["CDS", "tRNA", "rRNA"]]
        all_features.sort(key=lambda x: int(x.location.start))
        
        # Get gene positions
        for feature in all_features:
            gene_name = feature.qualifiers.get("gene", [None])[0]
            if gene_name:
                normalized_name = normalize_gene_name(gene_name)
                if normalized_name not in gene_positions:
                    gene_positions[normalized_name] = int(feature.location.start)
        
        # Get intron positions
        features_by_gene = defaultdict(list)
        for feature in all_features:
            gene_name = feature.qualifiers.get("gene", [None])[0]
            if gene_name:
                normalized_name = normalize_gene_name(gene_name)
                key = (normalized_name, feature.location.start)
                features_by_gene[key].append(feature)
        
        for (gene_name, gene_start), gene_features in features_by_gene.items():
            exon_parts = []
            for feature in gene_features:
                if hasattr(feature.location, "parts"):
                    exon_parts.extend(feature.location.parts)
                else:
                    exon_parts.append(feature.location)
            
            exon_parts.sort(key=lambda x: int(x.start))
            
            if len(exon_parts) >= 2:
                for i in range(len(exon_parts) - 1):
                    intron_start = int(exon_parts[i].end)
                    intron_end = int(exon_parts[i + 1].start)
                    intron_length = intron_end - intron_start
                    
                    if 0 < intron_length <= 10000:
                        intron_name = f"{gene_name}_intron{i+1}"
                        if intron_name not in intron_positions:
                            intron_positions[intron_name] = intron_start
        
        # Get intergenic spacer positions
        if len(all_features) > 1:
            for i in range(len(all_features) - 1):
                gene1 = all_features[i].qualifiers.get("gene", [None])[0]
                gene2 = all_features[i + 1].qualifiers.get("gene", [None])[0]
                
                if gene1 and gene2:
                    gene1_norm = normalize_gene_name(gene1)
                    gene2_norm = normalize_gene_name(gene2)
                    spacer_name = f"{gene1_norm}-{gene2_norm}"
                    
                    start = int(all_features[i].location.end)
                    end = int(all_features[i + 1].location.start)
                    
                    if start < end and (end - start) > 10:
                        if spacer_name not in intergenic_positions:
                            intergenic_positions[spacer_name] = start
        
        break  # Only need first record to get order
    
    return gene_positions, intron_positions, intergenic_positions

def write_fasta_files(features_dict, output_dir, prefix):
    """Write sequences to FASTA files for each feature."""
    fasta_files = []
    
    for feature_name, seq_records in features_dict.items():
        if len(seq_records) < 2:
            print(f"Skipping {feature_name}: only {len(seq_records)} sequence(s)")
            continue
        
        # Sanitize filename
        safe_name = feature_name.replace("/", "_").replace(" ", "_")
        fasta_file = os.path.join(output_dir, f"{prefix}_{safe_name}.fasta")
        
        SeqIO.write(seq_records, fasta_file, "fasta")
        fasta_files.append((feature_name, fasta_file))
    
    return fasta_files

def align_with_mafft(fasta_file, output_file):
    """Align sequences using MAFFT."""
    try:
        cmd = ["mafft", "--auto", "--quiet", fasta_file]
        with open(output_file, "w") as out:
            subprocess.run(cmd, stdout=out, stderr=subprocess.DEVNULL, check=True)
        return True
    except subprocess.CalledProcessError:
        print(f"Error aligning {fasta_file}")
        return False
    except FileNotFoundError:
        print("MAFFT not found. Please install MAFFT.")
        return False

def calculate_nucleotide_diversity(alignment_file):
    """
    Calculate nucleotide diversity (π) from aligned sequences.
    π = average number of nucleotide differences per site between any two sequences.
    """
    try:
        alignment = AlignIO.read(alignment_file, "fasta")
    except:
        return None, None, None
    
    n_sequences = len(alignment)
    alignment_length = alignment.get_alignment_length()
    
    if n_sequences < 2:
        return 0.0, 0, alignment_length
    
    # Calculate pairwise differences
    total_differences = 0
    total_comparisons = 0
    valid_sites = 0
    
    # Check each site
    for i in range(alignment_length):
        column = alignment[:, i]
        
        # Skip sites with gaps
        if '-' in column or 'N' in column.upper():
            continue
        
        valid_sites += 1
        
        # Count pairwise differences at this site
        for j in range(n_sequences):
            for k in range(j + 1, n_sequences):
                if column[j].upper() != column[k].upper():
                    total_differences += 1
                total_comparisons += 1
    
    # Calculate diversity
    if total_comparisons == 0 or valid_sites == 0:
        return 0.0, 0, alignment_length
    
    # π = average pairwise differences per site
    pi = total_differences / total_comparisons
    
    return pi, valid_sites, alignment_length

def main():
    # Configuration
    if len(sys.argv) < 2:
        print("Usage: python genbank_diversity_analysis.py <genbank_file1> <genbank_file2> ...")
        print("       python genbank_diversity_analysis.py <directory>")
        print("Example: python genbank_diversity_analysis.py *.gb")
        print("Example: python genbank_diversity_analysis.py /path/to/genbank_folder/")
        sys.exit(1)
    
    input_paths = sys.argv[1:]
    genbank_files = []
    
    # Process input paths - handle both files and directories
    for path in input_paths:
        if not os.path.exists(path):
            print(f"Error: Path {path} not found")
            sys.exit(1)
        
        if os.path.isdir(path):
            # If it's a directory, find all GenBank files
            print(f"Searching for GenBank files in directory: {path}")
            import glob
            for ext in ['*.gb', '*.gbk', '*.genbank', '*.gbff']:
                found_files = glob.glob(os.path.join(path, ext))
                genbank_files.extend(found_files)
        else:
            # It's a file
            genbank_files.append(path)
    
    if not genbank_files:
        print("Error: No GenBank files found")
        print("Supported extensions: .gb, .gbk, .genbank, .gbff")
        sys.exit(1)
    
    print(f"Processing {len(genbank_files)} GenBank files...")
    
    # Create output directories
    output_base = "Module9_Nucleotide_Diversity"
    os.makedirs(output_base, exist_ok=True)
    
    genes_dir = os.path.join(output_base, "genes")
    intron_dir = os.path.join(output_base, "introns")
    intergenic_dir = os.path.join(output_base, "intergenic")
    alignments_dir = os.path.join(output_base, "alignments")
    
    os.makedirs(genes_dir, exist_ok=True)
    os.makedirs(intron_dir, exist_ok=True)
    os.makedirs(intergenic_dir, exist_ok=True)
    os.makedirs(alignments_dir, exist_ok=True)
    
    # Extract features
    print("\nExtracting genes, introns, and intergenic regions...")
    genes_dict, intron_dict, intergenic_dict, ambiguous_genes, genes_without_anticodon, synonym_mapping = extract_features_from_genbank(genbank_files, output_base)
    
    print(f"\nFound {len(genes_dict)} unique genes")
    print(f"Found {len(intron_dict)} unique introns")
    print(f"Found {len(intergenic_dict)} intergenic regions")
    
    # Report gene synonyms that were merged
    if synonym_mapping:
        print(f"\n✓ Merged {len(synonym_mapping)} genes with alternative names:")
        for standard_name, original_names in sorted(synonym_mapping.items()):
            if len(original_names) > 1:
                names_str = ", ".join(sorted(original_names))
                print(f"  {standard_name} ← [{names_str}]")
    
    # Report problematic genes
    if ambiguous_genes:
        print(f"\n⚠ Warning: {len(ambiguous_genes)} genes without clear identification:")
        for gene in sorted(ambiguous_genes):
            print(f"  - {gene}")
    
    if genes_without_anticodon:
        print(f"\n⚠ Warning: {len(genes_without_anticodon)} tRNA genes without anticodon information:")
        for gene in sorted(genes_without_anticodon):
            print(f"  - {gene}")
        print("  Note: These tRNAs may not align properly with tRNAs from other species")
        print("        that have anticodon information. Consider manual inspection.")
    
    # Write FASTA files
    print("\nWriting FASTA files...")
    gene_fasta_files = write_fasta_files(genes_dict, genes_dir, "gene")
    intron_fasta_files = write_fasta_files(intron_dict, intron_dir, "intron")
    intergenic_fasta_files = write_fasta_files(intergenic_dict, intergenic_dir, "spacer")
    
    # Align and calculate diversity
    results = []
    
    print("\nAligning genes and calculating diversity...")
    for gene_name, fasta_file in gene_fasta_files:
        alignment_file = os.path.join(alignments_dir, f"gene_{os.path.basename(fasta_file)}")
        
        if align_with_mafft(fasta_file, alignment_file):
            pi, valid_sites, total_length = calculate_nucleotide_diversity(alignment_file)
            if pi is not None:
                results.append(("Gene", gene_name, pi, valid_sites, total_length))
                print(f"  {gene_name}: π = {pi:.6f}")
    
    print("\nAligning introns and calculating diversity...")
    for intron_name, fasta_file in intron_fasta_files:
        alignment_file = os.path.join(alignments_dir, f"intron_{os.path.basename(fasta_file)}")
        
        if align_with_mafft(fasta_file, alignment_file):
            pi, valid_sites, total_length = calculate_nucleotide_diversity(alignment_file)
            if pi is not None:
                results.append(("Intron", intron_name, pi, valid_sites, total_length))
                print(f"  {intron_name}: π = {pi:.6f}")
    
    print("\nAligning intergenic regions and calculating diversity...")
    for spacer_name, fasta_file in intergenic_fasta_files:
        alignment_file = os.path.join(alignments_dir, f"spacer_{os.path.basename(fasta_file)}")
        
        if align_with_mafft(fasta_file, alignment_file):
            pi, valid_sites, total_length = calculate_nucleotide_diversity(alignment_file)
            if pi is not None:
                results.append(("Intergenic", spacer_name, pi, valid_sites, total_length))
                print(f"  {spacer_name}: π = {pi:.6f}")
    
    # Write results to file
    results_file = os.path.join(output_base, "nucleotide_diversity_results.txt")
    with open(results_file, "w") as f:
        f.write("Nucleotide Diversity Analysis Results\n")
        f.write("=" * 80 + "\n\n")
        f.write(f"Input files: {len(genbank_files)} GenBank files\n")
        f.write(f"Total regions analyzed: {len(results)}\n\n")
        
        # Report gene synonyms that were merged
        if synonym_mapping:
            f.write("GENE SYNONYMS MERGED:\n")
            f.write("-" * 80 + "\n")
            f.write("The following alternative gene names were normalized to standard names:\n\n")
            for standard_name, original_names in sorted(synonym_mapping.items()):
                if len(original_names) > 1:
                    names_str = ", ".join(sorted(original_names))
                    f.write(f"  {standard_name} ← [{names_str}]\n")
            f.write("\n")
        
        # Report problematic genes in the file
        if ambiguous_genes or genes_without_anticodon:
            f.write("WARNINGS:\n")
            f.write("-" * 80 + "\n")
            
            if ambiguous_genes:
                f.write(f"\nGenes without clear identification ({len(ambiguous_genes)}):\n")
                for gene in sorted(ambiguous_genes):
                    f.write(f"  - {gene}\n")
            
            if genes_without_anticodon:
                f.write(f"\ntRNA genes without anticodon information ({len(genes_without_anticodon)}):\n")
                for gene in sorted(genes_without_anticodon):
                    f.write(f"  - {gene}\n")
                f.write("\nNote: These tRNAs may not align properly across species.\n")
            
            f.write("\n")
        
        f.write("-" * 80 + "\n")
        f.write(f"{'Type':<15} {'Name':<40} {'π':<12} {'Valid Sites':<12} {'Length':<10}\n")
        f.write("-" * 80 + "\n")
        
        for region_type, name, pi, valid_sites, length in results:
            f.write(f"{region_type:<15} {name:<40} {pi:<12.6f} {valid_sites:<12} {length:<10}\n")
        
        f.write("-" * 80 + "\n\n")
        
        # Summary statistics
        gene_results = [r for r in results if r[0] == "Gene"]
        intron_results = [r for r in results if r[0] == "Intron"]
        intergenic_results = [r for r in results if r[0] == "Intergenic"]
        
        if gene_results:
            gene_pi_values = [r[2] for r in gene_results]
            f.write(f"\nGene Statistics:\n")
            f.write(f"  Number of genes: {len(gene_results)}\n")
            f.write(f"  Mean π: {np.mean(gene_pi_values):.6f}\n")
            f.write(f"  Median π: {np.median(gene_pi_values):.6f}\n")
            f.write(f"  Min π: {np.min(gene_pi_values):.6f}\n")
            f.write(f"  Max π: {np.max(gene_pi_values):.6f}\n")
        
        if intron_results:
            intron_pi_values = [r[2] for r in intron_results]
            f.write(f"\nIntron Statistics:\n")
            f.write(f"  Number of introns: {len(intron_results)}\n")
            f.write(f"  Mean π: {np.mean(intron_pi_values):.6f}\n")
            f.write(f"  Median π: {np.median(intron_pi_values):.6f}\n")
            f.write(f"  Min π: {np.min(intron_pi_values):.6f}\n")
            f.write(f"  Max π: {np.max(intron_pi_values):.6f}\n")
        
        if intergenic_results:
            intergenic_pi_values = [r[2] for r in intergenic_results]
            f.write(f"\nIntergenic Region Statistics:\n")
            f.write(f"  Number of regions: {len(intergenic_results)}\n")
            f.write(f"  Mean π: {np.mean(intergenic_pi_values):.6f}\n")
            f.write(f"  Median π: {np.median(intergenic_pi_values):.6f}\n")
            f.write(f"  Min π: {np.min(intergenic_pi_values):.6f}\n")
            f.write(f"  Max π: {np.max(intergenic_pi_values):.6f}\n")
    
    print(f"\n{'='*80}")
    print(f"Analysis complete!")
    print(f"Results written to: {results_file}")
    print(f"FASTA files in: {genes_dir}, {intron_dir}, and {intergenic_dir}")
    print(f"Alignments in: {alignments_dir}")
    
    if ambiguous_genes or genes_without_anticodon:
        print(f"\n⚠ Check the results file for warnings about ambiguous genes and tRNAs")
    
    # ========================================
    # Generate additional ordered output files
    # ========================================
    print(f"\nGenerating genomic position-ordered results...")
    
    # Get genomic positions from first GenBank file
    gene_positions, intron_positions, intergenic_positions = get_feature_order(genbank_files[0])
    
    # Create results organized by genomic position
    # File 1: Separate sections for genes and non-coding regions
    ordered_results_file = os.path.join(output_base, "nucleotide_diversity_by_position.txt")
    with open(ordered_results_file, "w") as f:
        f.write("Nucleotide Diversity Results Organized by Genomic Position\n")
        f.write("=" * 80 + "\n\n")
        
        # GENES section
        f.write("PROTEIN-CODING AND RNA GENES\n")
        f.write("-" * 80 + "\n")
        f.write(f"{'Position':<12} {'Gene':<30} {'π':<12} {'Valid Sites':<12} {'Length':<10}\n")
        f.write("-" * 80 + "\n")
        
        gene_results = [r for r in results if r[0] == "Gene"]
        gene_results_with_pos = []
        for region_type, name, pi, valid_sites, length in gene_results:
            pos = gene_positions.get(name, 999999)
            gene_results_with_pos.append((pos, region_type, name, pi, valid_sites, length))
        
        gene_results_with_pos.sort(key=lambda x: x[0])
        
        for pos, region_type, name, pi, valid_sites, length in gene_results_with_pos:
            f.write(f"{pos:<12} {name:<30} {pi:<12.6f} {valid_sites:<12} {length:<10}\n")
        
        f.write("\n\n")
        
        # NON-CODING REGIONS section (introns and intergenic)
        f.write("NON-CODING REGIONS (Introns and Intergenic Spacers)\n")
        f.write("-" * 80 + "\n")
        f.write(f"{'Position':<12} {'Type':<12} {'Name':<30} {'π':<12} {'Valid Sites':<12} {'Length':<10}\n")
        f.write("-" * 80 + "\n")
        
        noncoding_results = [r for r in results if r[0] in ["Intron", "Intergenic"]]
        noncoding_with_pos = []
        
        for region_type, name, pi, valid_sites, length in noncoding_results:
            if region_type == "Intron":
                pos = intron_positions.get(name, 999999)
            else:  # Intergenic
                pos = intergenic_positions.get(name, 999999)
            noncoding_with_pos.append((pos, region_type, name, pi, valid_sites, length))
        
        noncoding_with_pos.sort(key=lambda x: x[0])
        
        for pos, region_type, name, pi, valid_sites, length in noncoding_with_pos:
            f.write(f"{pos:<12} {region_type:<12} {name:<30} {pi:<12.6f} {valid_sites:<12} {length:<10}\n")
    
    # File 2: All results combined in genomic order
    combined_ordered_file = os.path.join(output_base, "nucleotide_diversity_all_regions_by_position.txt")
    with open(combined_ordered_file, "w") as f:
        f.write("Complete Nucleotide Diversity Results in Genomic Order\n")
        f.write("=" * 80 + "\n")
        f.write("All regions (genes, introns, intergenic spacers) ordered by genomic position\n\n")
        f.write("-" * 80 + "\n")
        f.write(f"{'Position':<12} {'Type':<12} {'Name':<30} {'π':<12} {'Valid Sites':<12} {'Length':<10}\n")
        f.write("-" * 80 + "\n")
        
        # Combine all results with positions
        all_with_pos = []
        
        for region_type, name, pi, valid_sites, length in results:
            if region_type == "Gene":
                pos = gene_positions.get(name, 999999)
            elif region_type == "Intron":
                pos = intron_positions.get(name, 999999)
            elif region_type == "Intergenic":
                pos = intergenic_positions.get(name, 999999)
            else:
                pos = 999999
            
            all_with_pos.append((pos, region_type, name, pi, valid_sites, length))
        
        # Sort by genomic position
        all_with_pos.sort(key=lambda x: x[0])
        
        for pos, region_type, name, pi, valid_sites, length in all_with_pos:
            f.write(f"{pos:<12} {region_type:<12} {name:<30} {pi:<12.6f} {valid_sites:<12} {length:<10}\n")
        
        f.write("-" * 80 + "\n")
        f.write(f"\nTotal regions: {len(all_with_pos)}\n")
    
    print(f"✓ Ordered results (coding/non-coding separate): {ordered_results_file}")
    print(f"✓ Ordered results (all combined): {combined_ordered_file}")
    
    # ========================================
    # Generate R-compatible output files for plotting
    # ========================================
    print(f"\nGenerating R-compatible data files for plotting...")
    
    # File for Panel A: Genes only (ordered by position)
    r_genes_file = os.path.join(output_base, "genes_for_R_plot.txt")
    with open(r_genes_file, "w") as f:
        f.write("Region\tValue\n")
        
        gene_results = [r for r in results if r[0] == "Gene"]
        gene_results_with_pos = []
        for region_type, name, pi, valid_sites, length in gene_results:
            pos = gene_positions.get(name, 999999)
            gene_results_with_pos.append((pos, name, pi))
        
        gene_results_with_pos.sort(key=lambda x: x[0])
        
        for pos, name, pi in gene_results_with_pos:
            f.write(f"{name}\t{pi:.5f}\n")
    
    # File for Panel B: Introns and Intergenic spacers mixed (ordered by position)
    r_noncoding_file = os.path.join(output_base, "noncoding_for_R_plot.txt")
    with open(r_noncoding_file, "w") as f:
        f.write("Region\tValue\n")
        
        noncoding_results = [r for r in results if r[0] in ["Intron", "Intergenic"]]
        noncoding_with_pos = []
        
        for region_type, name, pi, valid_sites, length in noncoding_results:
            if region_type == "Intron":
                pos = intron_positions.get(name, 999999)
                # Format intron names for display: "trnK-UUU_intron1" -> "trnK intron"
                display_name = name.replace("_intron", " intron")
            else:  # Intergenic
                pos = intergenic_positions.get(name, 999999)
                display_name = name
            
            noncoding_with_pos.append((pos, display_name, pi))
        
        noncoding_with_pos.sort(key=lambda x: x[0])
        
        for pos, display_name, pi in noncoding_with_pos:
            f.write(f"{display_name}\t{pi:.5f}\n")
    
    print(f"✓ R-compatible genes data: {r_genes_file}")
    print(f"✓ R-compatible non-coding data: {r_noncoding_file}")
    print(f"\nTo use with your R script:")
    print(f"  1. Load genes: df_A <- read_delim('{r_genes_file}', delim='\\t')")
    print(f"  2. Load non-coding: df_B <- read_delim('{r_noncoding_file}', delim='\\t')")
    
    print(f"{'='*80}")

if __name__ == "__main__":
    main()
```
